# Identification of Key Genes Potentially Related to Triple Receptor Negative Breast Cancer by Microarray Analysis

**DOI:** 10.1101/2020.12.21.423796

**Authors:** Basavaraj Vastrad, Chanabasayya Vastrad, Iranna Kotturshetti

**Author notes:** Chanabasayya Vastrad, Chanabasava Nilaya, Bharthinagar, Dharwad 580001, Karanataka, India.

## Abstract

Triple receptor negative breast cancer (TNBC) is the type of gynecological cancer in the elderly women. This study is aimed to explore molecular mechanism of TNBC via bioinformatics analysis. The gene expression profiles of GSE88715 (including 38 TNBC and 38 normal control) was downloaded from the Gene Expression Omnibus (GEO) database. Differentially expressed genes (DEGs) were screened using the limma package in R software. Pathway and gene ontology (GO) enrichment analysis were performed based on various pathway dabases and GO database. Then, InnateDb interactome database, Cytoscape and PEWCC1 were applied to construct the protein-protein interaction (PPI) network and screen hub genes. Similarly, miRNet database, NetworkAnalyst database and Cytoscape were applied to construct the target gene - miRNA network and target gene - TF network, and screen targate genes. Pathway and GO enrichment analysis was further performed for hub genes, gene clusters identified via module analysis and targate genes. The expression of hub genes with prognostic values was validated on the UALCAN, cBio Portal, The Human Protein Atlas, receiver operator characteristic (ROC) curve analysis, RT-PCR analysis and immune infiltration analysis. A total of 949 DEGs were identified in TNBC (469 up regulated genes, and 480 down regulated genes), and they were mainly enriched in the terms of phospholipases, toxoplasmosis, immune response, cell surface, glycolysis, biosynthesis of amino acids, carboxylic acid metabolic process and organic substance catabolic process extracellular space. Hub genes including UBD, HLA-B, MYC and HSP90AB1 were identified via PPI network and modules, which were mainly enriched in immune response, antigen processing and presentation, cell cycle and pathways in cancer. Targate genes including CCDC80, PEG10, HOPX and CCNA2 were identified via target gene - miRNA network and target gene - TF network, which were mainly enriched in extracellular structure organization, validated targets of C-MYC transcriptional activation, ensemble of genes encoding core extracellular matrix including ECM glycoproteins and cell cycle. The top five significantly overexpressed mRNA (ADAM15, BATF, NOTCH3, ITGAX and SDC1) and the top five significantly underexpressed mRNA (RPL4, EEF1G, RPL3, RBMX and ABCC2) were selected for further validation in TNBCpatients and healthy controls. Analysis of the expression of genes in the various databases showed that ADAM15, BATF, NOTCH3, ITGAX, SDC1, RPL4, EEF1G, RPL3, RBMX and ABCC2 expressions have a cancer specific pattern in TNBC. Collectively, ADAM15, BATF, NOTCH3, ITGAX, SDC1, RPL4, EEF1G, RPL3, RBMX and ABCC2 may be useful candidate biomarkers for TNBC diagnosis, prognosis and theraputic targates.

## Introduction

Triple receptor negative breast cancer (TNBC) is one of the most common malignant tumors in women and a major cause of cancer-related death worldwide, especially in developing countries [Boyle, 2012]. This cancer is linked with absence of estrogen receptor (ER), progesterone receptor (PR), and HER2 [Bauer et al. 2007]. The mean survival time of TNBC patients without intervention is estimated between 12 and 18 months [Kassam et al. 2009]. Although surgical resection [Puglisi, 2010], radiation therapy [Abdulkarim et al. 2011], chemotherapy [Bhola et al. 2013] and immunotherapy [Stagg and Allard, 2013] have been used to treat HCC, treatment outcomes are still unsatisfactory due to postsurgical recurrence and chemo resistance. Therefore, a further examination into the underlying molecular mechanisms of TNBC initiation and development is urgently needed, which will contribute to the discovery of new prognostic, diagnostic and therapeutic targets for TNBC.

In the past decades, there have been many reports of genes and signaling pathways associated in TNBC initiation and advancement. Genes such as Cav-1[Witkiewicz et al. 2010], Notch-1 and Notch-4 [Speiser et al. 2012], androgen receptor, EGFR, and BRCA1 [Zhang et al. 2015], p53 [Bae et al. 2018] and ZEB1 [Jang et al. 2015] were associated with pathogenesis of TNBC. Signaling pathways such as Wnt-β Catenin pathway [De et al. 2017], FGFR signaling pathway [Sharpe et al. 2011], PI3K-AKT pathway [Wu et al. 2018], insulin-like growth factor-binding protein-3 signaling pathway [Martin et al. 2014] and TDO2-AhR signaling pathway [D’Amato et al. 2015] were responsible for pathogenesis of TNBC. Therefore, it is meaningful to conduct an analysis of gene profiles to provide novel clues to the pathogenesis of TNBC.

DNA microarray is multiplex labs on a chip. This screening uses miniaturized and multiplexed corollary transform and disclosure methods. It is favored technology in current years [Schena et al. 1995] and is frequently used to obtain data regarding genetic modifications during cancer progression [Mischel et al. 2004]. This data may be important in the recognition of molecular markers. In the current study, data from mRNA microarray dataset was used to analyze the differentially expressed genes (DEGs) in TNBC (tumor stroma) compared with normal tissue (normal stroma). Bioinformatics methods, including pathway enrichment analysis and Gene Ontology (GO) enrichment analysis using toppcluster online tool, protein-protein interaction network and module analysis using InnateDB and PEWCC1, target genes - miRNA regulatory network and target genes - miRNA regulatory network using mirnet and Jasper databases, and validation of hub genes using the UALCAN, cBio Portal, The Human Protein Atlas, receiver operator characteristic (ROC) curve analysis, RT-PCR analysis and immune infiltration anaysis, were used to identify the important genes in TNBC.

## Materials and methods

### Microarray dataset and data processing

The mRNA dataset GSE88715 [Gruosso et al. 2019] was downloaded from Gene Expression Omnibus (GEO) database (www.ncbi.nlm.nih.gov/geo/) [Barrett et al. 2013]. GSE88715 was produced on the platform GPL14550 Agilent-028004 SurePrint G3 Human GE 8x60K Microarray (Probe Name Version). A total of 38 tissue samples from TNBC (tumor stroma) and 38 tissue samples from normal tissue (normal stroma) were included in the GSE88715 dataset.

Normalization for the downloaded original data was performed using limma package (version: 3.9) [Ritchie et al. 2013] in R software. The preprocessing in this investigation enclosed background correction, normalization, and expression quantification. If different probes were profiled to the same mRNA symbol, the mean value of these probes was treated as the final expression value of this mRNA.

### Identification of DEGs

Based on classical Bayes decision method, the P values of DEGs between TNBC (tumor stroma) samples and normal tissue (normal stroma) samples were calculated respectively by limma package (version: 3.9) [Ritchie et al. 2013] in R software (version: 3.6.1). Then, differentially expressed genes (DEGs) meeting the cut-off criteria of adjusted P<0.05, |log2fold-change (FC)|>2.28 were selected as the up regulated genes and |log2fold-change (FC)|>-1.947 were selected as the down regulated genes. The bidirectional hierarchical clustering and volcano plot analysis for DEGs were performed by R software (version: 3.6.1).

### Pathway enrichment analysis of DEGs

BIOCYC (https://biocyc.org/) [Caspi et al. 2016], Kyoto Encyclopedia of Genes and Genomes (KEGG) (http://www.genome.jp/kegg/pathway.html) [Kanehisa et al. 2018], Pathway Interaction Database (PID) (https://wiki.nci.nih.gov/pages/viewpage.action?pageId=315491760) [Schaefer et al. 2009], REACTOME (https://reactome.org/) [Fabregat et al. 2018], GenMAPP (http://www.genmapp.org/) [Dahlquist et al. 2002], MSigDB C2 BIOCARTA (http://software.broadinstitute.org/gsea/msigdb/collections.jsp) [Subramanian et al. 2005], PantherDB (http://www.pantherdb.org/) [Mi et al. 2017], Pathway Ontology (http://www.obofoundry.org/ontology/pw.html) [Petri et al. 2017] and Small Molecule Pathway Database (SMPDB) (http://smpdb.ca/) [Jewison et al. 2014], a comprehensive knowledge databases, plays an essential role in both functional interpretation and practical application of genomic information. The DEGs (up and down regulated genes) list was uploaded to online tool ToppCluster (https://toppcluster.cchmc.org/) [Kaimal et al. 2010] to obtain enriched significant pathway analysis with P<0.05.

### Gene ontology (GO) enrichment analysis of DEGs

Gene ontology (GO) (http://www.geneontology.org/) [Harris et al. 2004] enrichment analysis were performed using the online tool ToppCluster (https://toppcluster.cchmc.org/) [Kaimal et al. 2010]. By this way, the biological process (BP), cellular component (CC) and molecule function (MF) of target genes can be defined. The up and down regulated genes were detached. Enriched items with p value < 0.05 was considered to be statistically significant.

### PPI network construction and module analysis

InnateDb interactome database (https://www.innatedb.com/) [Breuer et al. 2013] is an online database of known and predicted protein - protein interactions (PPI). InnateDb interactome integrates various PPI databases such as IntAct Molecular Interaction Database (https://www.ebi.ac.uk/intact/) [Orchard et al. 2014], Biological General Repository for Interaction Datasets (BioGRID) (https://thebiogrid.org/) [Chatr-Aryamontri et al. 2017], the Molecular INTeraction database (MINT) (https://mint.bio.uniroma2.it/) [Licata et al. 2012], Database of Interacting Proteins (DIP) (http://dip.doe-mbi.ucla.edu/dip/Main.cgi) [Salwinski et al. 2004] and Biomolecular Interaction Network Database (BIND) (http://bond.unleashedinformatics.com/) [Willis and Hogue, 2006]. To assess the interactions among up and down regulated genes, we mapped them to InnateDb interactome database and and the coactions with a combined score of >0.4 were considered. Then, the PPI networks were visualized using Cytoscape software version 3.7.0 software (http://cytoscape.org/) [Shannon et al. 2003]. Network Analyzer plugin in Cytoscape topologically calculates the usefulness of nodes in a biological network by metrics such as node degree [Rito et al. 2010], betweenness [Yang and Lonardi, 2003], stress [Chen et al. 2014], closeness [Hsu et al. 2008] and clustering coefficient [Asur et al. 2007]. Modules of up and down regulated genes were established by the PEWCC1 [Zaki et al. 2013] with the concrete selection standards, which were as follows: PEWCC1 scores >2 and number of nodes >9.

### Construction of the target gene **-** miRNA network

To understand the associations between target gene expression and miRNAs, the target genes (up and down regulated genes) were predicted using various miRNA databases such as TarBase (http://diana.imis.athena-innovation.gr/DianaTools/index.php?r=tarbase/index) [Vlachos et al. 2015], miRTarBase (http://mirtarbase.mbc.nctu.edu.tw/php/download.php) [Chou et al. 2018], miRecords (http://miRecords.umn.edu/miRecords) [Xiao et al. 2009], miR2Disease (http://www.mir2disease.org/) [Jiang et al. 2009], HMDD (http://www.cuilab.cn/hmdd) [Huang et al. 2019], PhenomiR (http://mips.helmholtz-muenchen.de/phenomir/) [Ruepp et al. 2010], SM2miR (http://bioinfo.hrbmu.edu.cn/SM2miR/) [Liu et al. 2013], PharmacomiR (http://www.pharmaco-mir.org/) [Rukov et al. 2014], EpimiR (http://bioinfo.hrbmu.edu.cn/EpimiR/) [Dai et al. 2014] and starBase (http://starbase.sysu.edu.cn/) [Li et al. 2014]. The target gene -miRNA interactions were generated by miRNet (https://www.mirnet.ca/) [Fan and Xia, 2018] and was visualized using Cytoscape software version 3.7.0 software (http://cytoscape.org/) [Shannon et al. 2003] to further characterize correlations between potential target genes and miRNAs.

### Construction of the target gene **-** TF network

JASPAR (http://jaspar.genereg.net/) [Khan et al. 2018] is an international research consortium that uses large-scale, genome-wide assays to identify the role of all transcription factors (TFs) of the human genome. The transcription regulatory association between up and down regulated genes were explored by JASPAR, generated by NetworkAnalyst (https://www.networkanalyst.ca/) [Zhou et al. 2019] and was visualized using Cytoscape software version 3.7.0 software (http://cytoscape.org/) [Shannon et al. 2003] to further characterize correlations between potential target genes and TFs.

### Validation of hub genes

Overall survival (OS) analysis of hub genes were downloaded from TCGA database by UALCAN (http://ualcan.path.uab.edu/analysis.html) [Chandrashekar et al. 2017]. According to the median mRNA expression of each hub genes, the TNBC samples were divided into two groups: high (expression higher than median value) and low (expression lower than median value) and P<0.05 was thought to be significant. Expression analysis of hub genes in TNBC was performed (normal and TNBC patients) from TCGA database by UALCAN (http://ualcan.path.uab.edu/analysis.html) [Chandrashekar et al. 2017]. Stage analysis of hub genes in TNBC was performed (all ten stages of cancer) from TCGA database by UALCAN (http://ualcan.path.uab.edu/analysis.html) [Chandrashekar et al. 2017]. Mutation analysis of hub genes was performed from TCGA database by cBioPortal online platform (http://www.cbioportal.org) [Gao et al. 2013]. We used the immunohistochemical analysis (IHC) of hub genes from The Human Protein Atlas (https://www.proteinatlas.org) [Uhlén et al. 2015] to verify the protein expression level in TNBC. pROC package of the R software was used for receiver operator characteristic (ROC) curve analysis and were analyzed using the generalized linear model (GLM) in machine learning algorithms [Robin et al. 2011]. Area under the ROC curve (AUC) was used to evaluate the predictive accuracy of selected hub genes for TNBC. P < 0.05 was considered statistically significant. For validation of hub genes by RT-PCR first total RNA from the TNBC and normal breast tissue was extracted using TRI Reagent® (Sigma, USA) according to the manufacturer’s instructions. Total RNA was reverse-transcribed to cDNA using FastQuant RT kit (with gDNase; Tiangen Biotech Co., Ltd.). Real-time quantitative PCR was performed using QuantStudio 7 Flex real-time PCR system (Thermo Fisher Scientific, Waltham, MA, USA) with the following conditions: 50 ◦C for 2 min and 95 ◦C for 10 min, followed by 40 cycles of 95◦C for 10 s and 60◦C for 60 s. All PCR reactions were performed in triplicate. The mRNA expression levels of the tested hub genes relative to β ctin were determined using the 2^-ΔΔCt^ method and as fold induction [Livak and Schmittgen, 2001]. Primers used are listed in Table 1. The TIMER (https://cistrome.shinyapps.io/timer/) [Li et al. 2017] is a RNA-Seq expression profiling database which provides a comprehensive resource for systematic analysis of immune infiltrates across various cancer types. Immune cell types such as B cells, CD4+ T cells, CD8+ T cells, neutrophils, macrophages, and dendritic cells were assessed by TIMER on breast cancer sample data, and the correlation between 10 hub genes expression and immune infiltration.

**Table 1.**
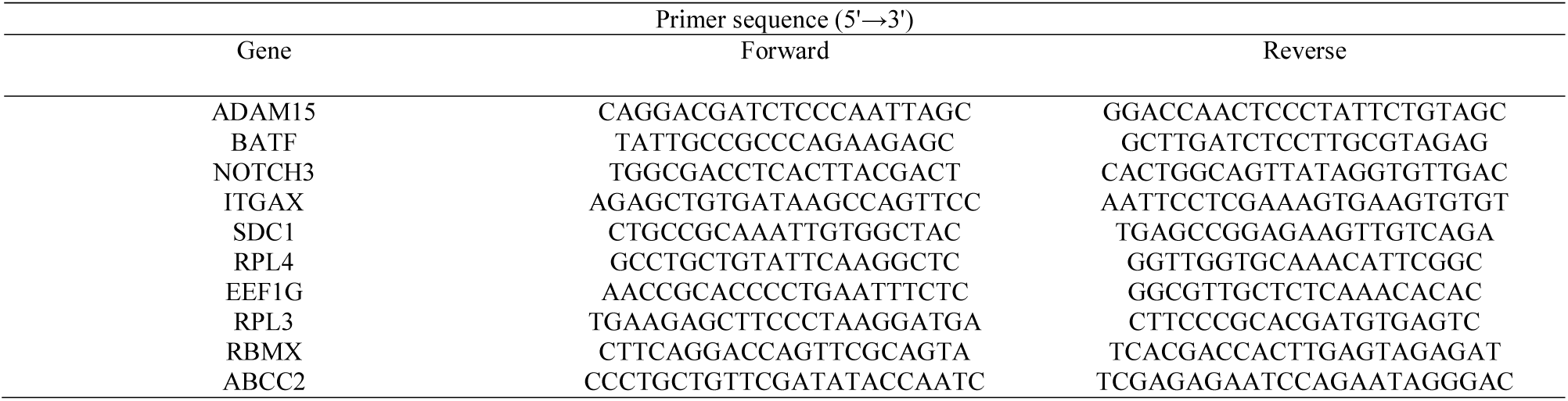
Primers used for quantitative PCR

## Results

### Identification of DEGs

The raw data of dataset was normalized following data pretreatment. Fig. 1A and Fig. 1B demonstrates the boxplot of each sample prior to and following data normalization. The study included 38 tissue samples from TNBC (tumor stroma) and 38 tissue samples from normal tissue (normal stroma). A total of 949 DEGs were identified after the analysis of GSE88715 by limma package in R software. Of these, 469 were up regulated and 480 were down regulated in TNBC (tumor stroma) samples compared with normal tissue (normal stroma) samples (Table 2). A volcano plot and heat map of DEGs is shown in Fig. 2. Heat map of up and down regulated genes are shown in Fig. 3 and Fig. 4, respectively.

**Fig. 1.**
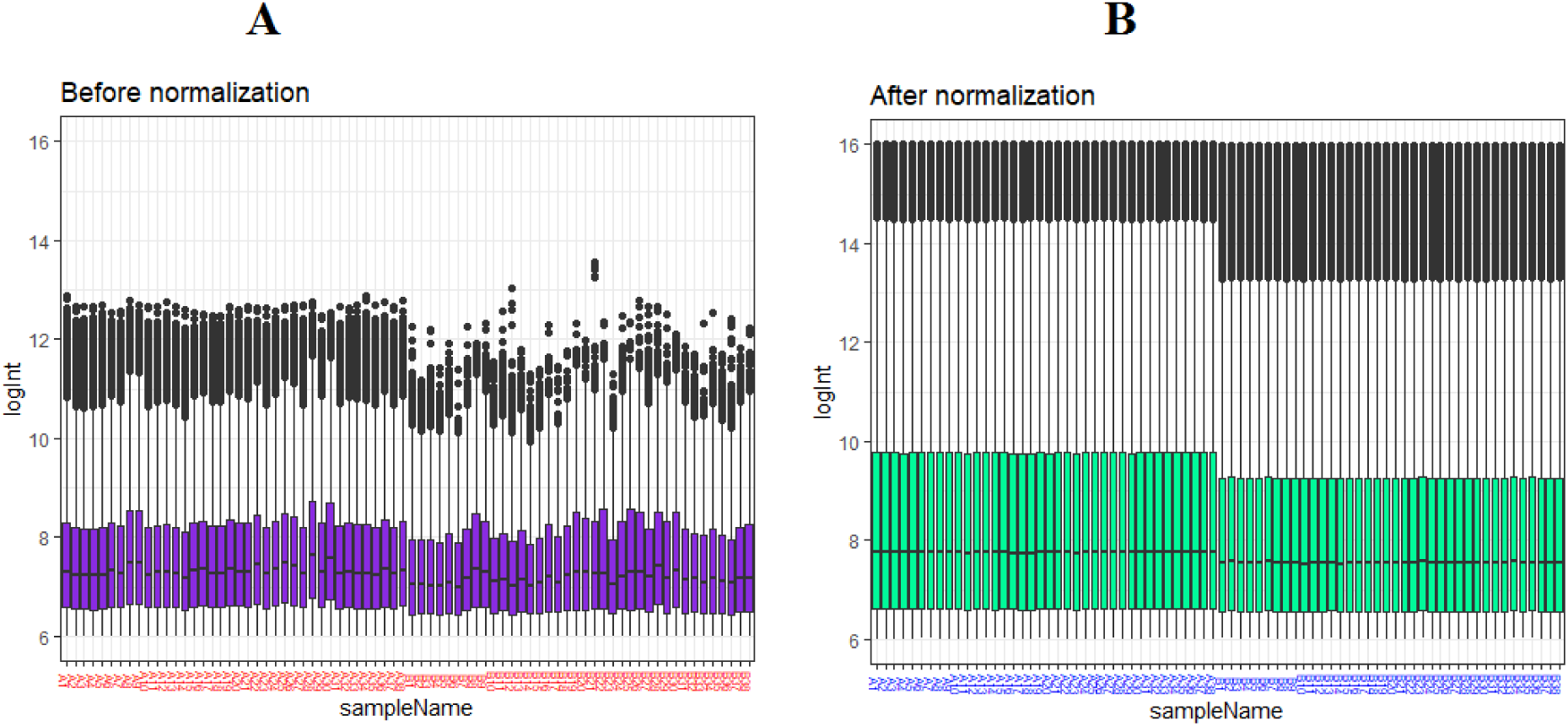
Box plots of the gene expression data before normalization (A) and after normalization (B). Horizontal axis represents the sample symbol and the vertical axis represents the gene expression values. The black line in the box plot represents the median value of gene expression. (A1 - A38 = TNBC (tumor stroma) samples; B1 - B38 = normal tissue (normal stroma) samples)

**Fig. 2.**
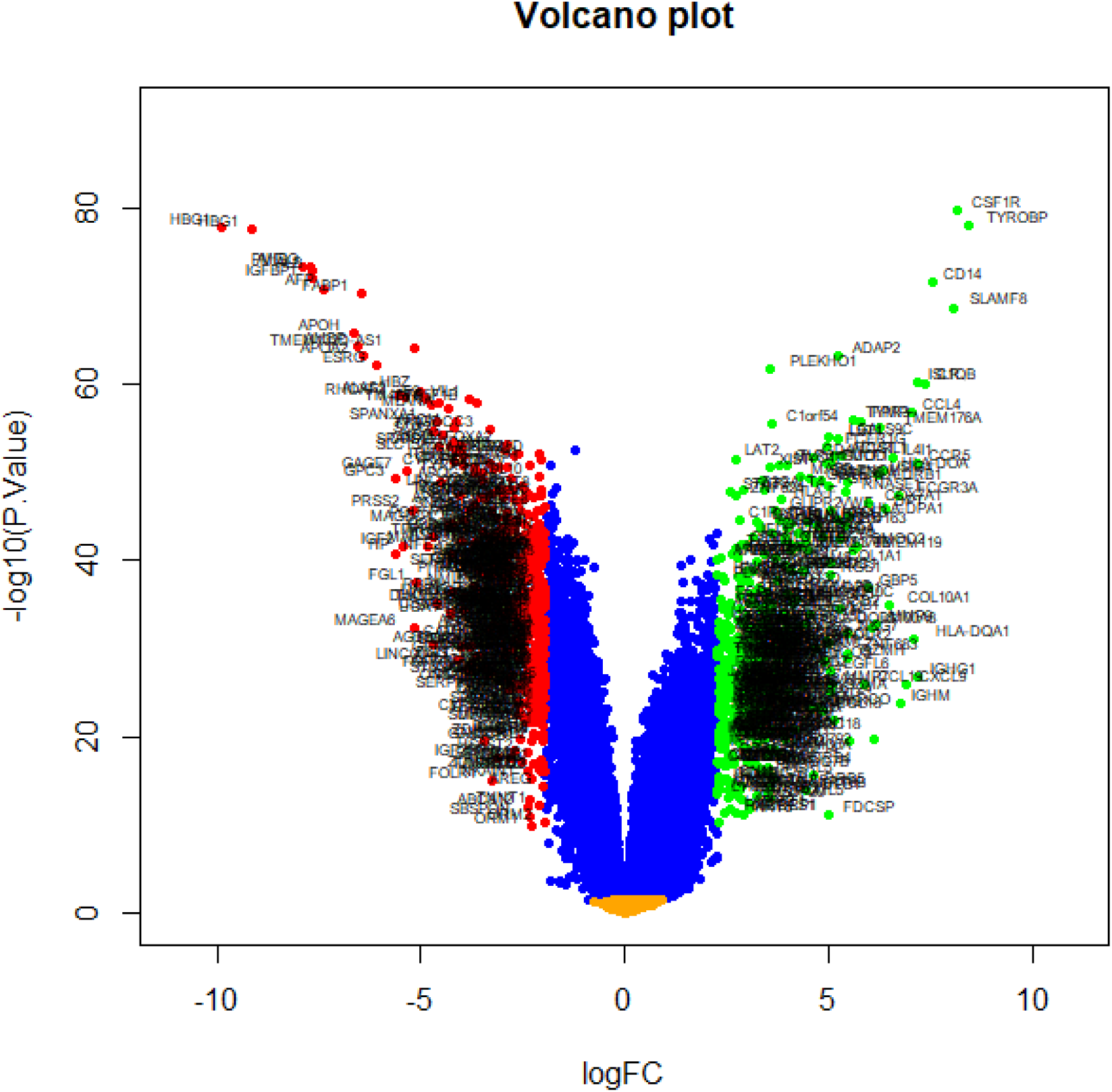
Volcano plot of differentially expressed genes. Genes with a significant change of more than two-fold were selected.

**Fig. 3.**
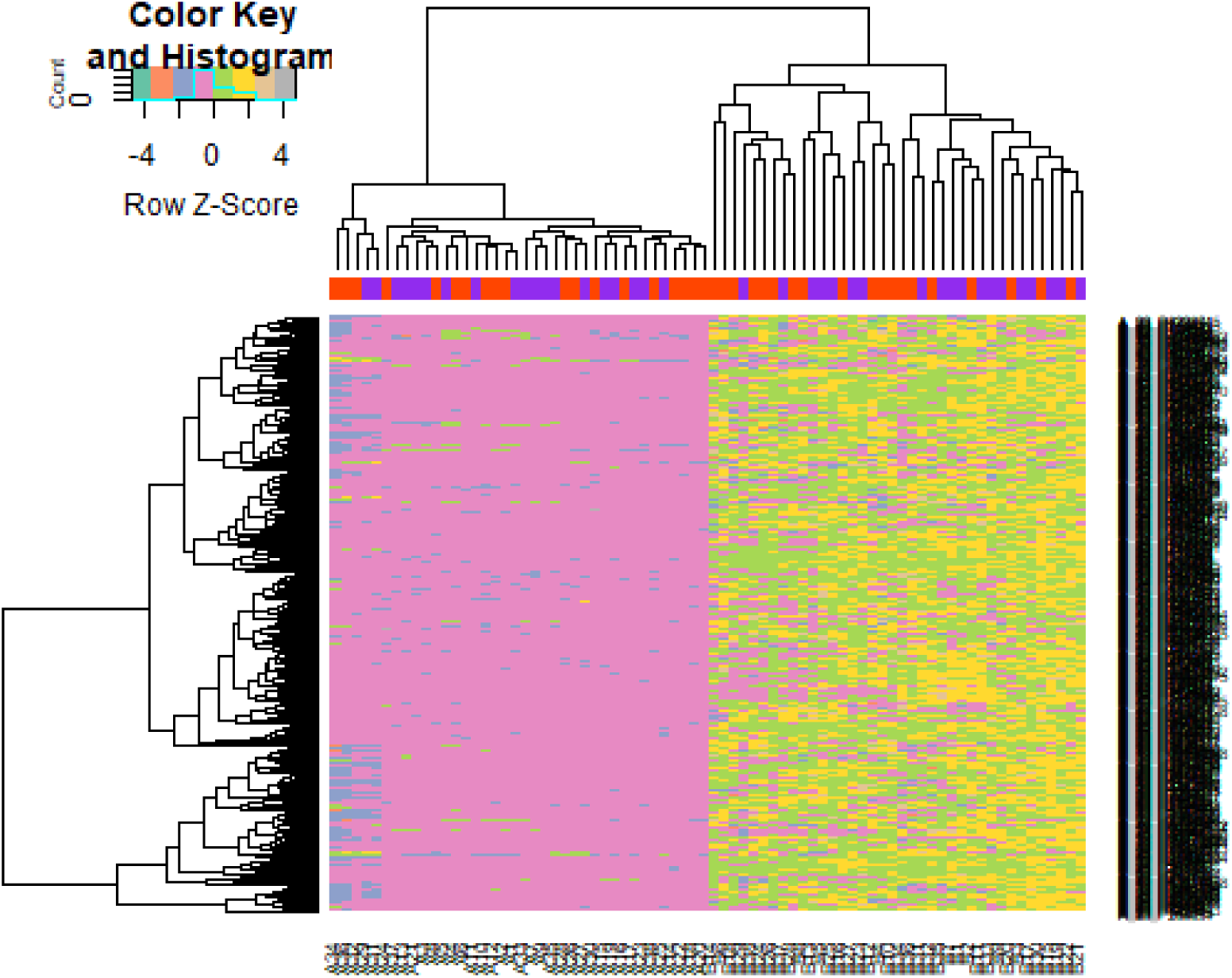
Heat map of up regulated differentially expressed genes. Legend on the top left indicate log fold change of genes. (A1 - A38 = TNBC (tumor stroma) samples; B1 - B38 = normal tissue (normal stroma) samples)

**Fig. 4.**
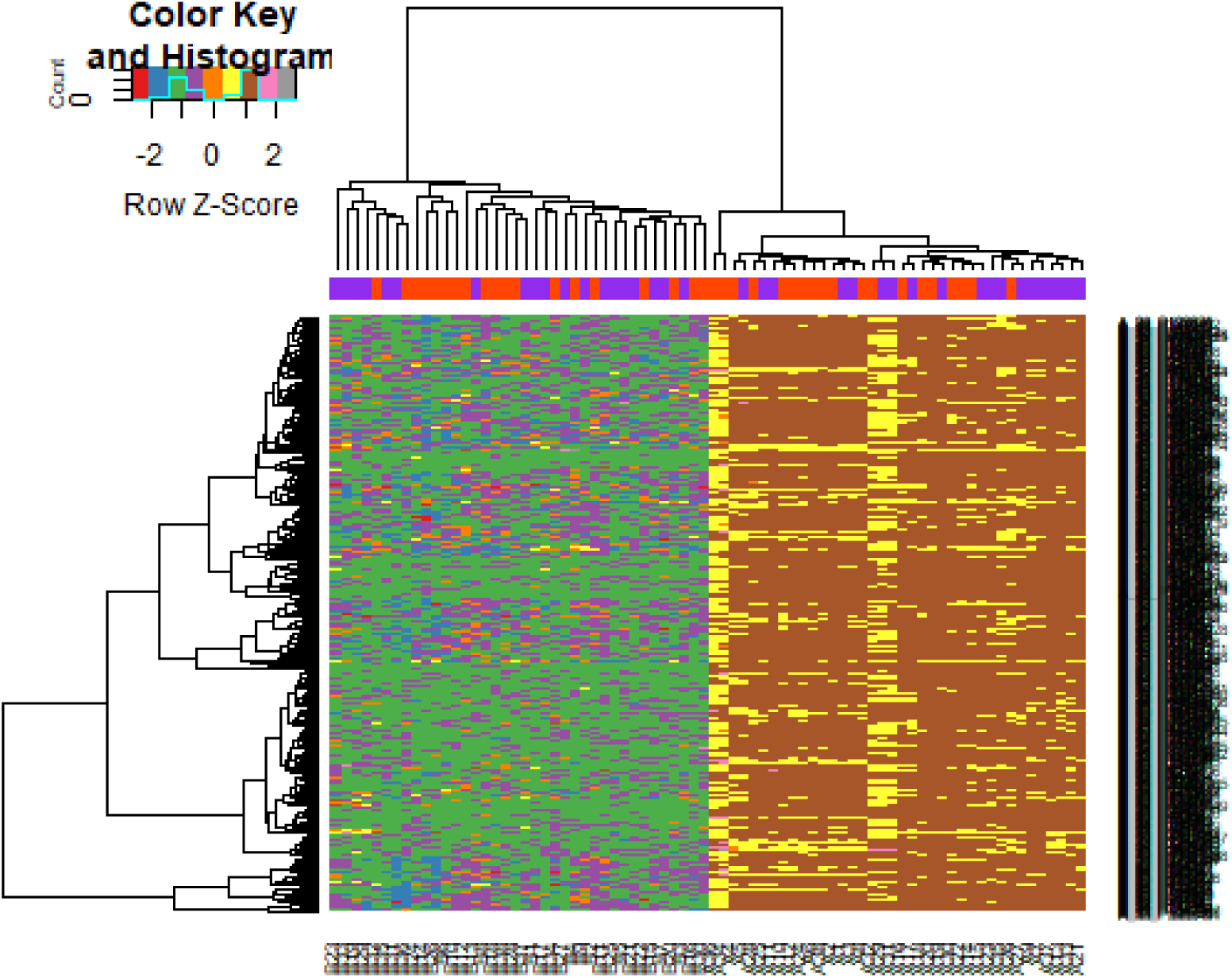
Heat map of down regulated differentially expressed genes. Legend on the top left indicate log fold change of genes. (A1 - A38 = TNBC (tumor stroma) samples; B1 - B38 = normal tissue (normal stroma) samples)

**Table 2.**
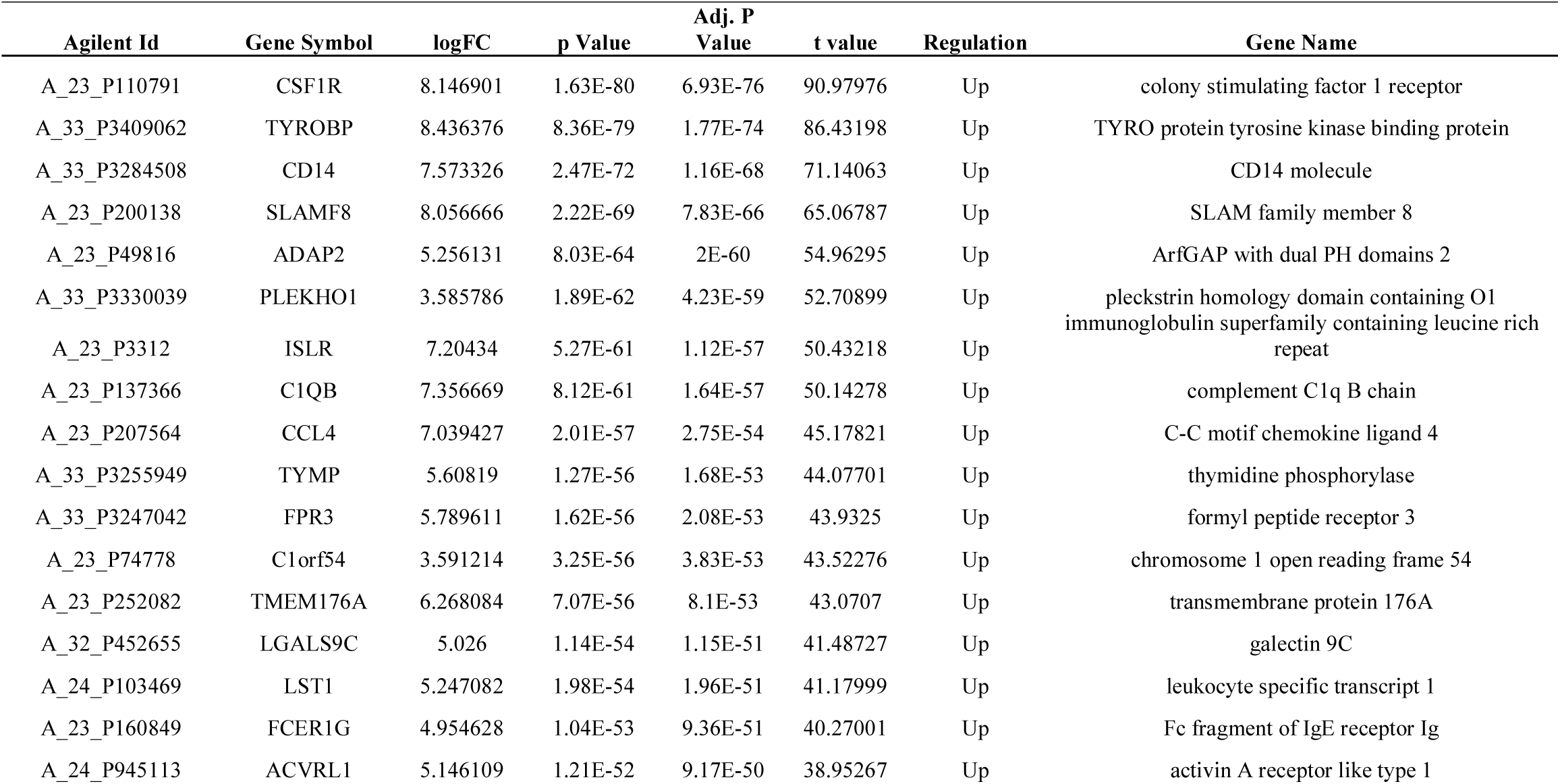
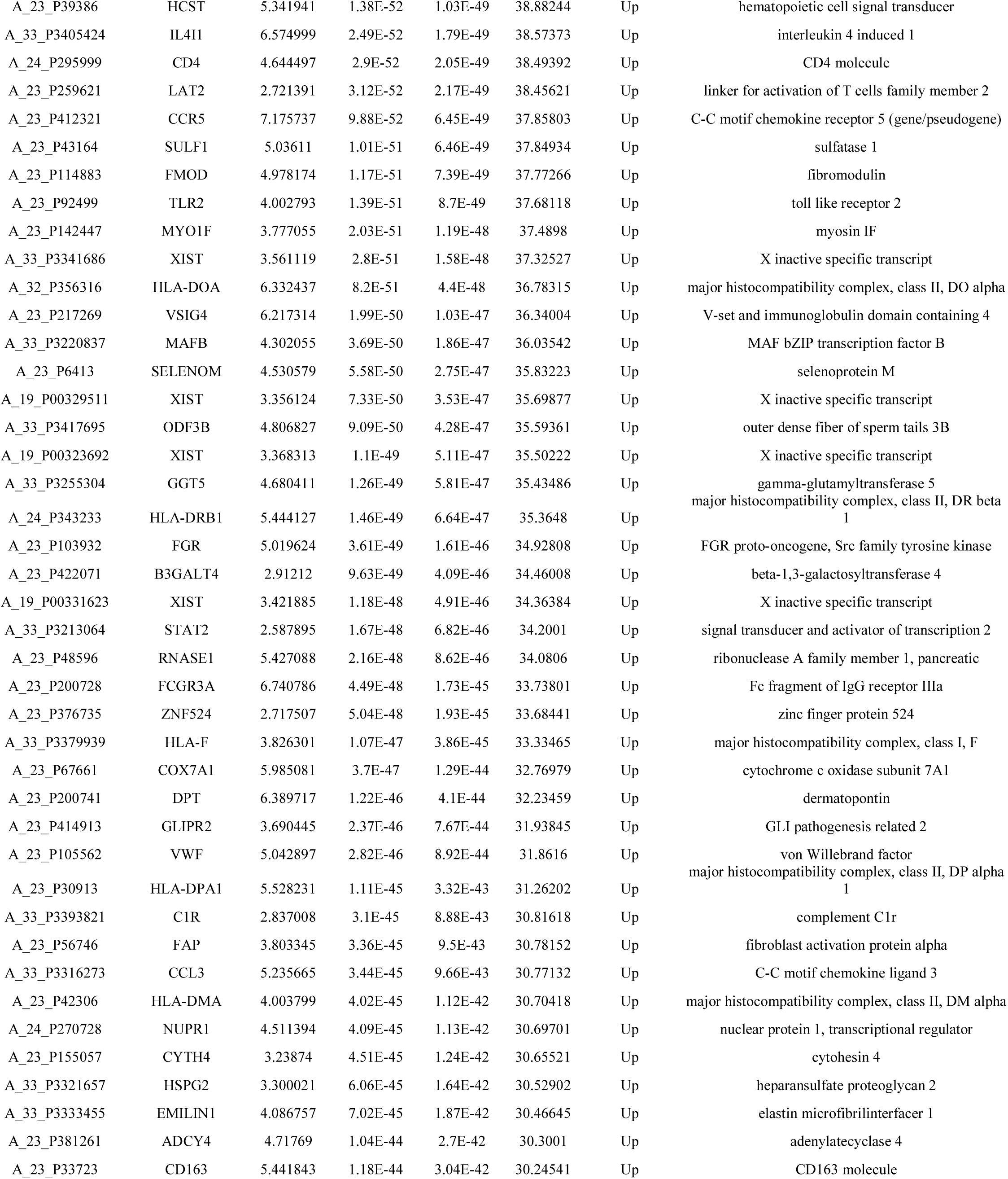
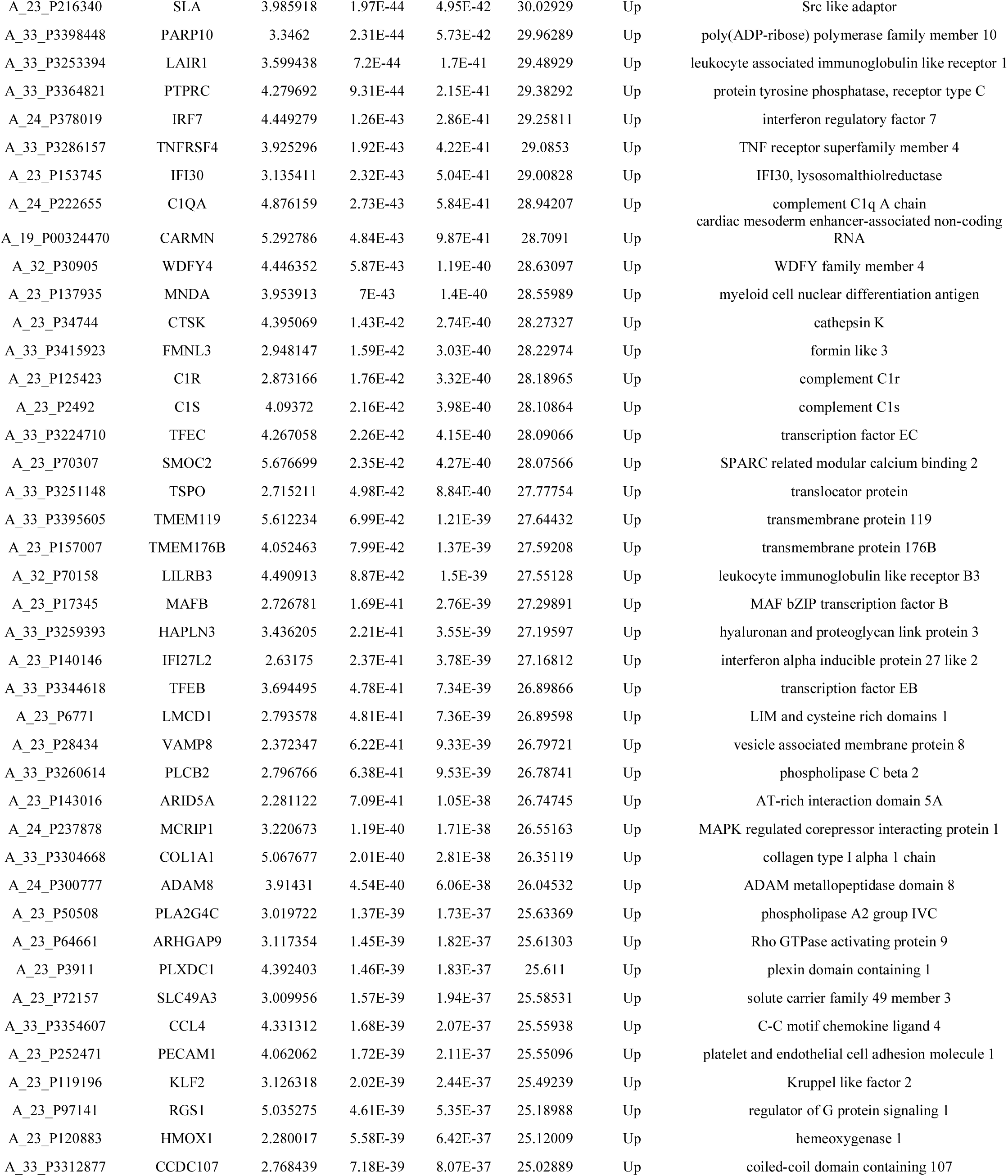
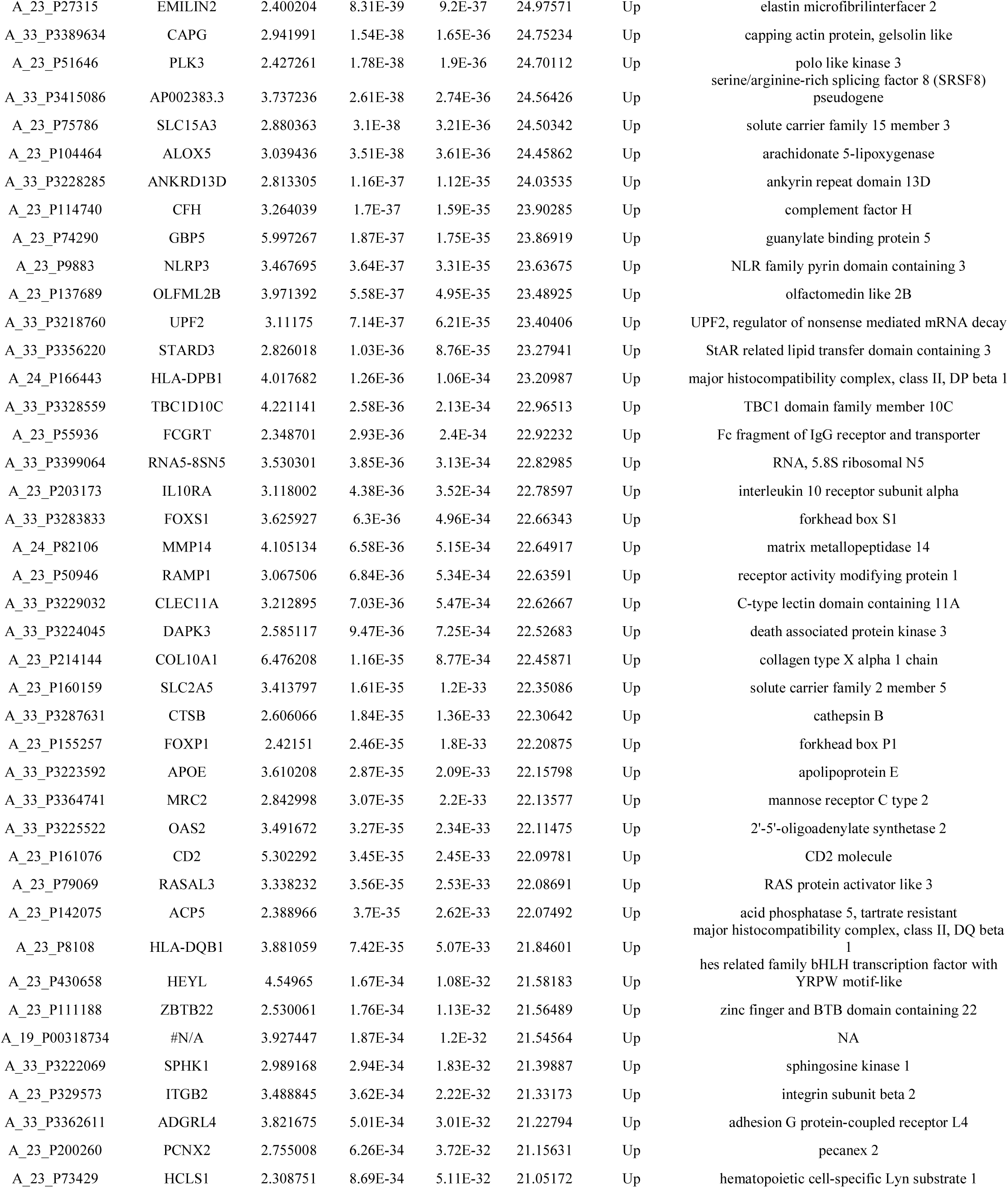
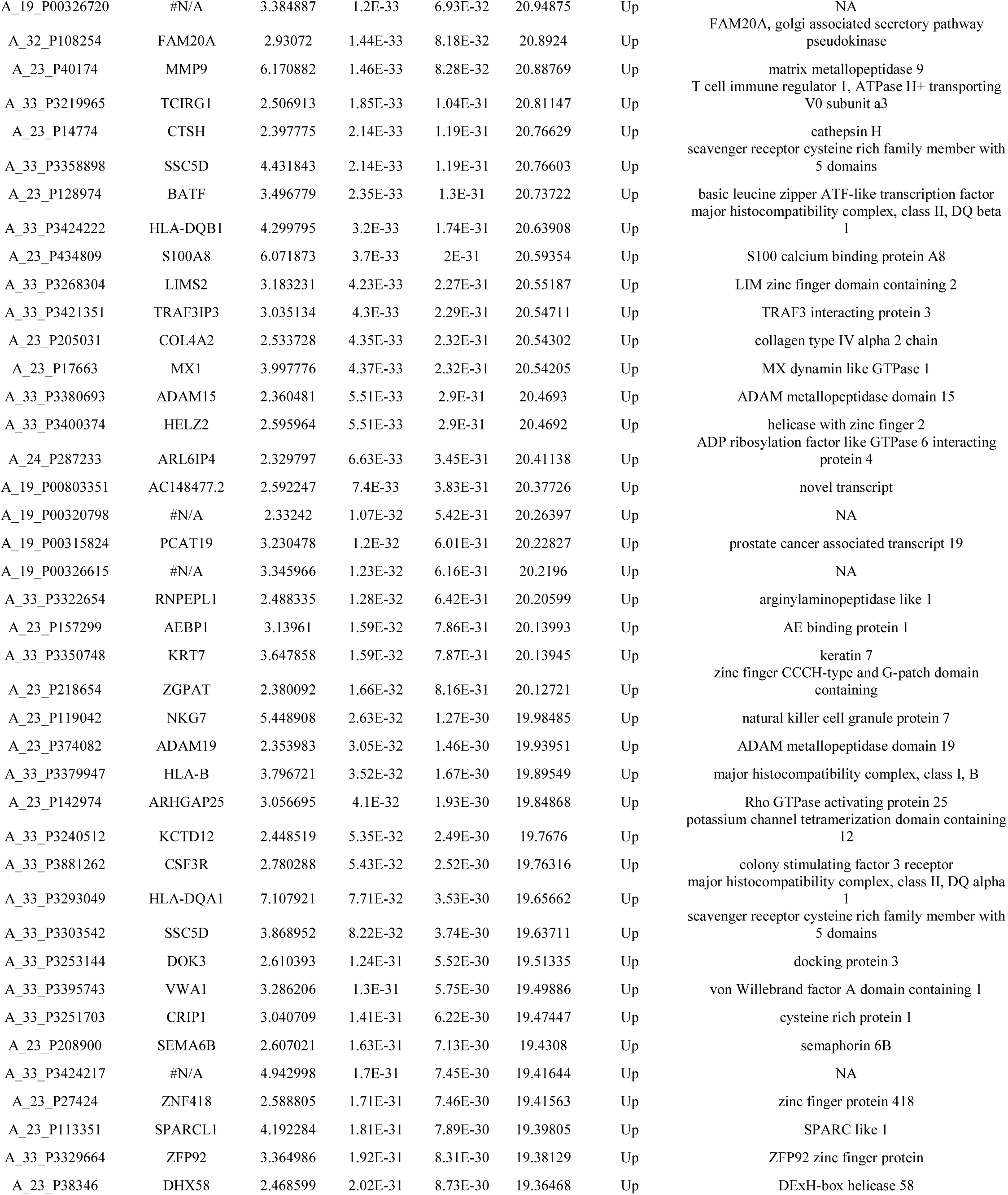
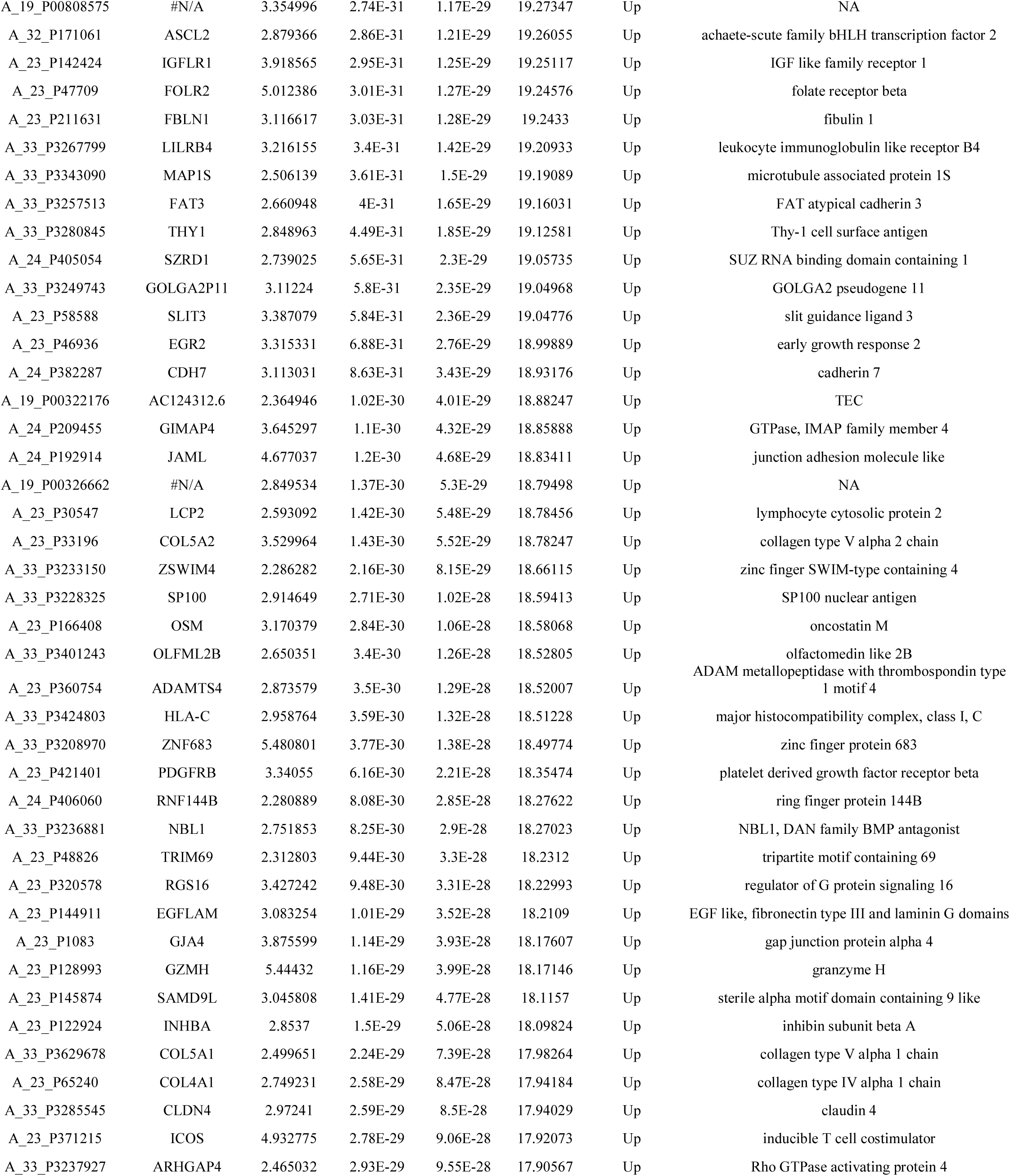
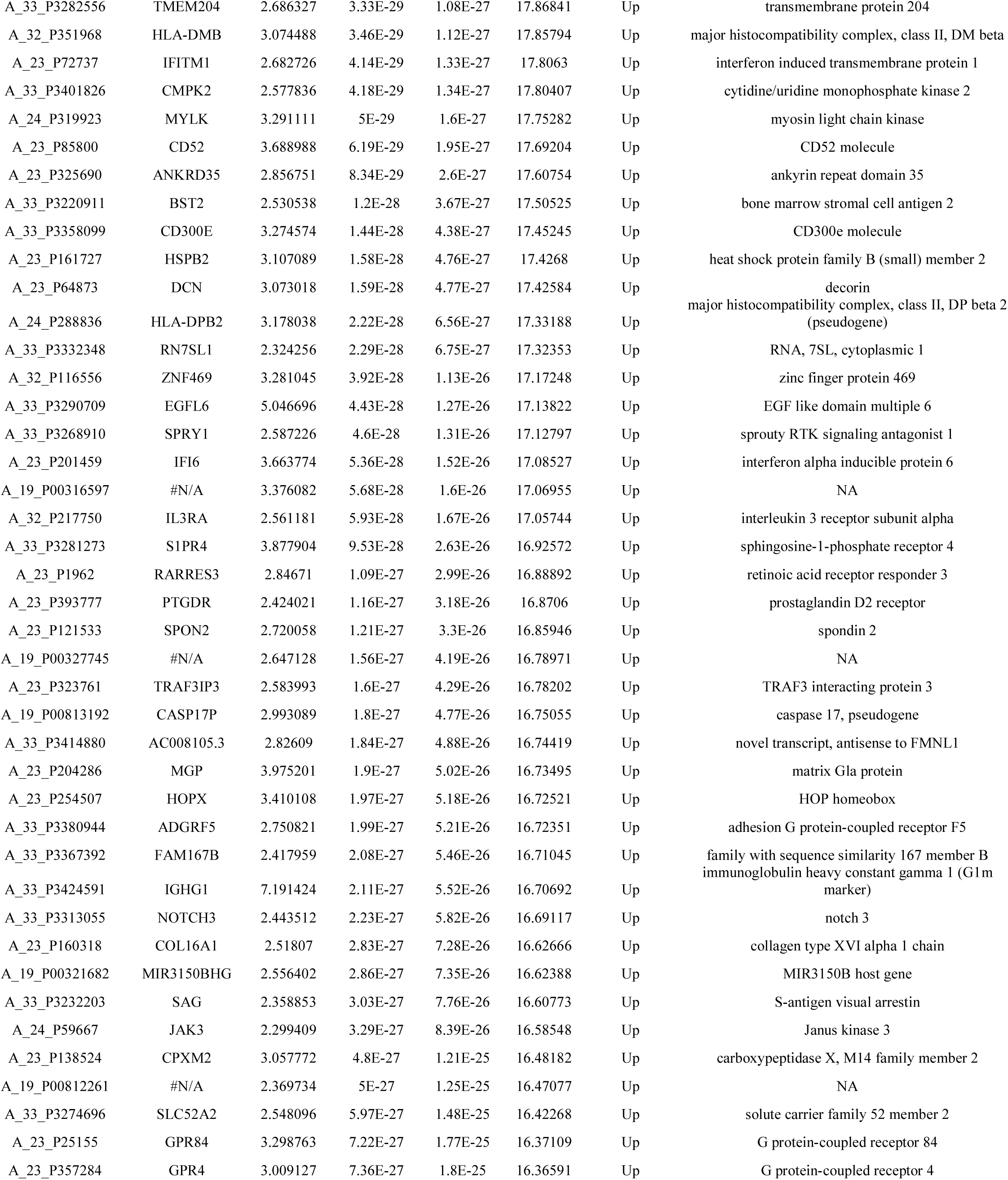
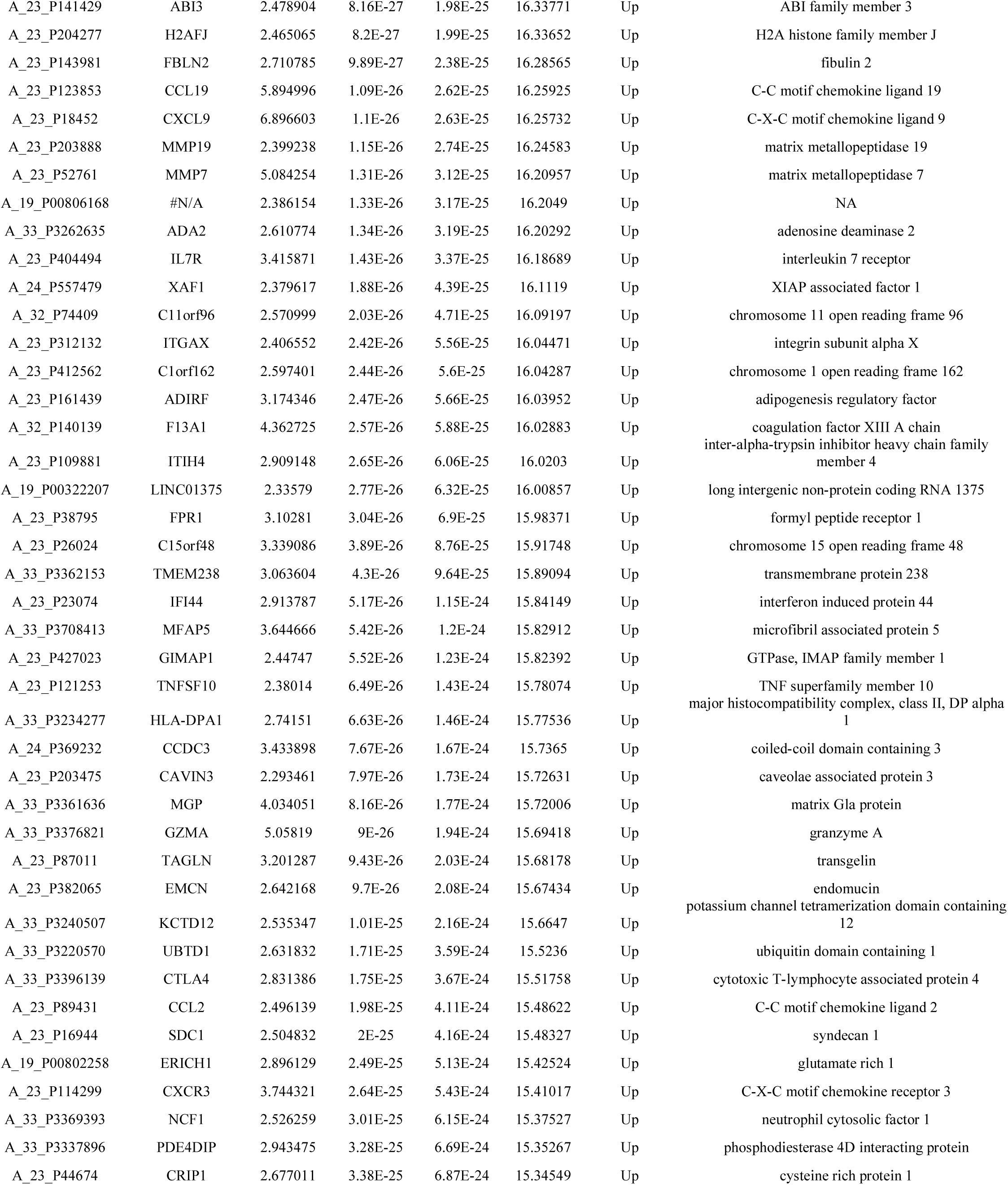
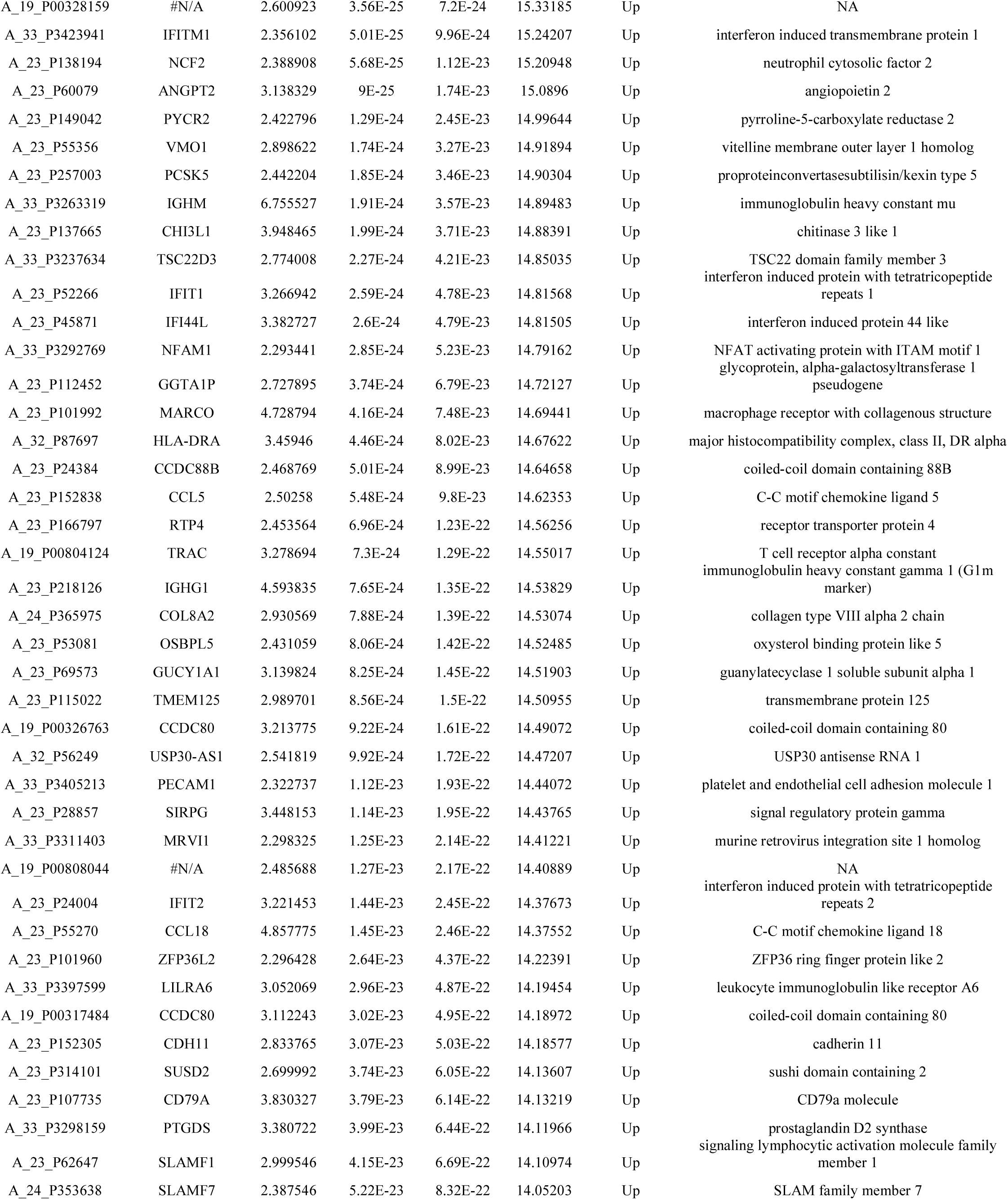
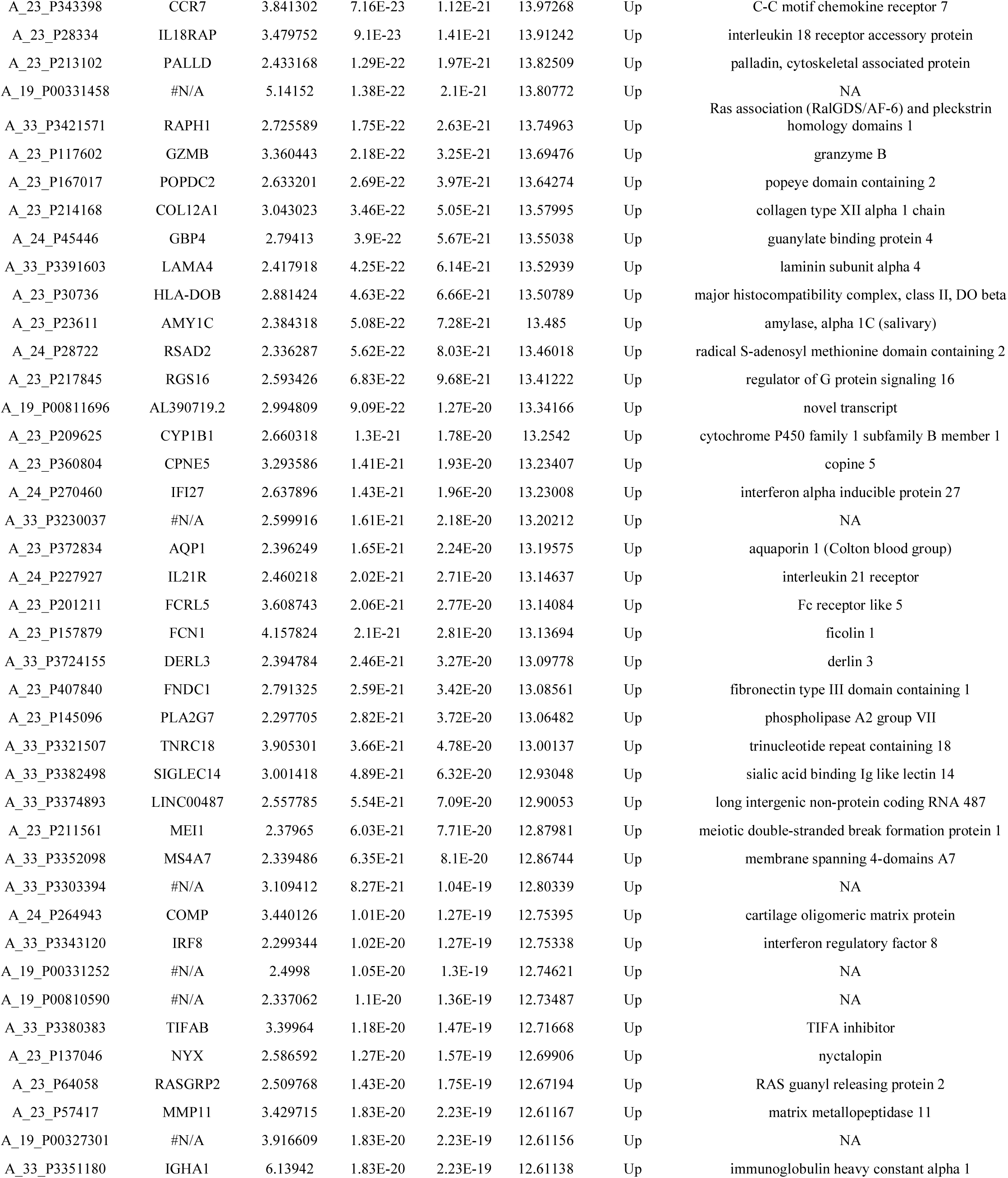
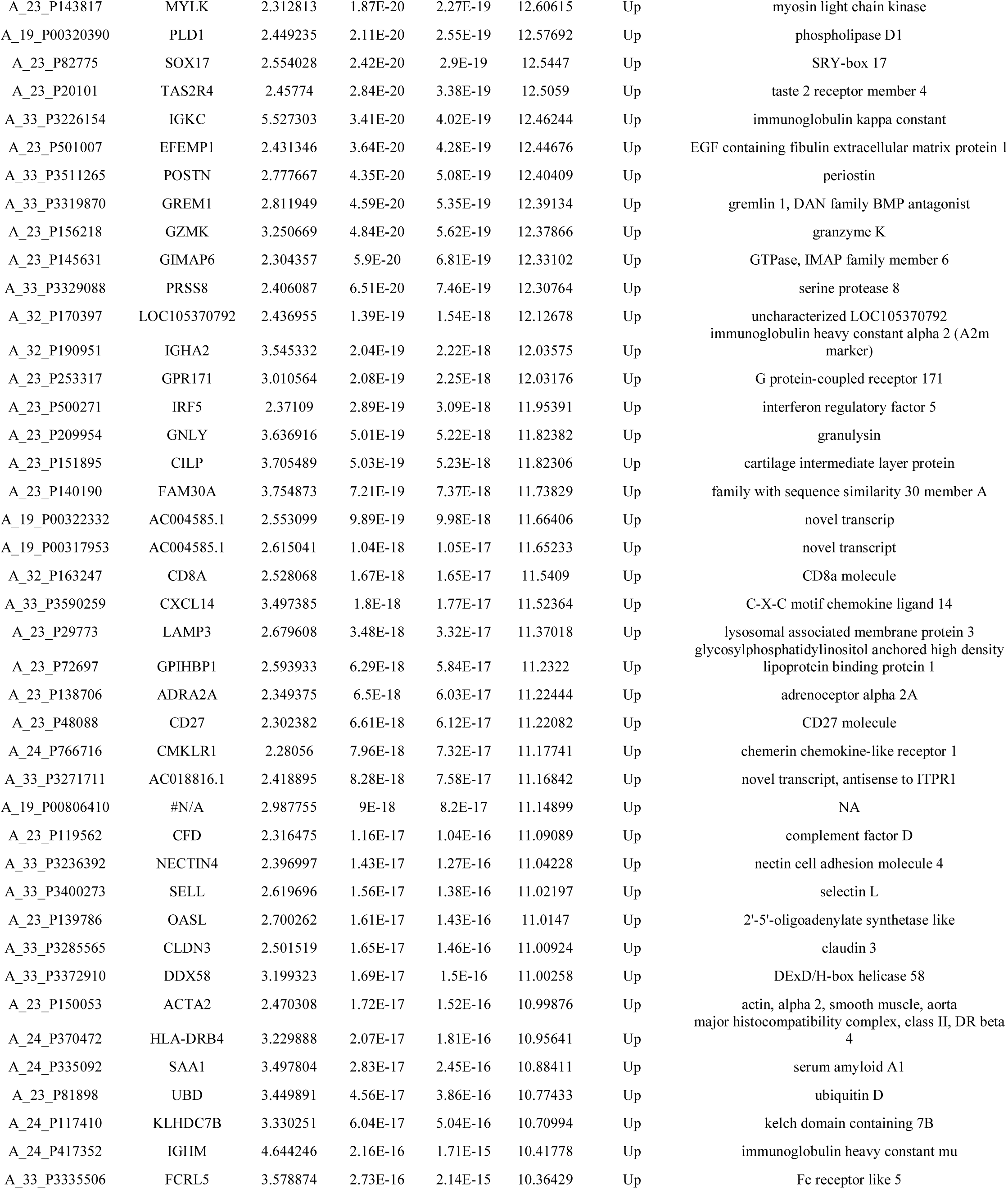
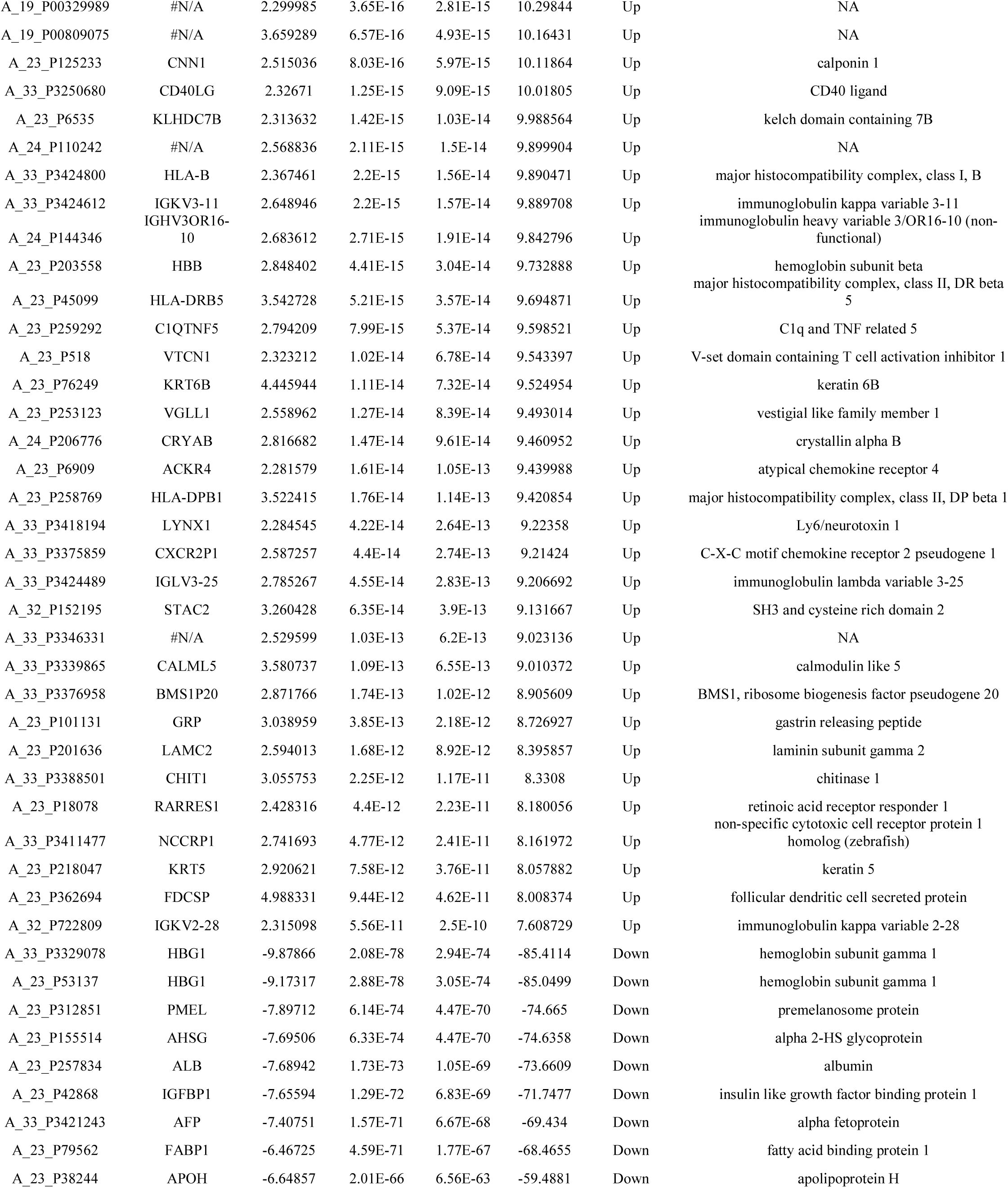
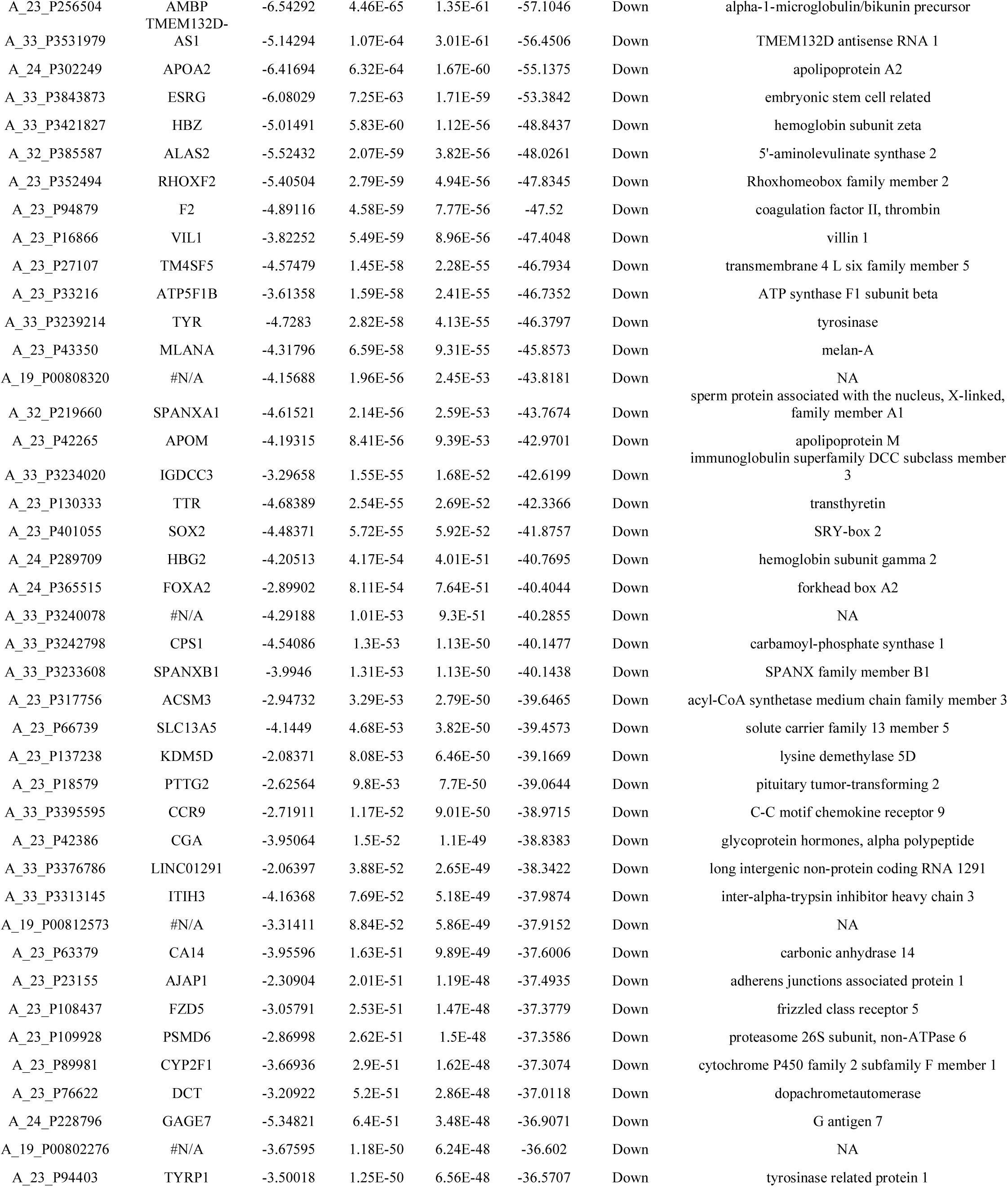
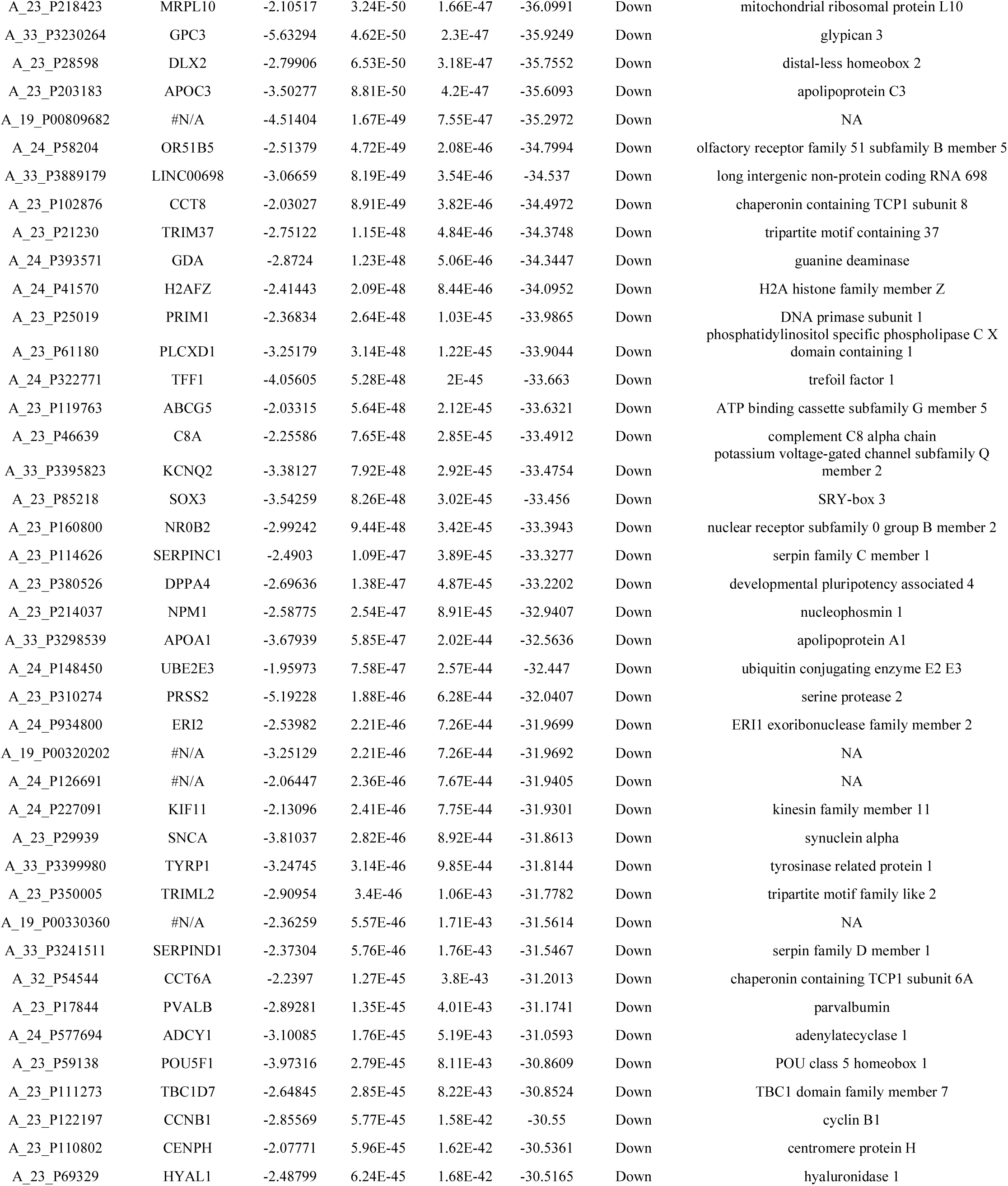
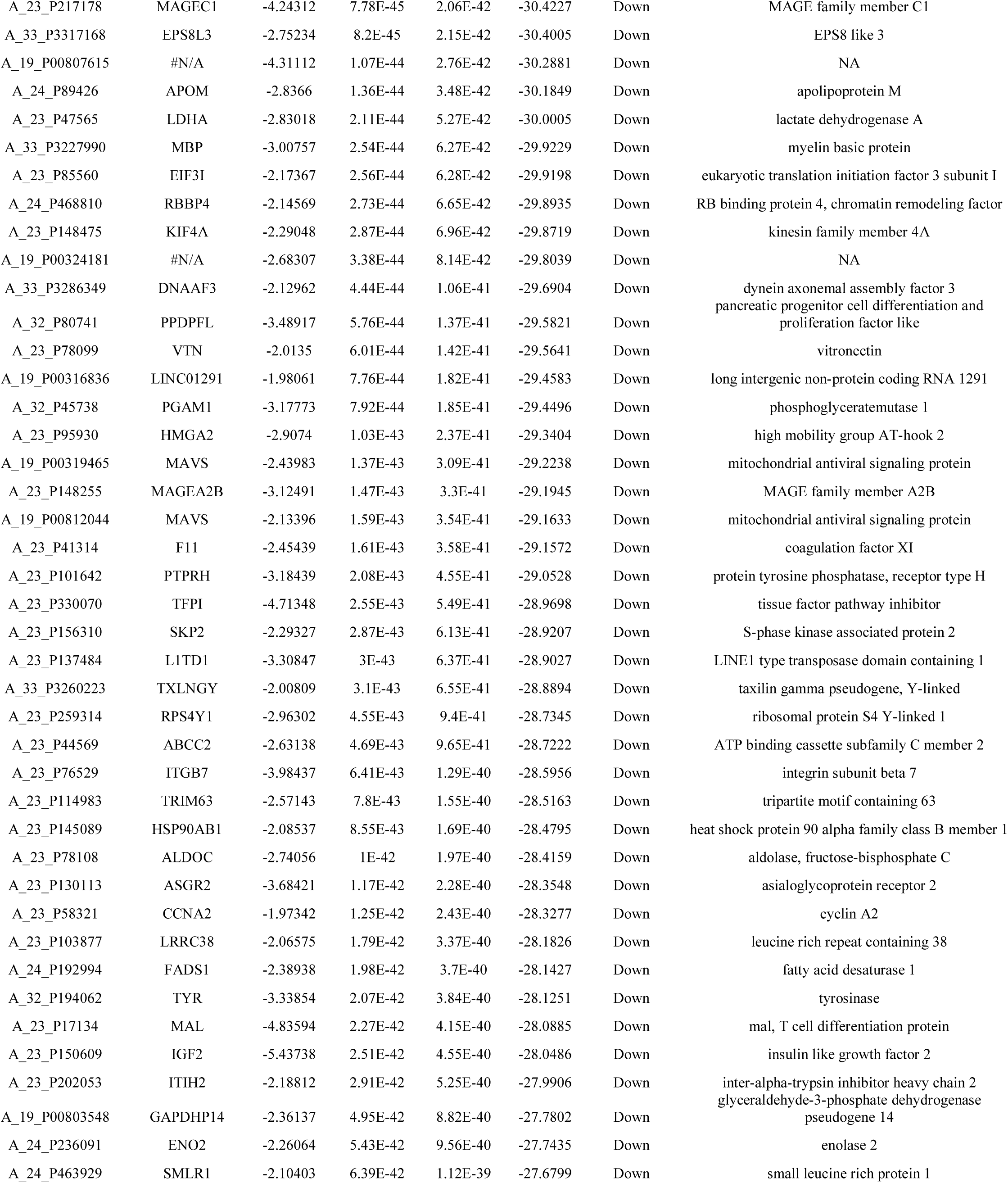
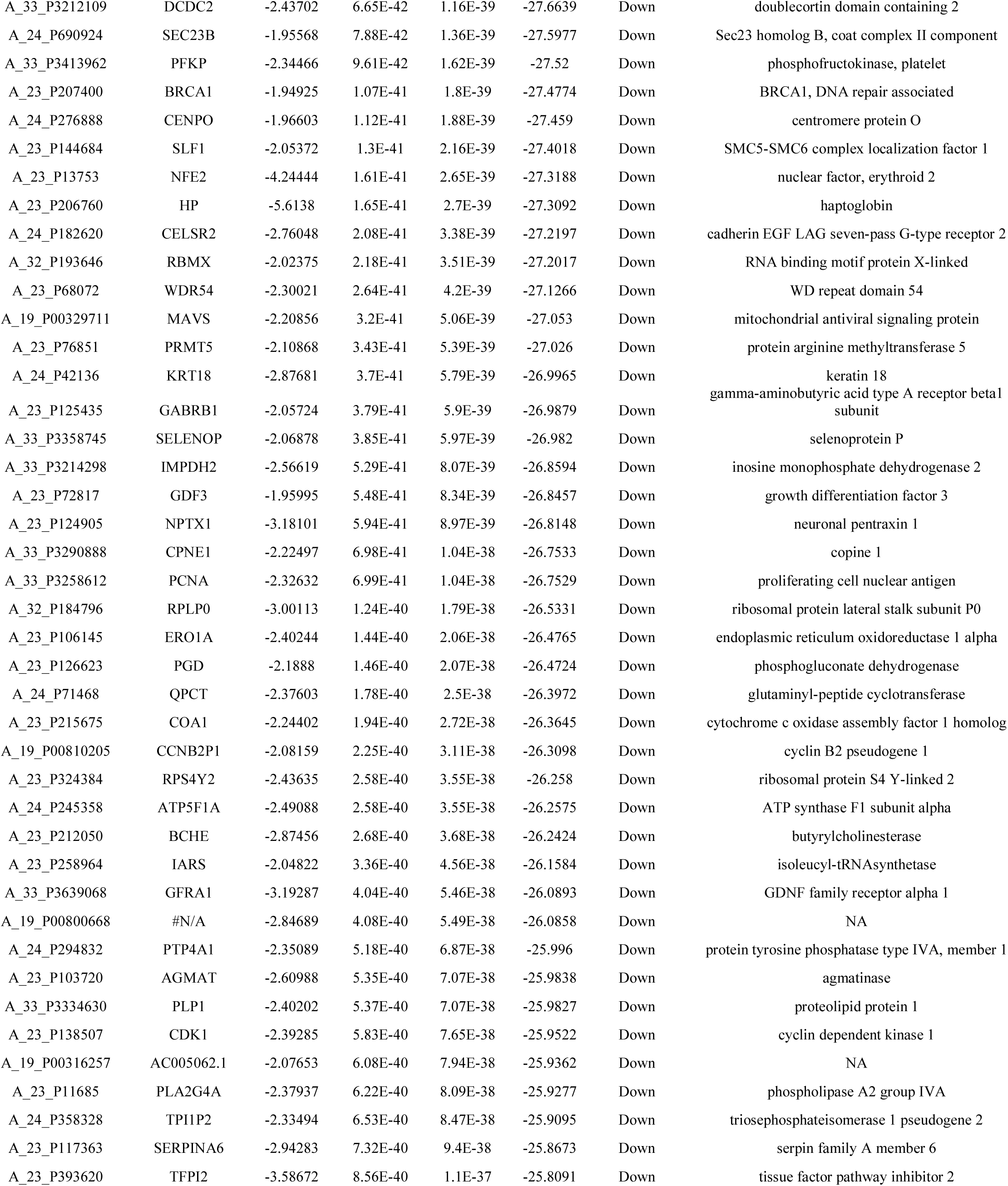
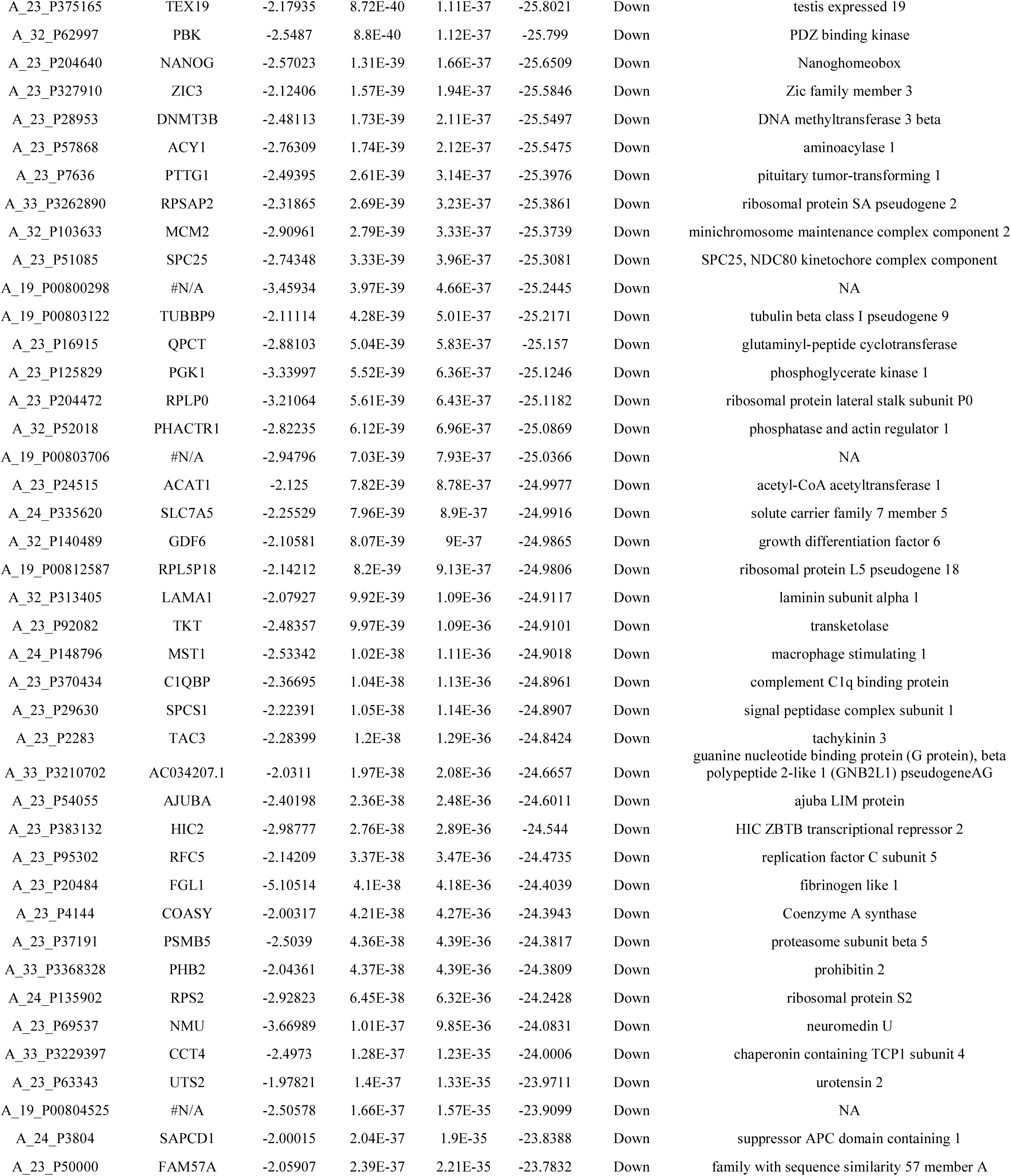
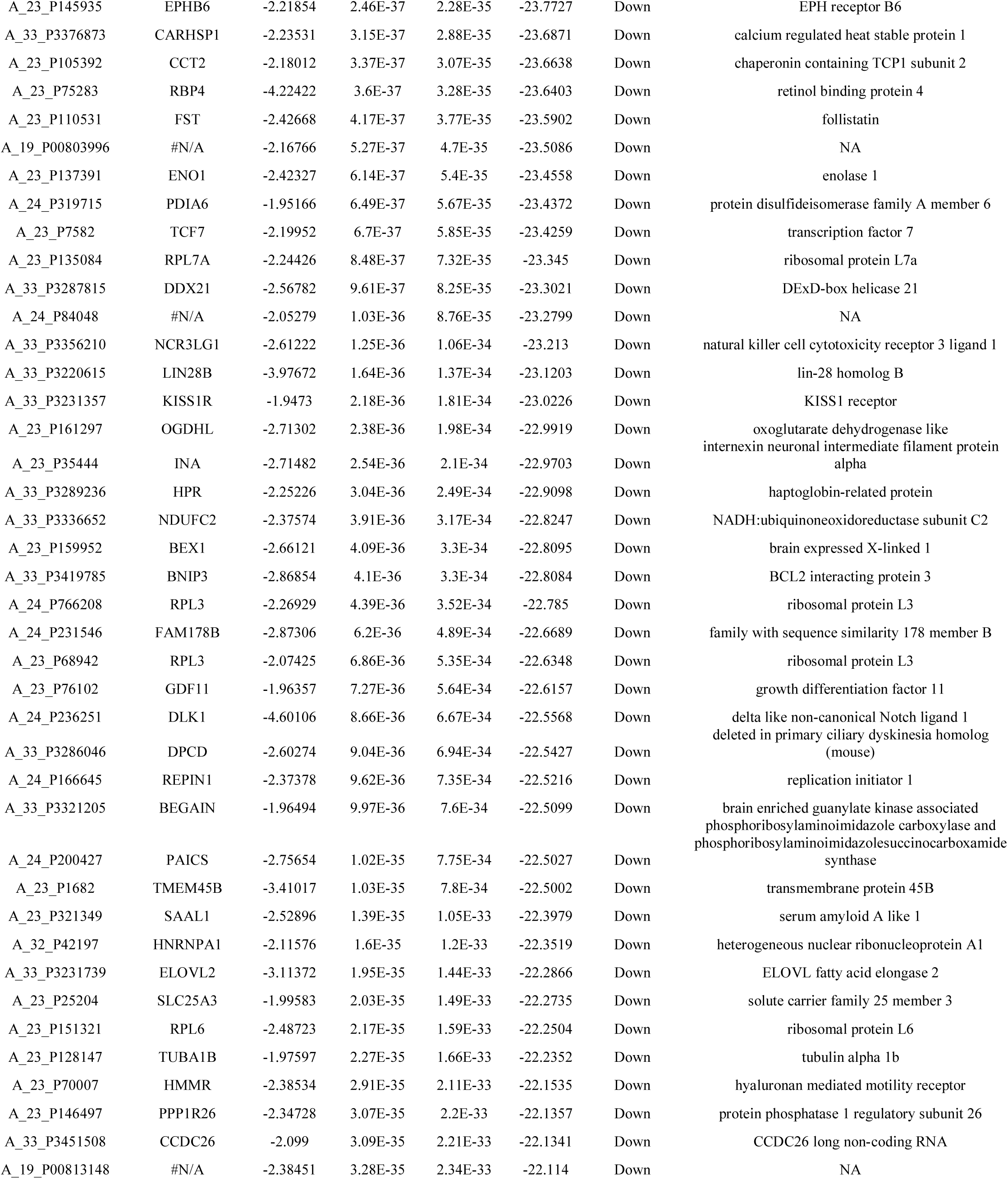
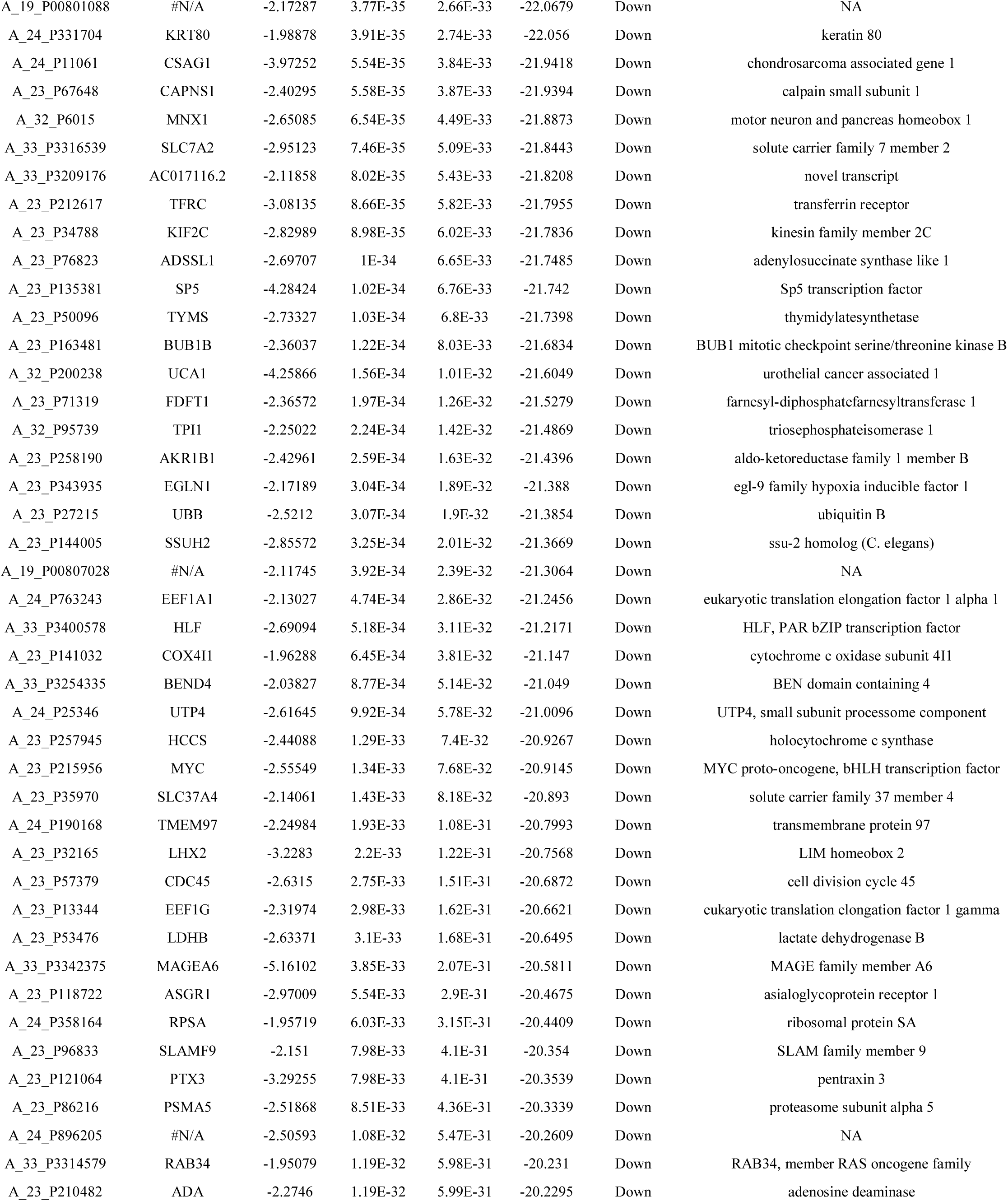
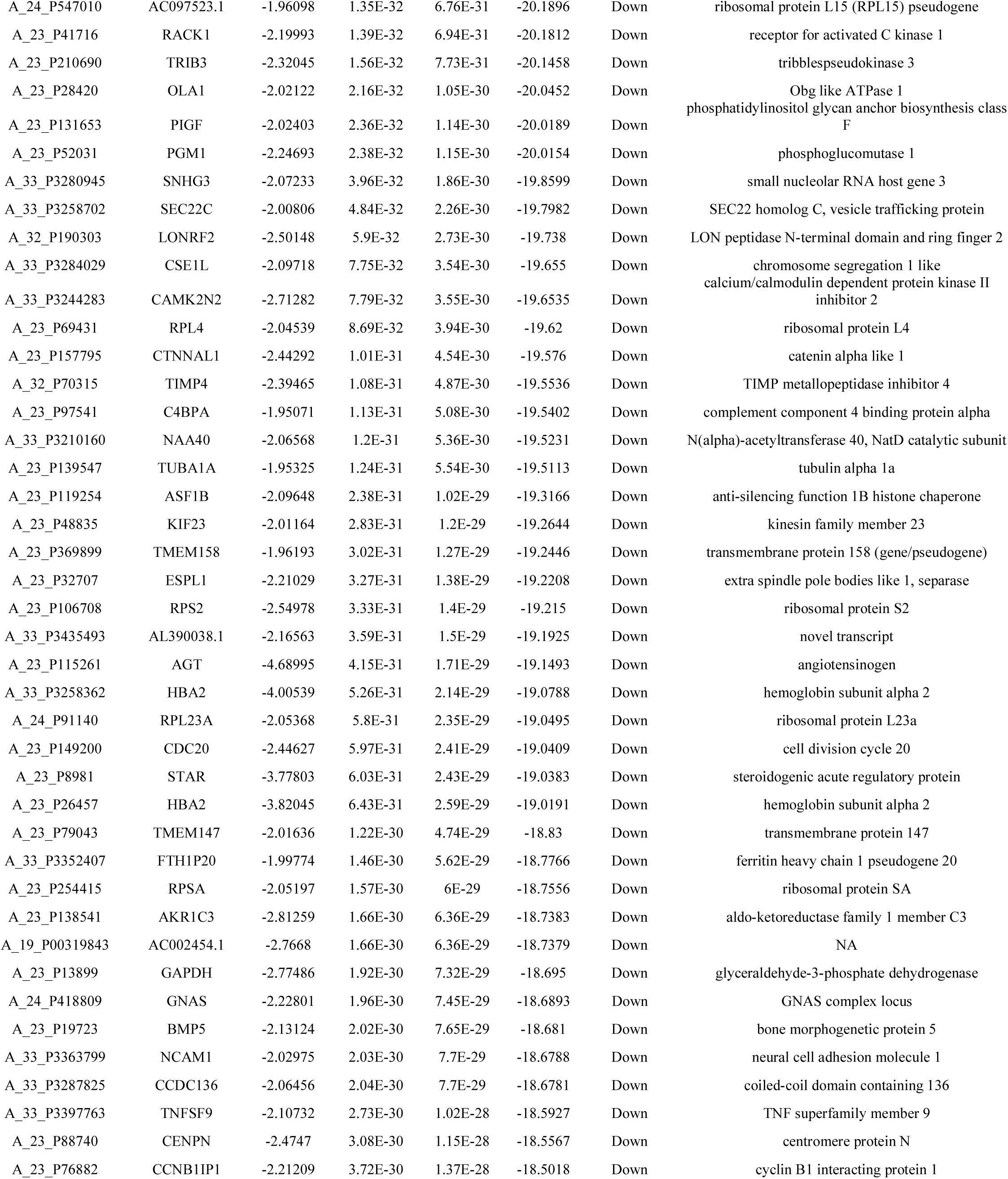
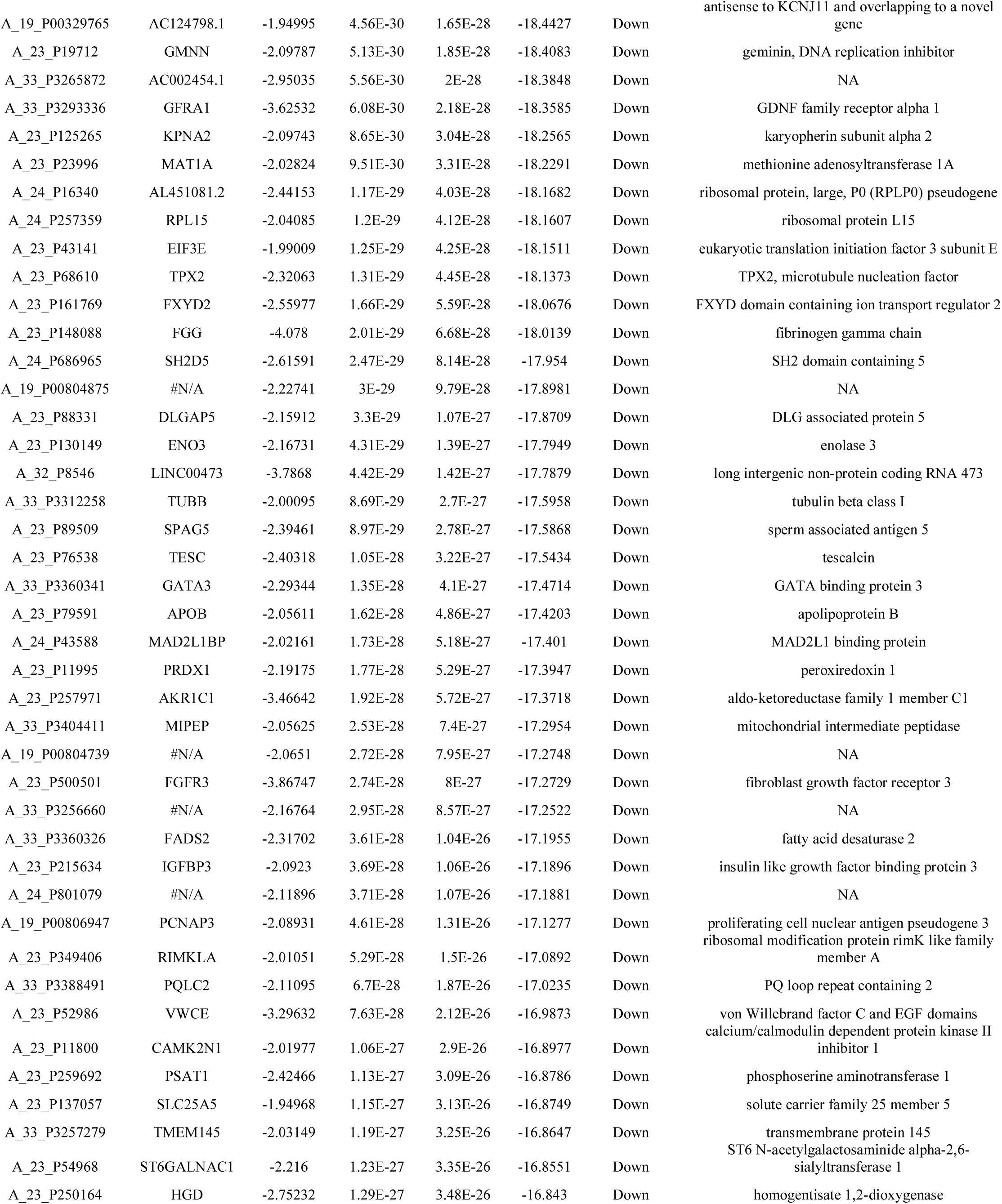
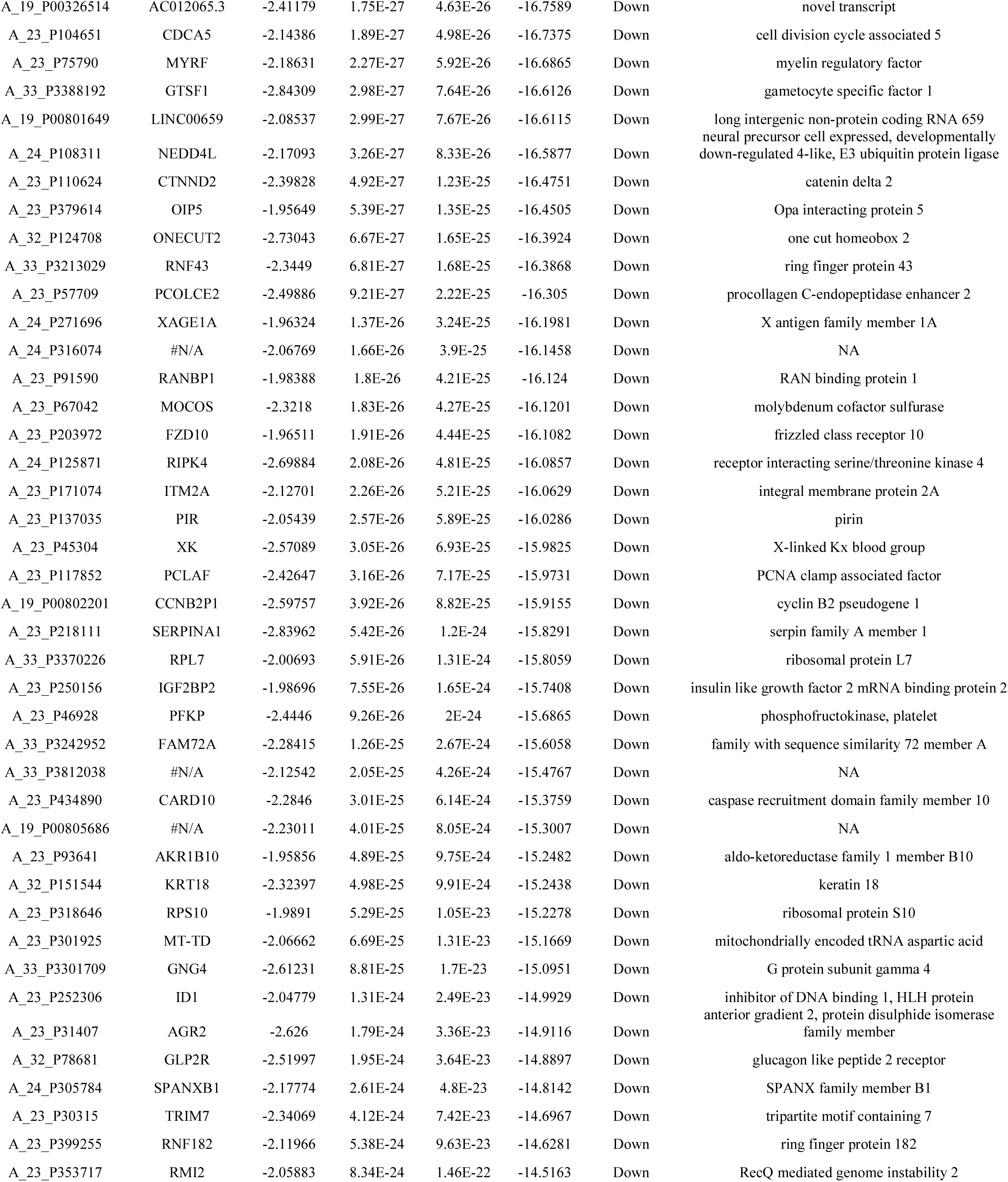
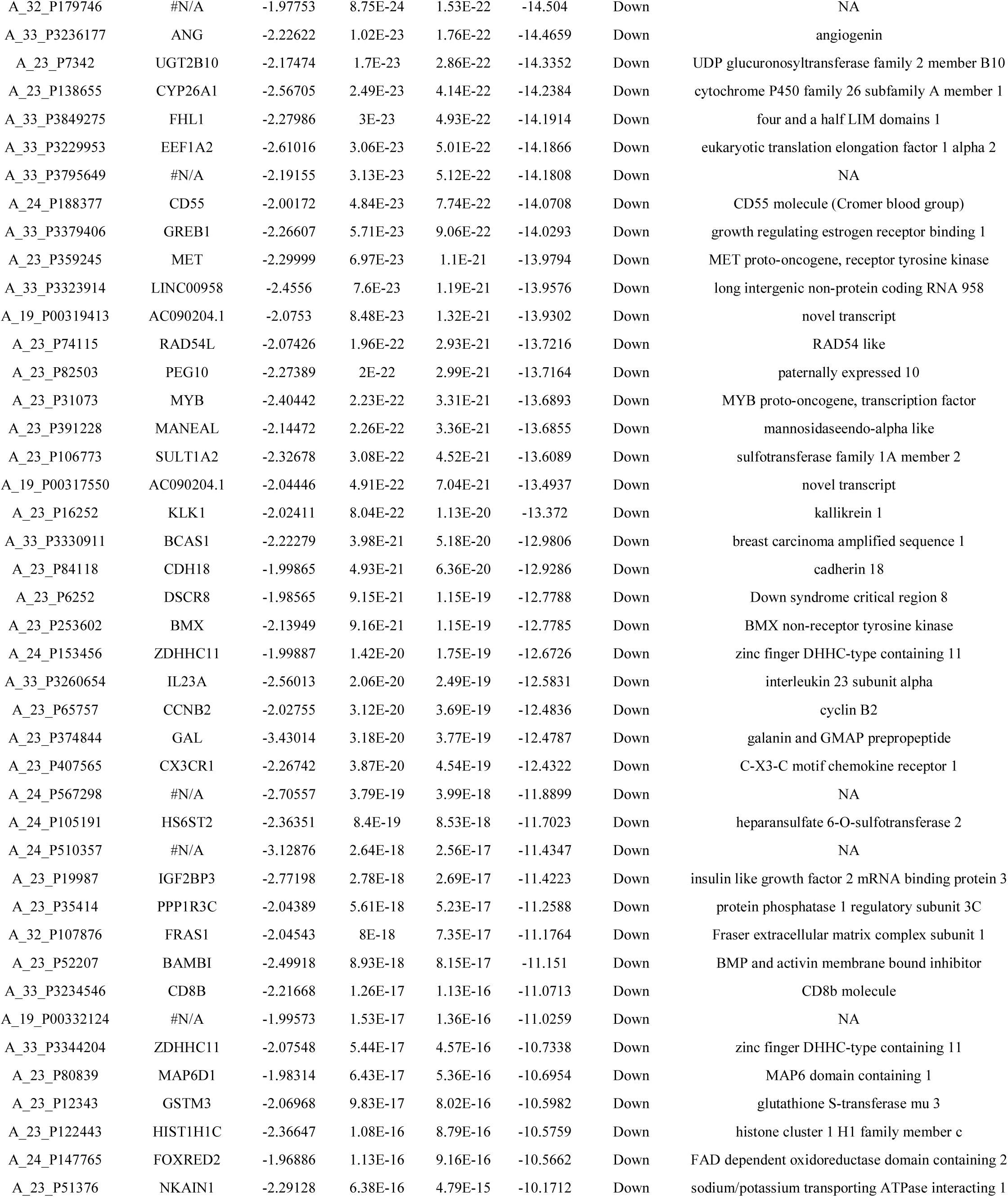
The statistical metrics for key differentially expressed genes (DEGs)

### Pathway enrichment analysis of DEGs

For a deeper insight into the DEGs (up and down regulated genes), we performed pathway enrichment analyses. Pathway enrichment analyses revealed that up regulated genes were mainly associated with phospholipases, resolvin D biosynthesis, toxoplasmosis, antigen processing and presentation, IL12-mediated signaling events, CXCR4-mediated signaling events, innate immune system, assembly of collagen fibrils and other multimeric structures, prostaglandin and leukotriene metabolism, sphingoglycolipid metabolism, ensemble of genes encoding core extracellular matrix including ECM glycoproteins, collagens and proteoglycans, genes encoding collagen proteins, integrin signalling pathway, inflammation mediated by chemokine and cytokine signaling pathway, eicosanoids metabolic, gamma-hexachlorocyclohexane degradation, MNGIE (mitochondrial neurogastrointestinal encephalopathy) and intracellular signalling through PGD2 receptor and prostaglandin D2 are listed in Table 3; while down regulated genes were mainly associated with glycolysis, lactate fermentation (reoxidation of cytosolic NADH), biosynthesis of amino acids, carbon metabolism, FOXA2 and FOXA3 transcription factor networks, HIF-1-alpha transcription factor network, peptide chain elongation, cell cycle, glycolysis gluconeogenesis, pentose phosphate, extrinsic prothrombin activation pathway, ATP synthesis and enoxaparin pathway are listed in Table 4.

**Table 3.**
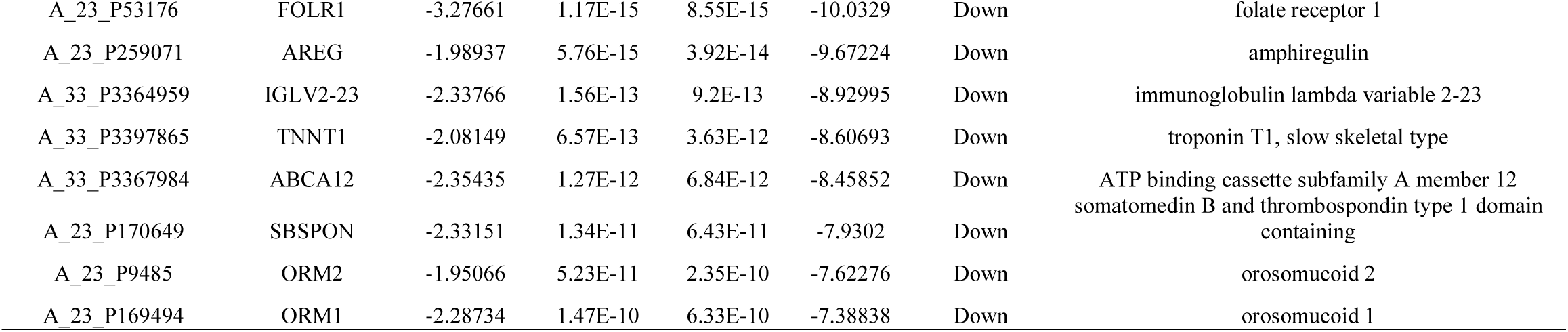
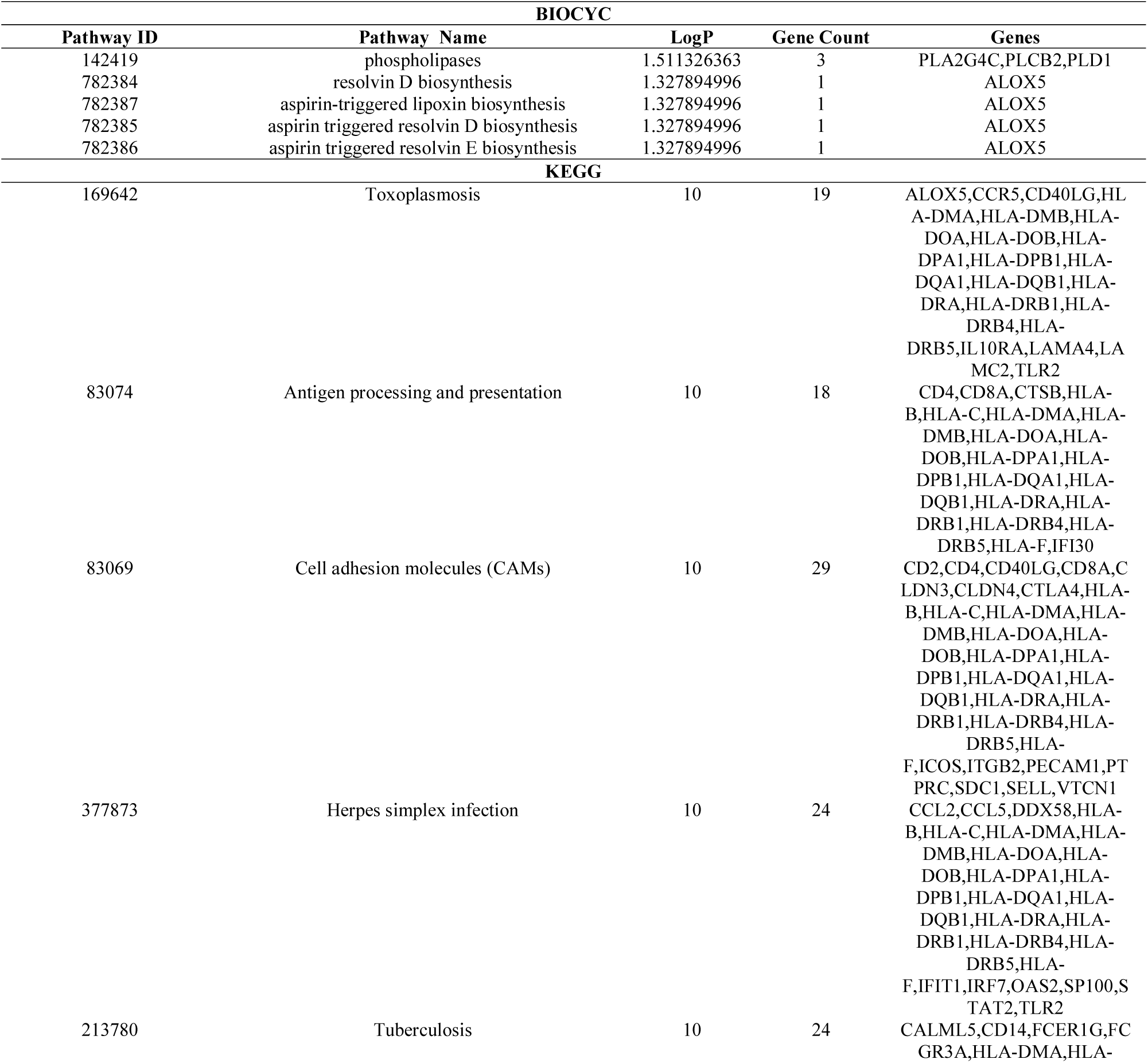
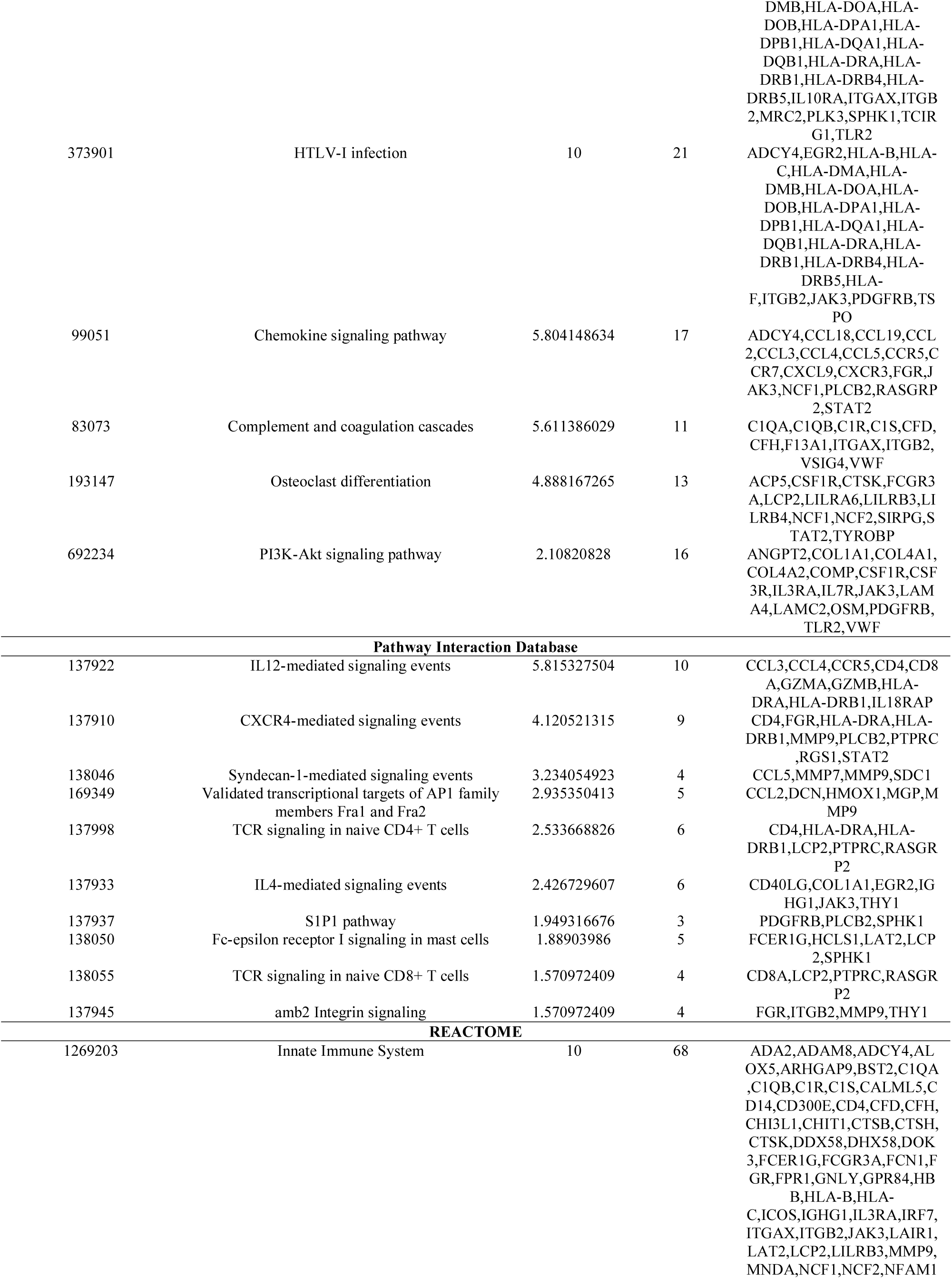
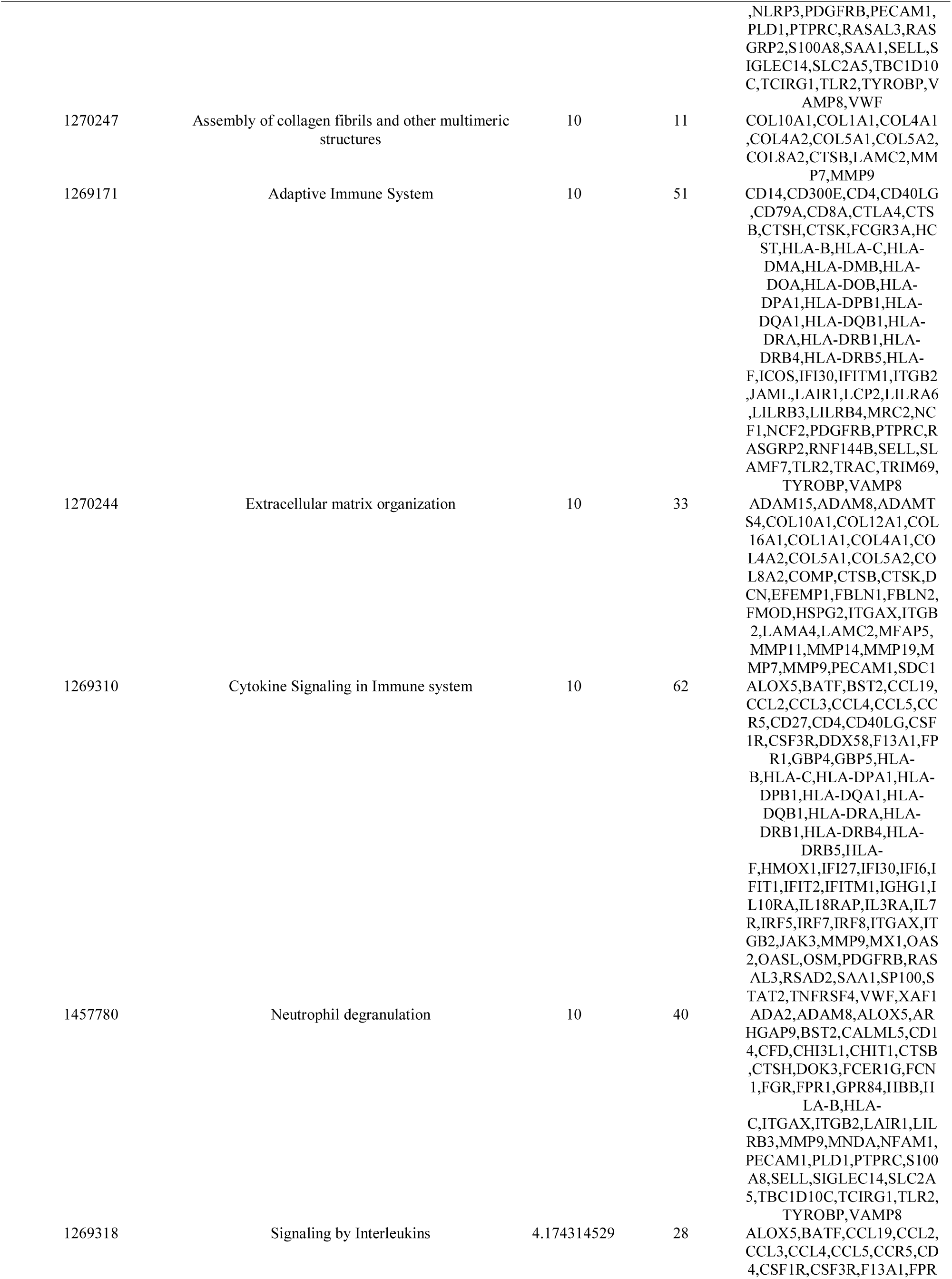
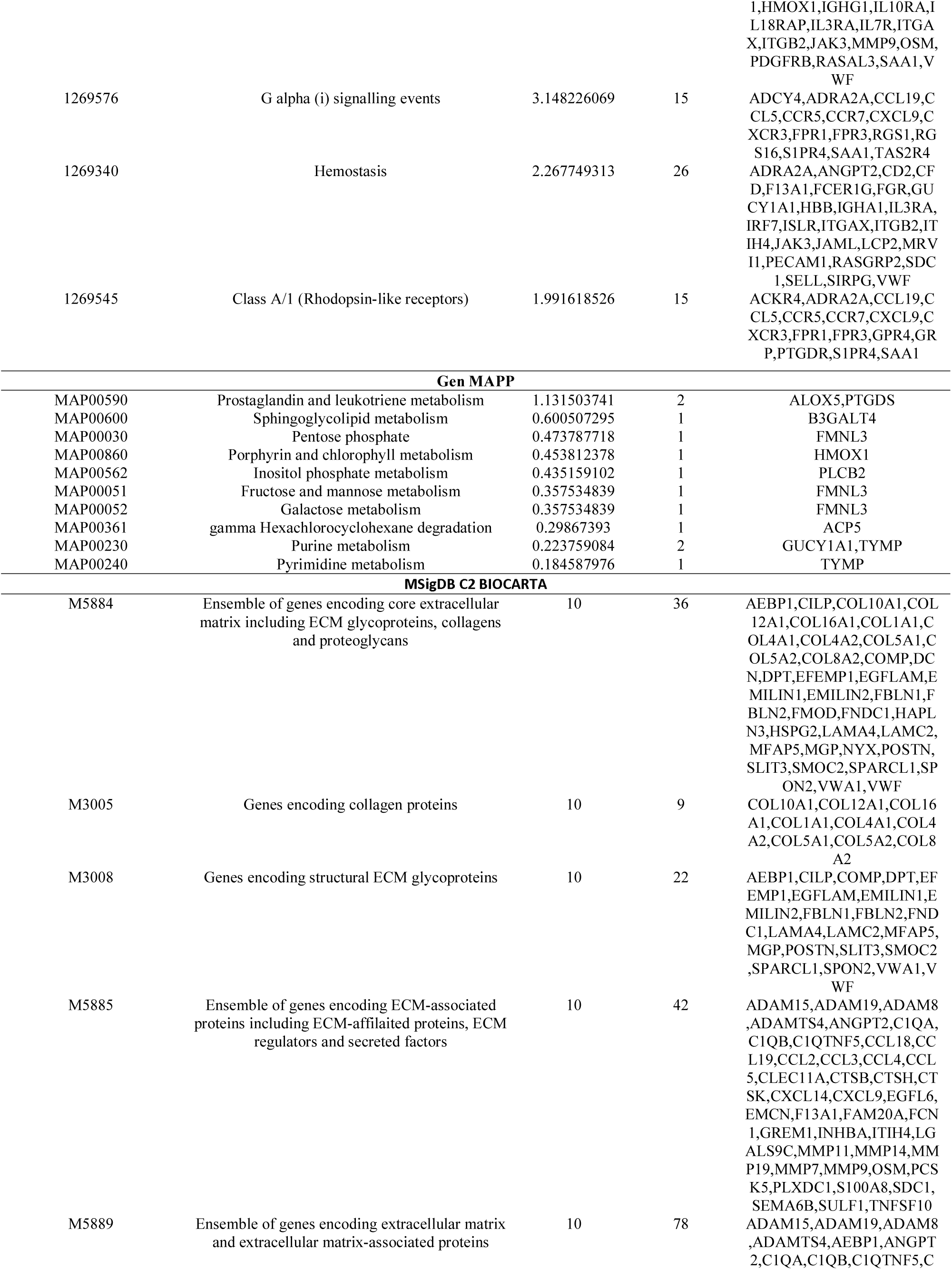
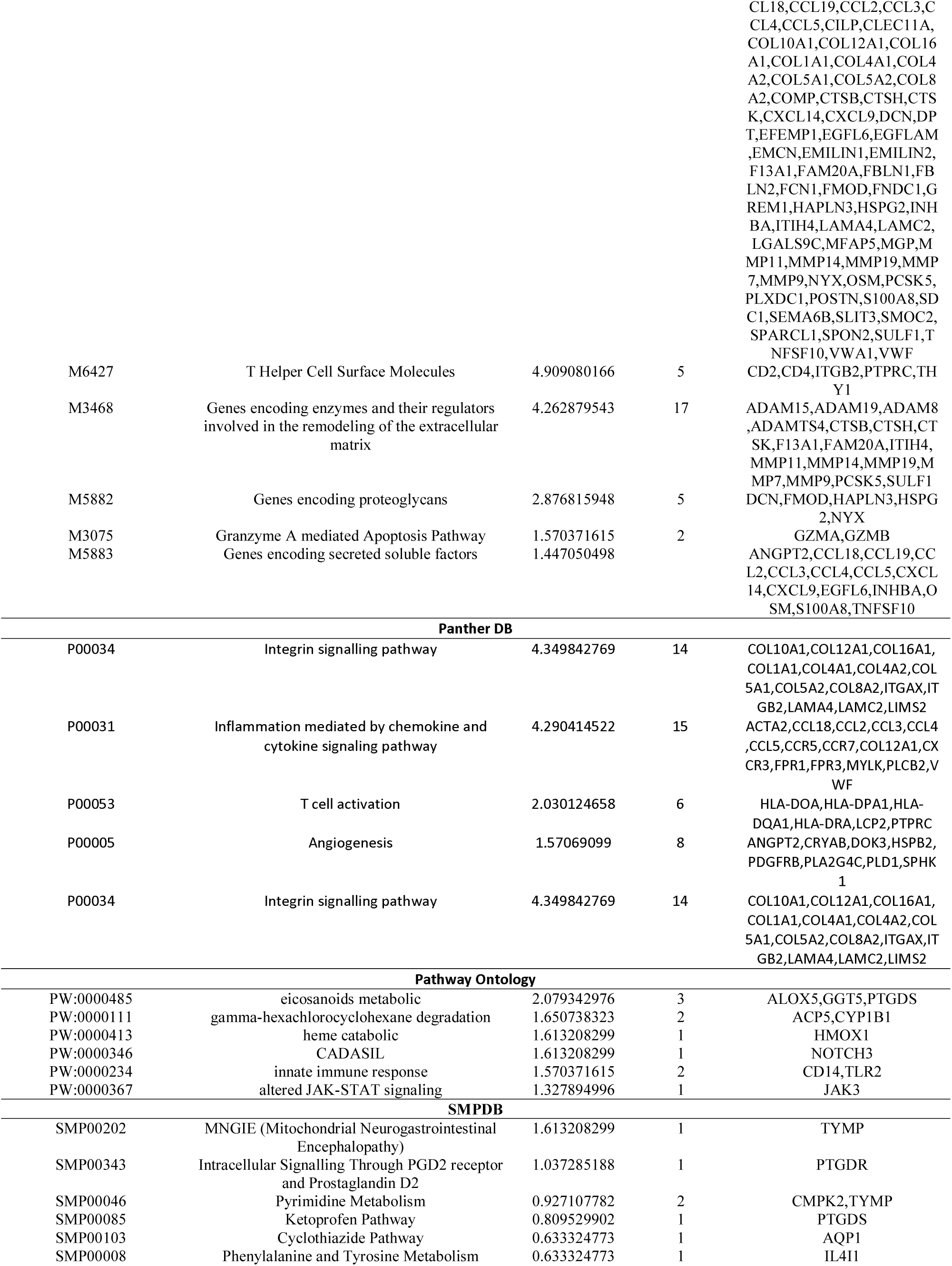
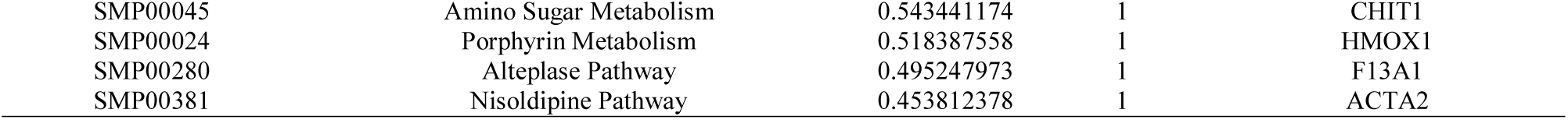
The enriched pathway terms of the up regulated differentially expressed genes

**Table 4.**
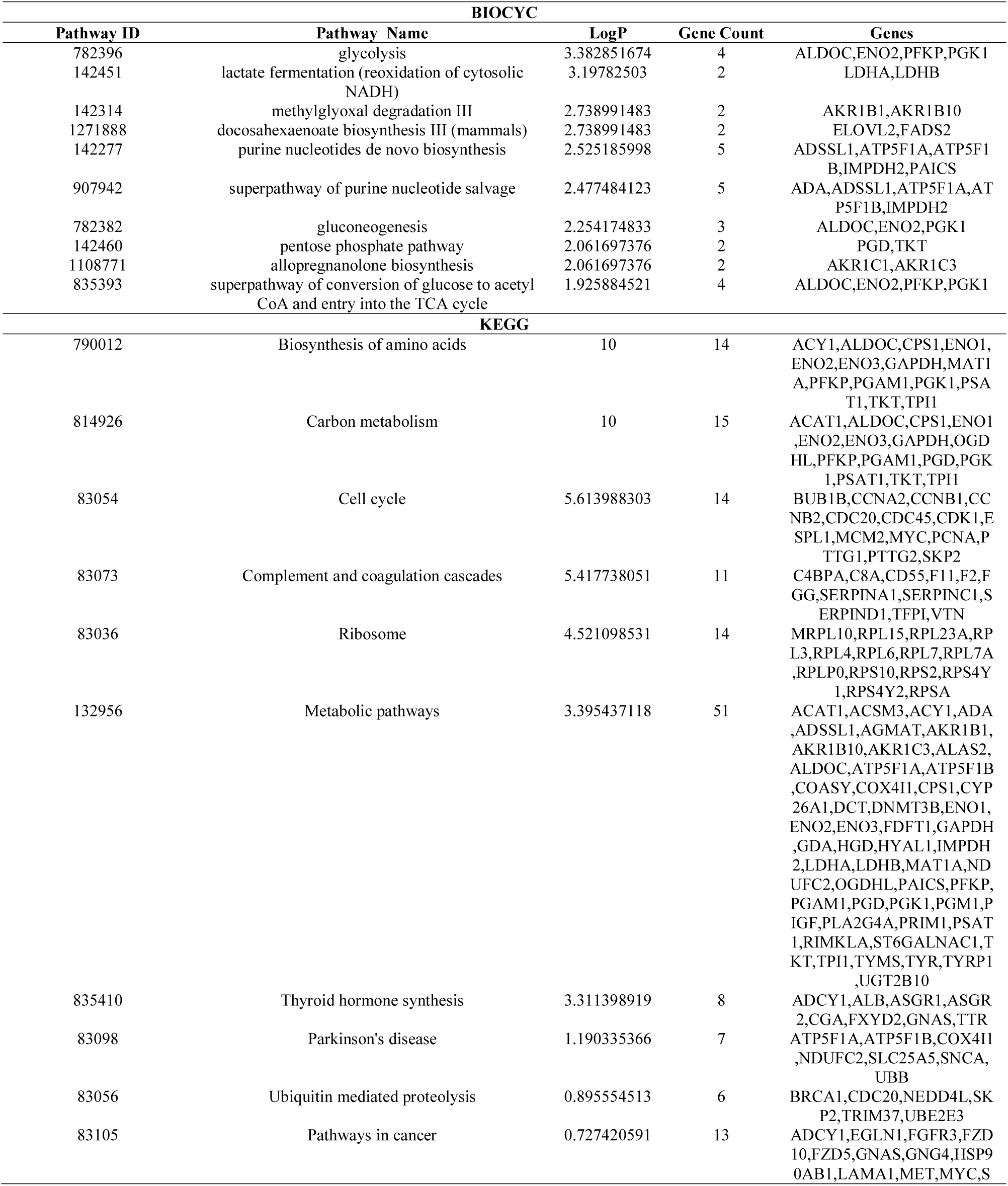
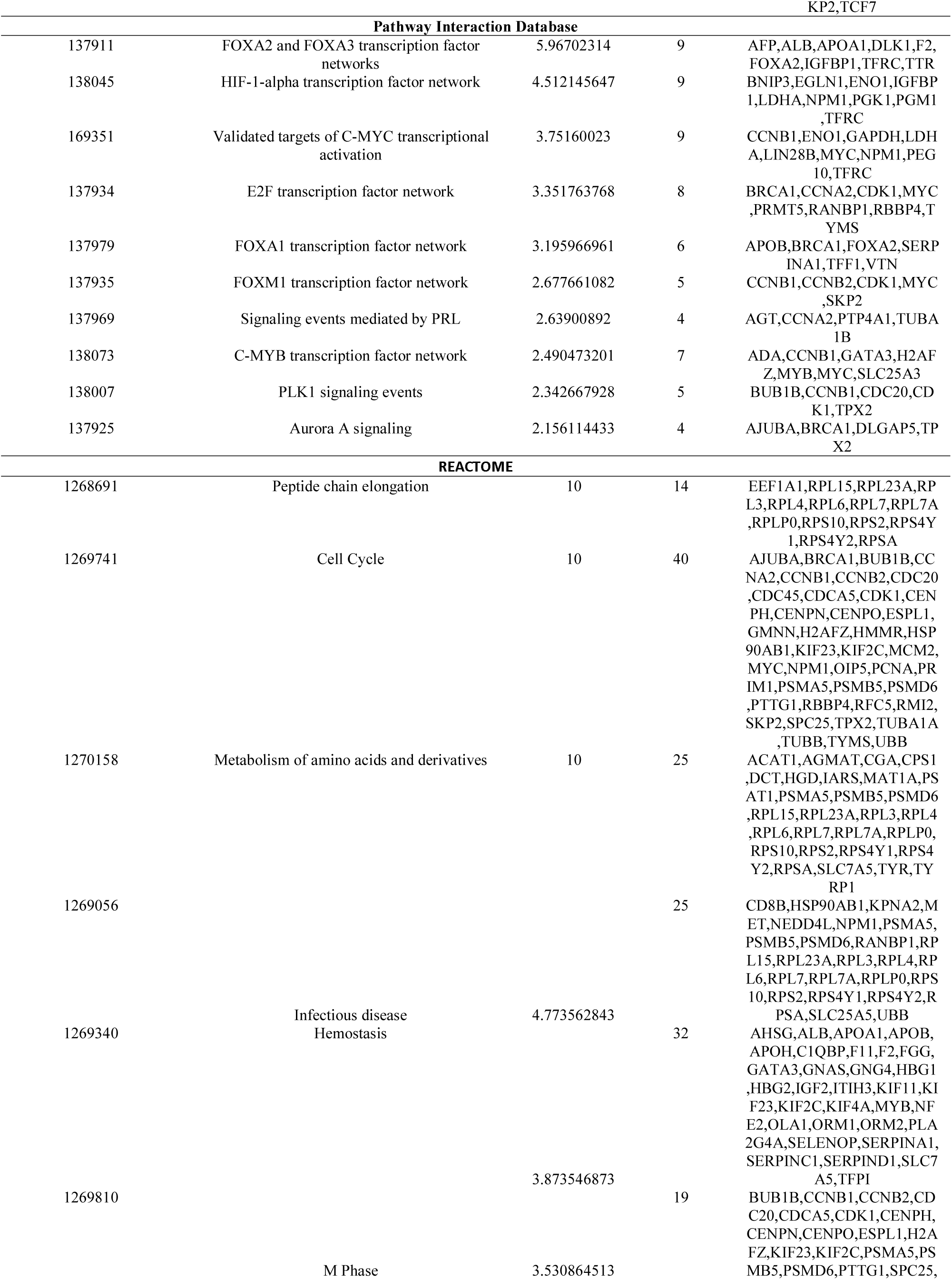
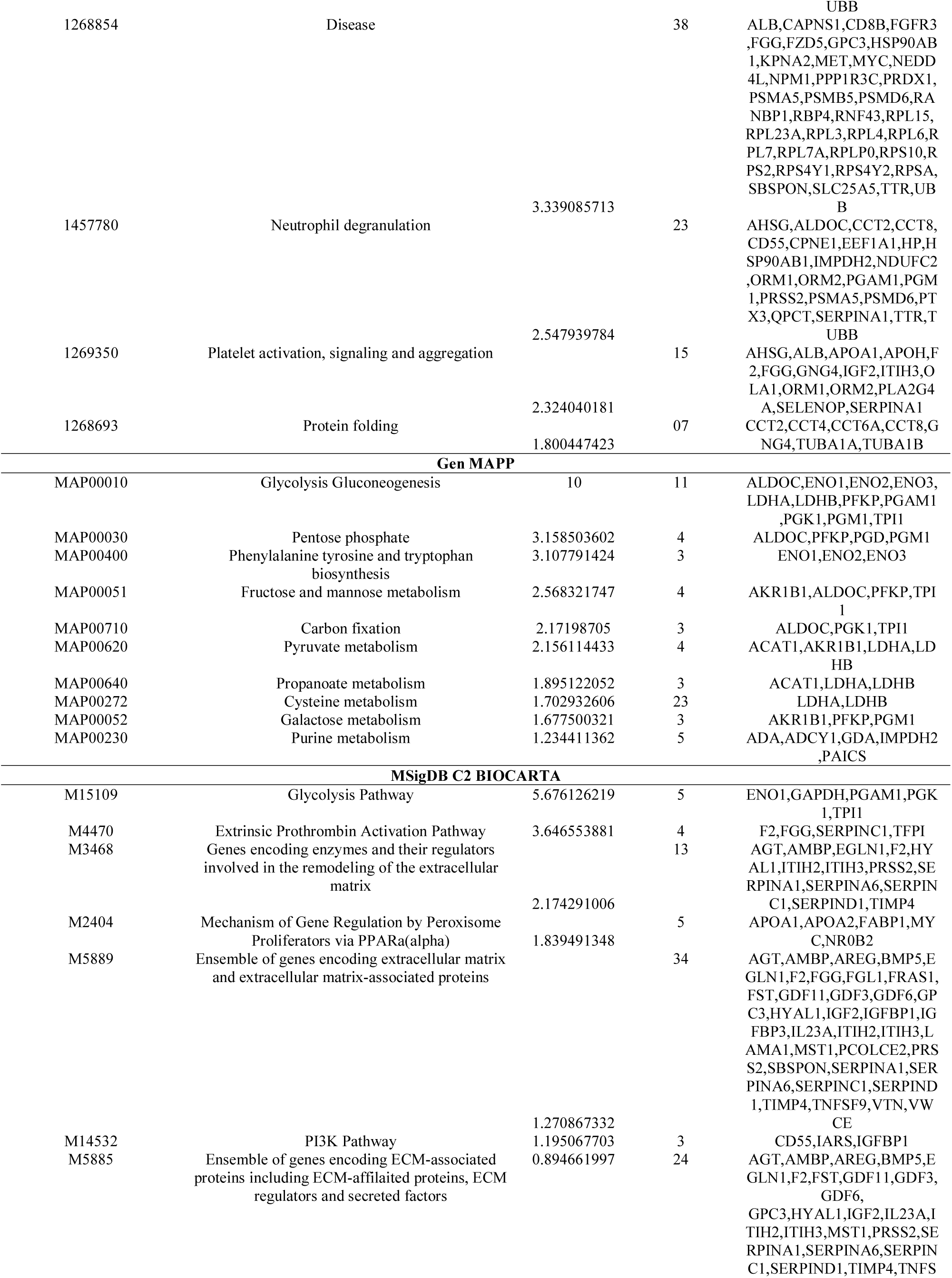
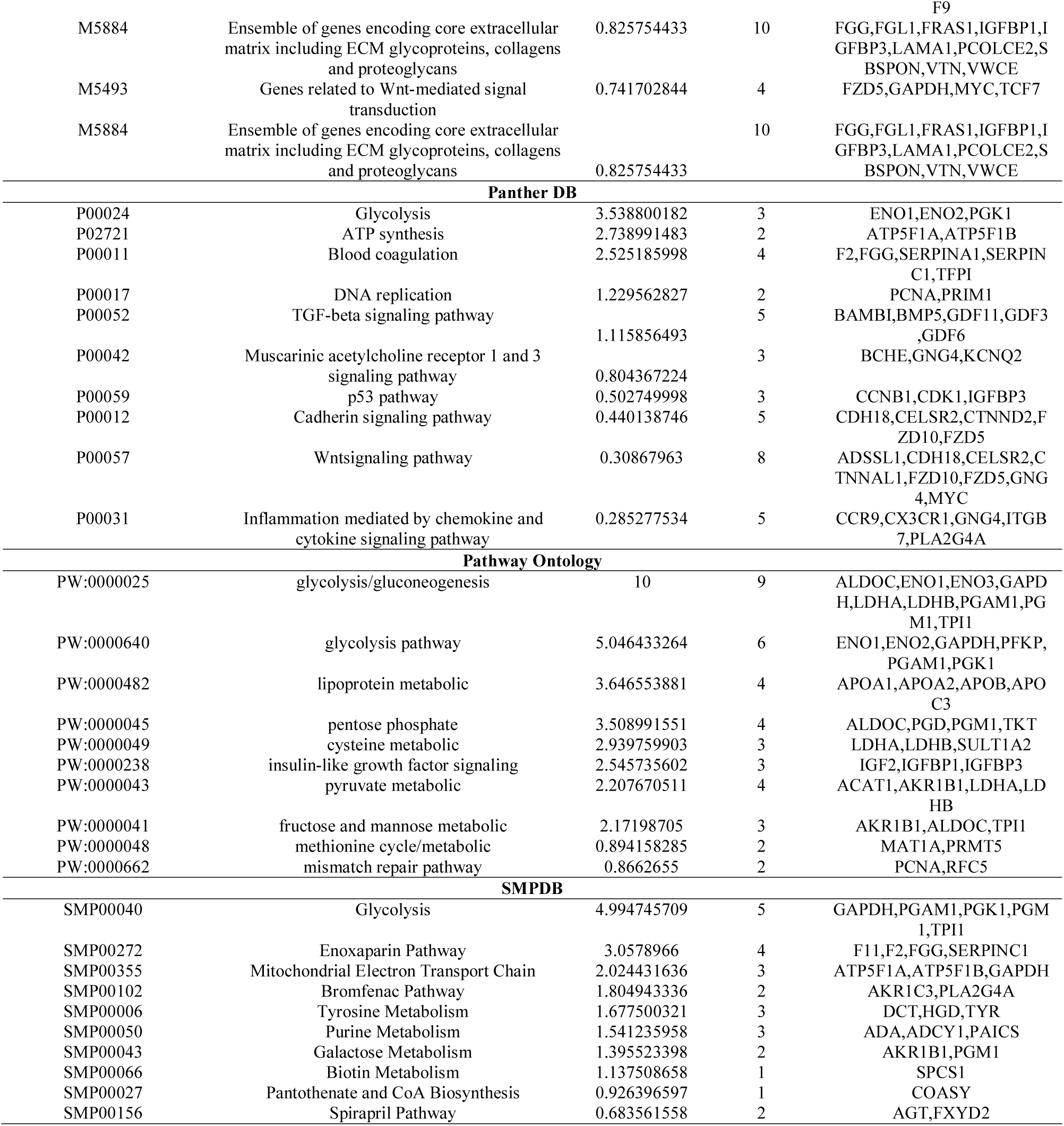
The enriched pathway terms of the down regulated differentially expressed genes

### Gene ontology (GO) enrichment analysis of DEGs

In order to investigate the biological functions of these up and down regulated genes in TNBC, GO enrichment analysis was performed using ToppCluster (Table 5 and Table 6). For BPs, GO analysis results indicated that up and down regulated genes were significantly enriched in immune response, regulation of immune system process, carboxylic acid metabolic process and oxoacid metabolic process. CC analysis showed that these up and down regulated genes were particularly associated in cell surface, side of membrane, extracellular space and cytosolic part. Similarly, changes in MF of up and down regulated genes were significantly enriched in serine-type peptidase activity, antigen binding, protein-containing complex binding and enzyme binding.

**Table 5.**
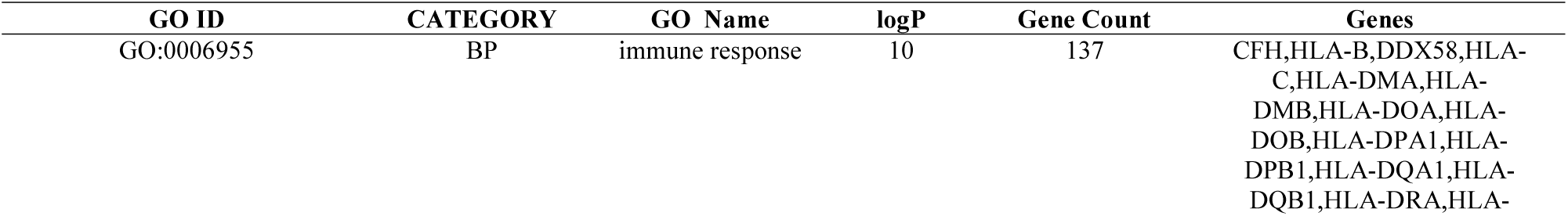
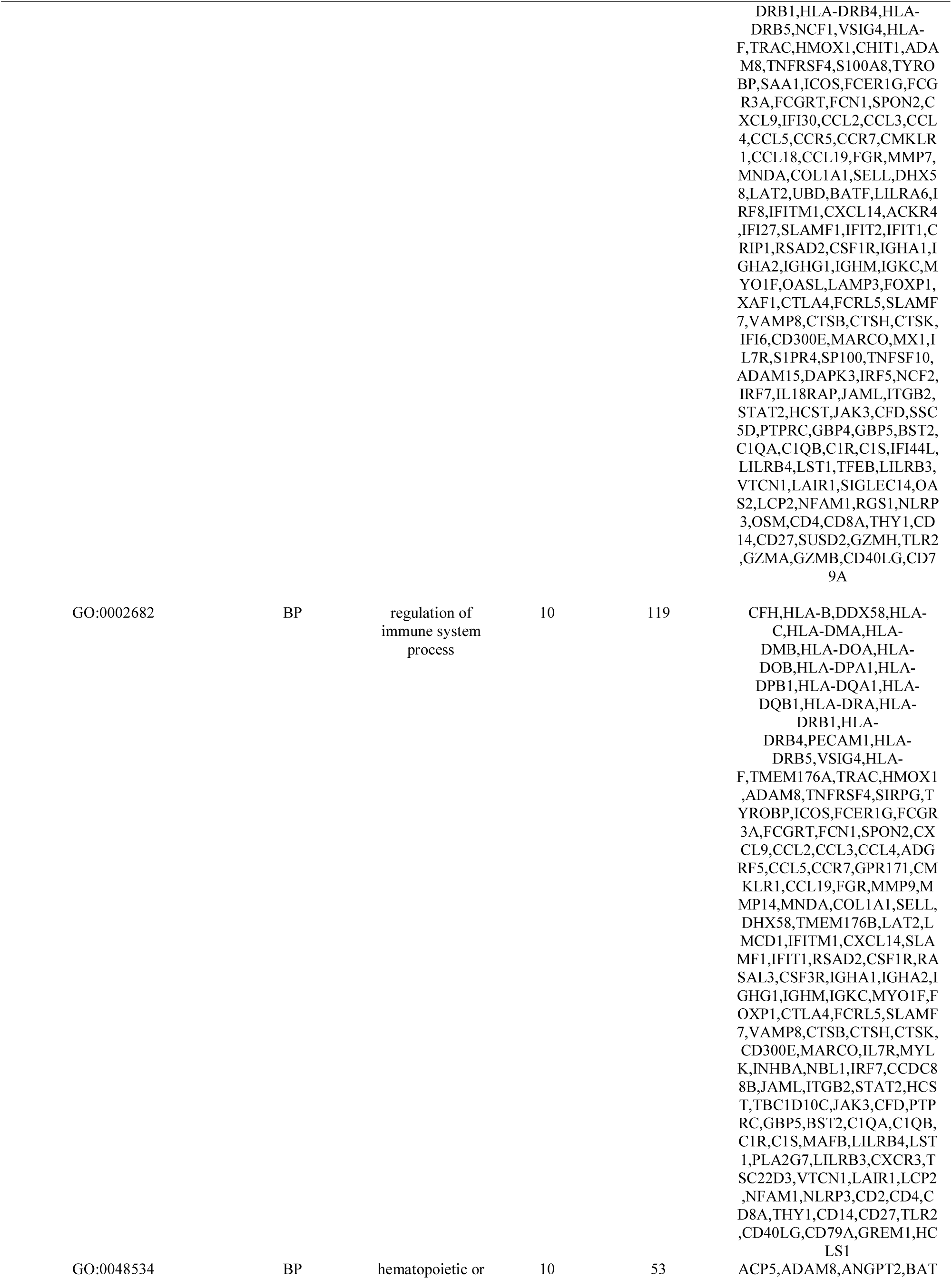
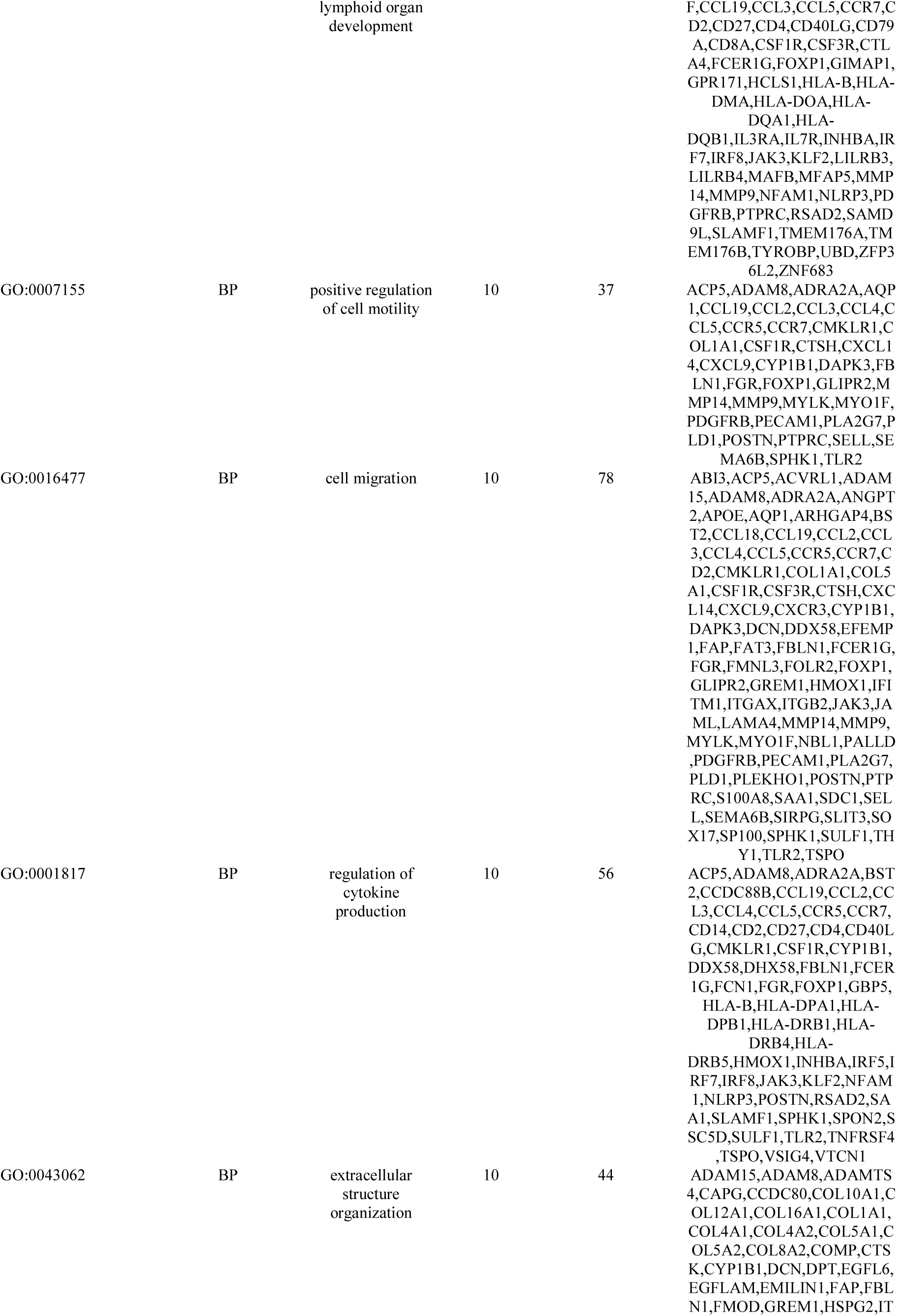
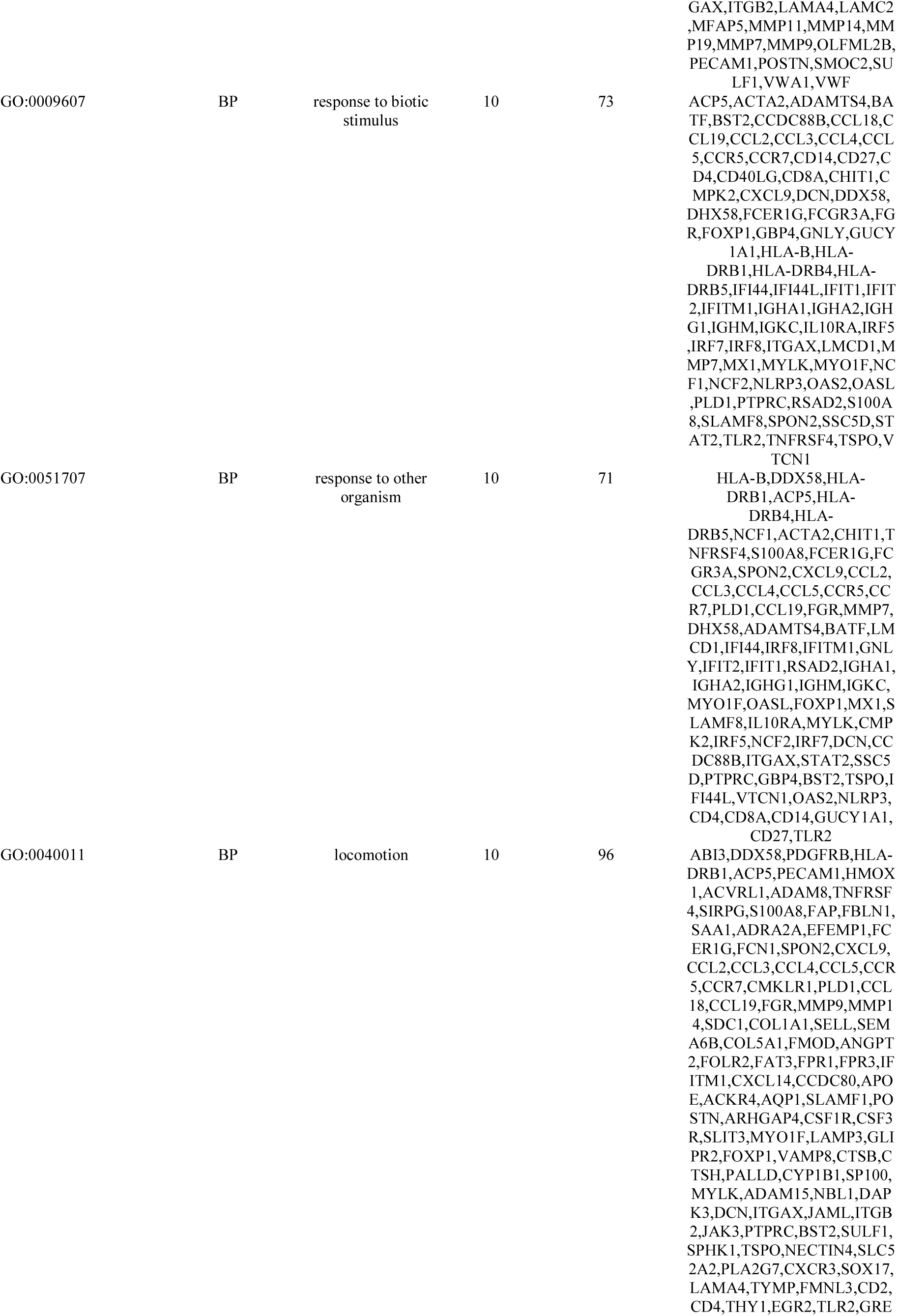
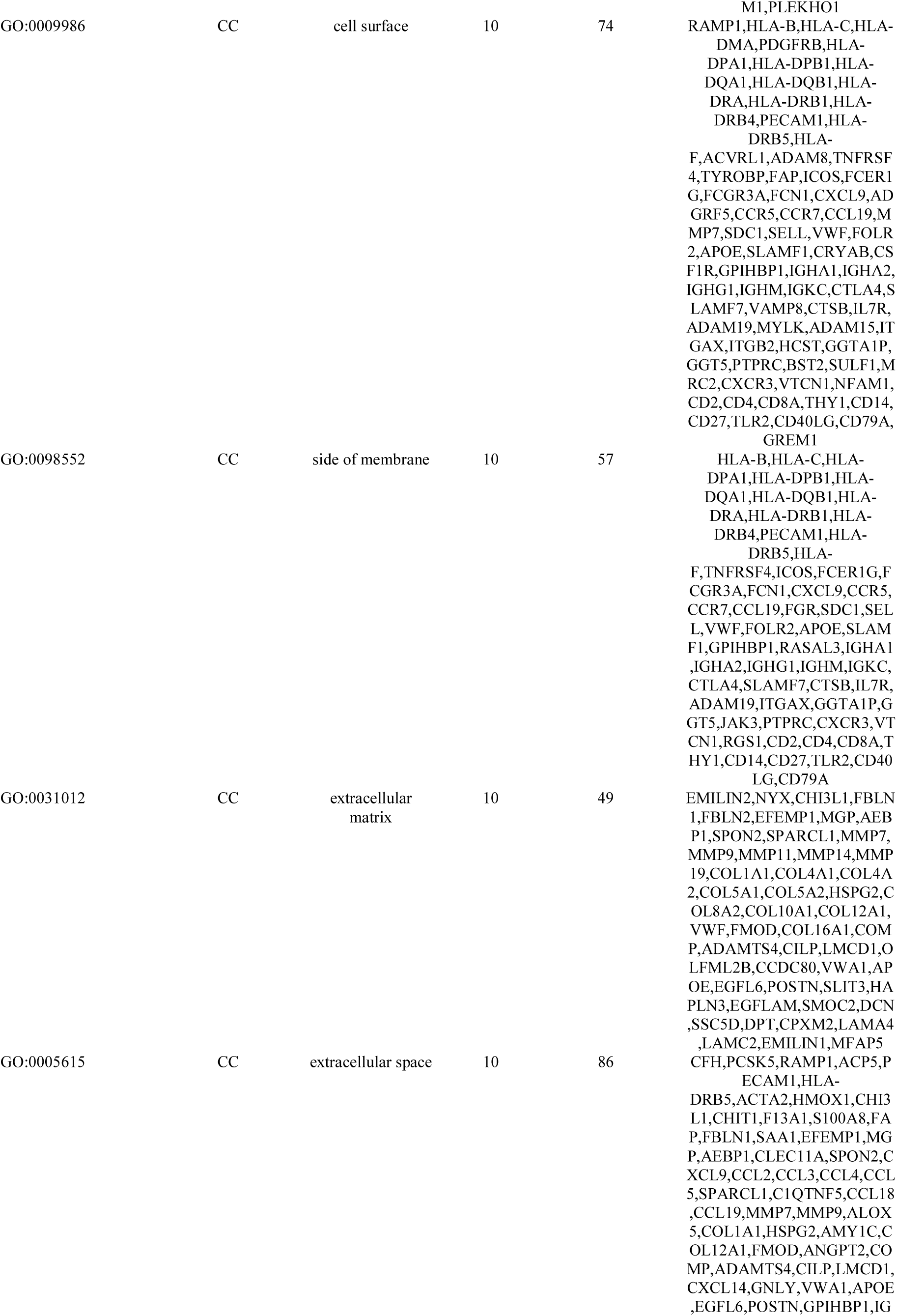
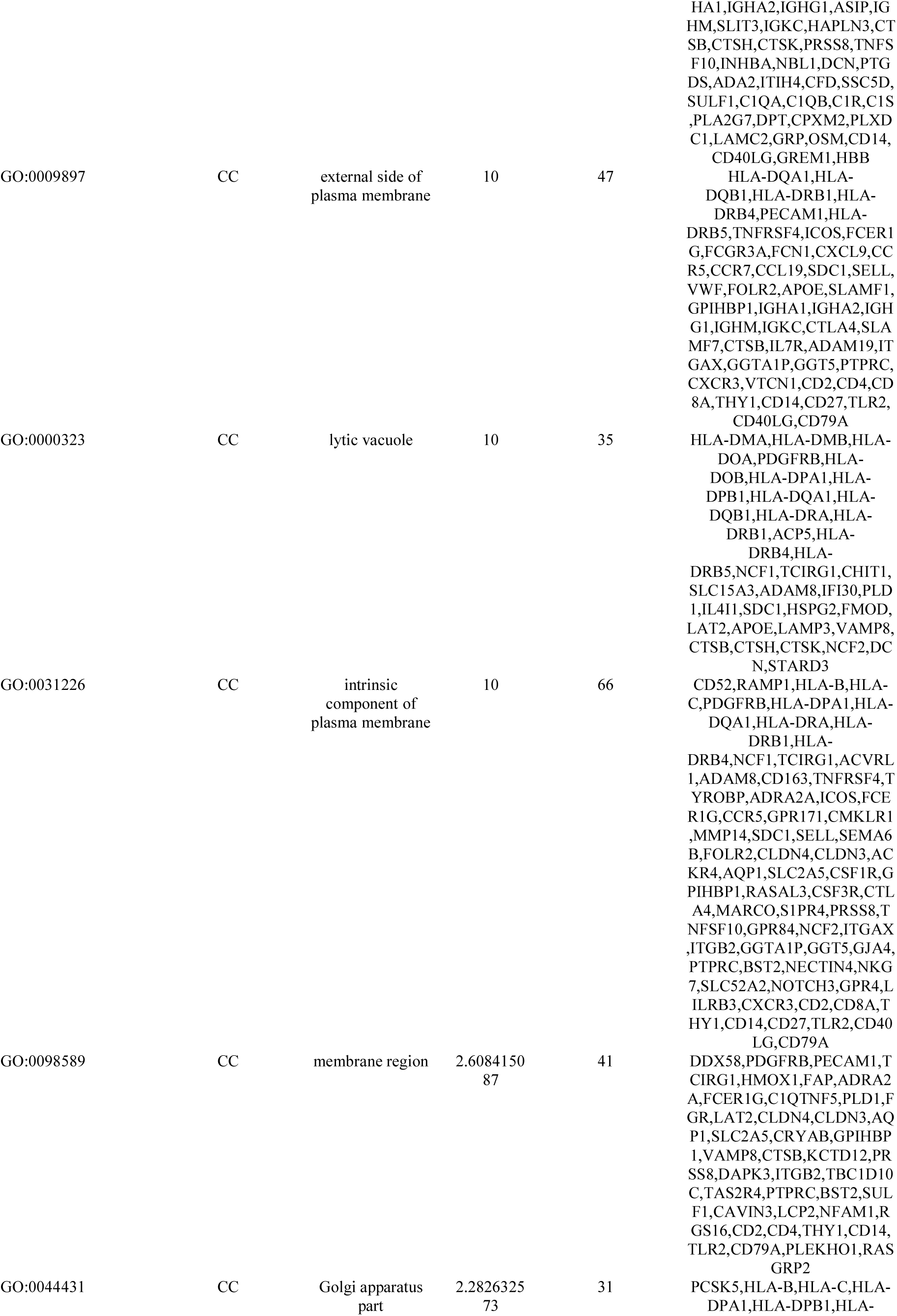
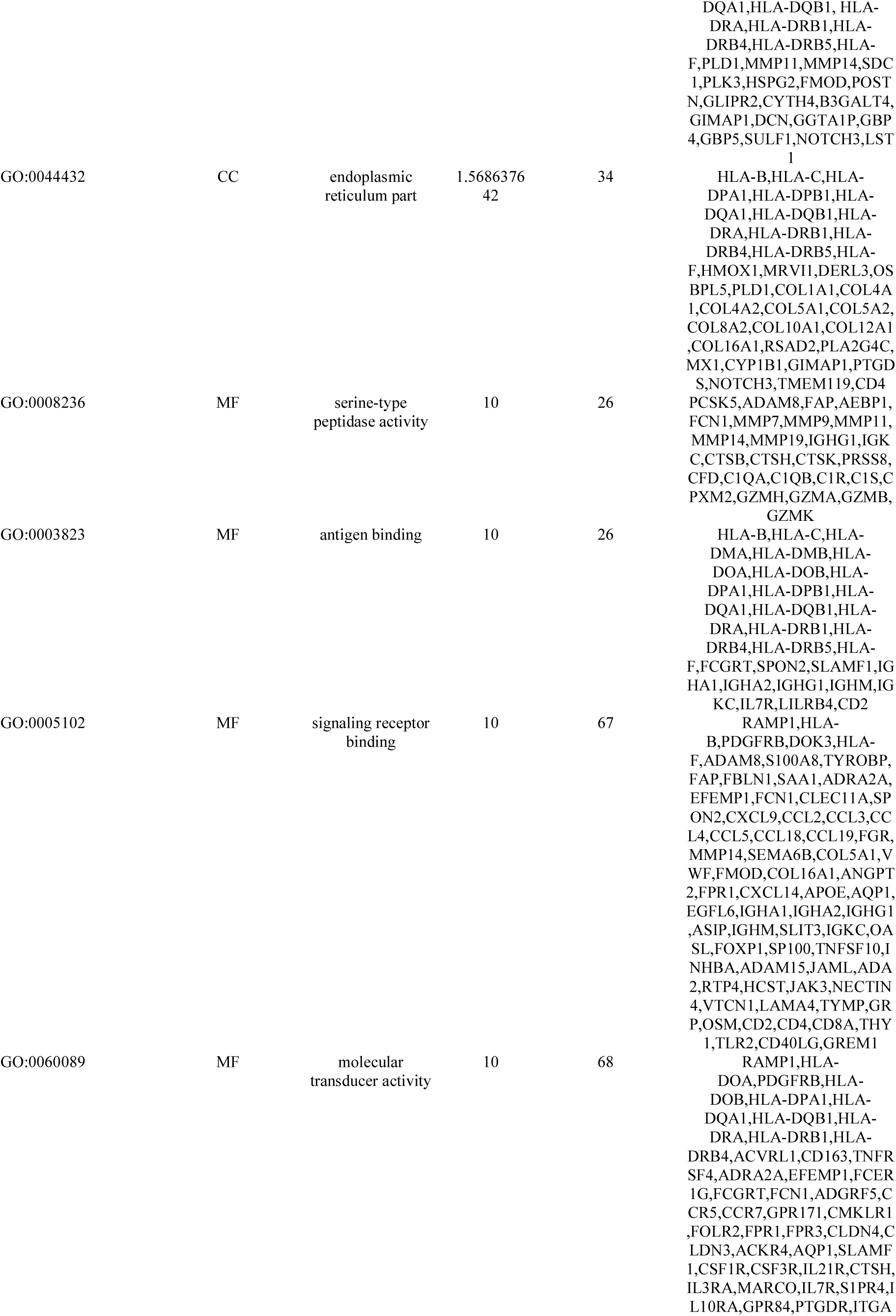
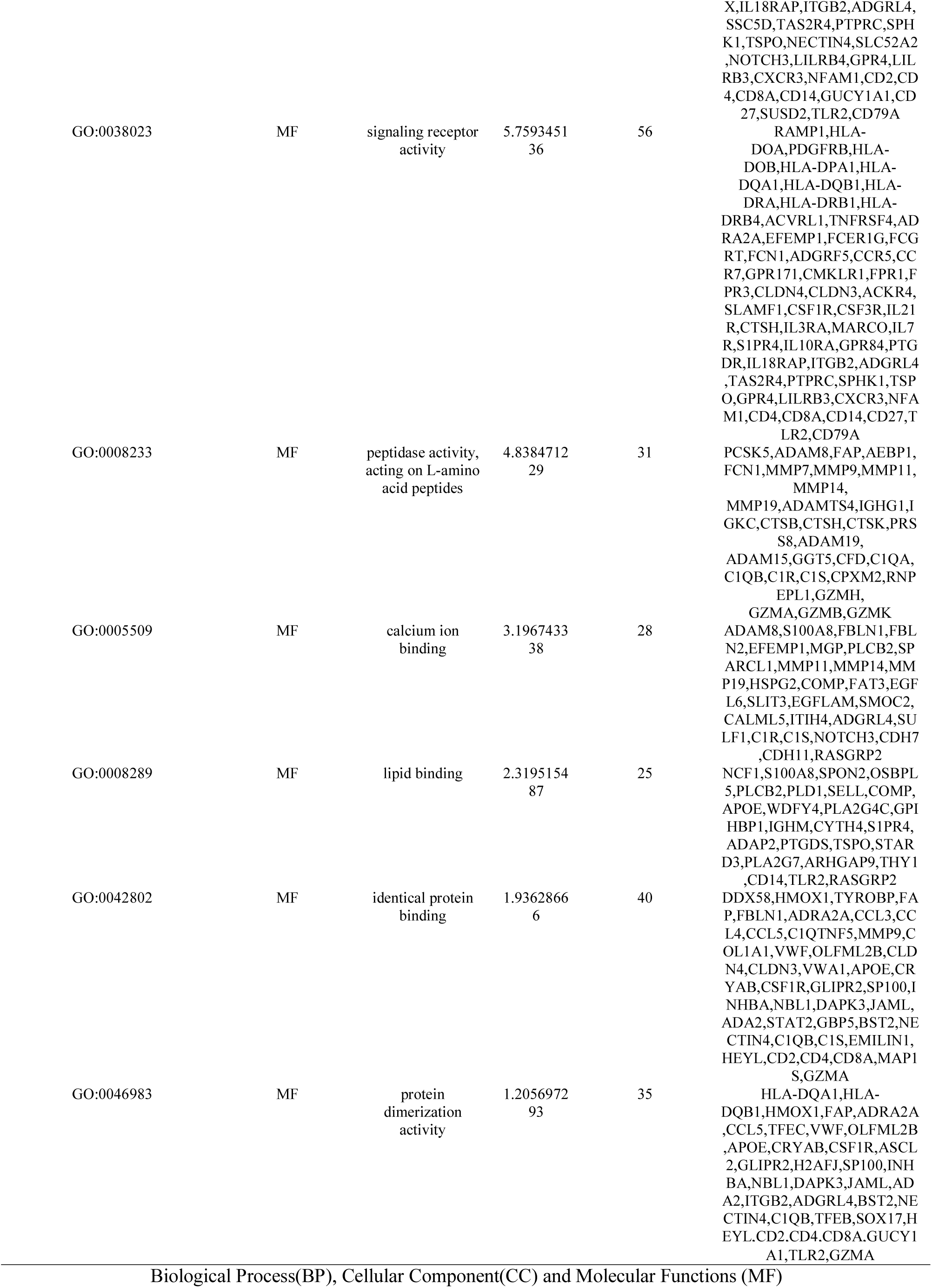
The enriched GO terms of the up regulated differentially expressed genes

**Table 6.**
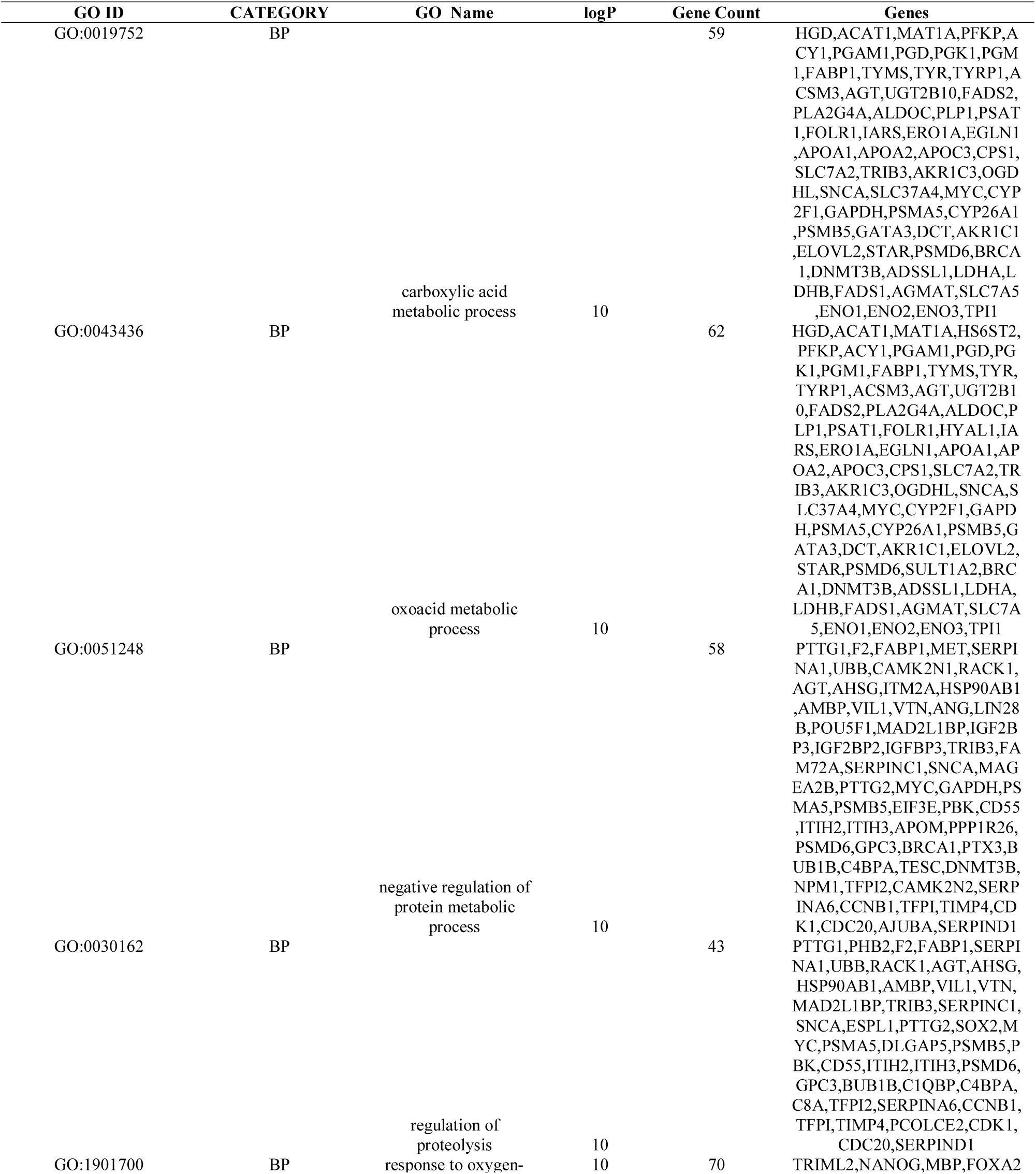
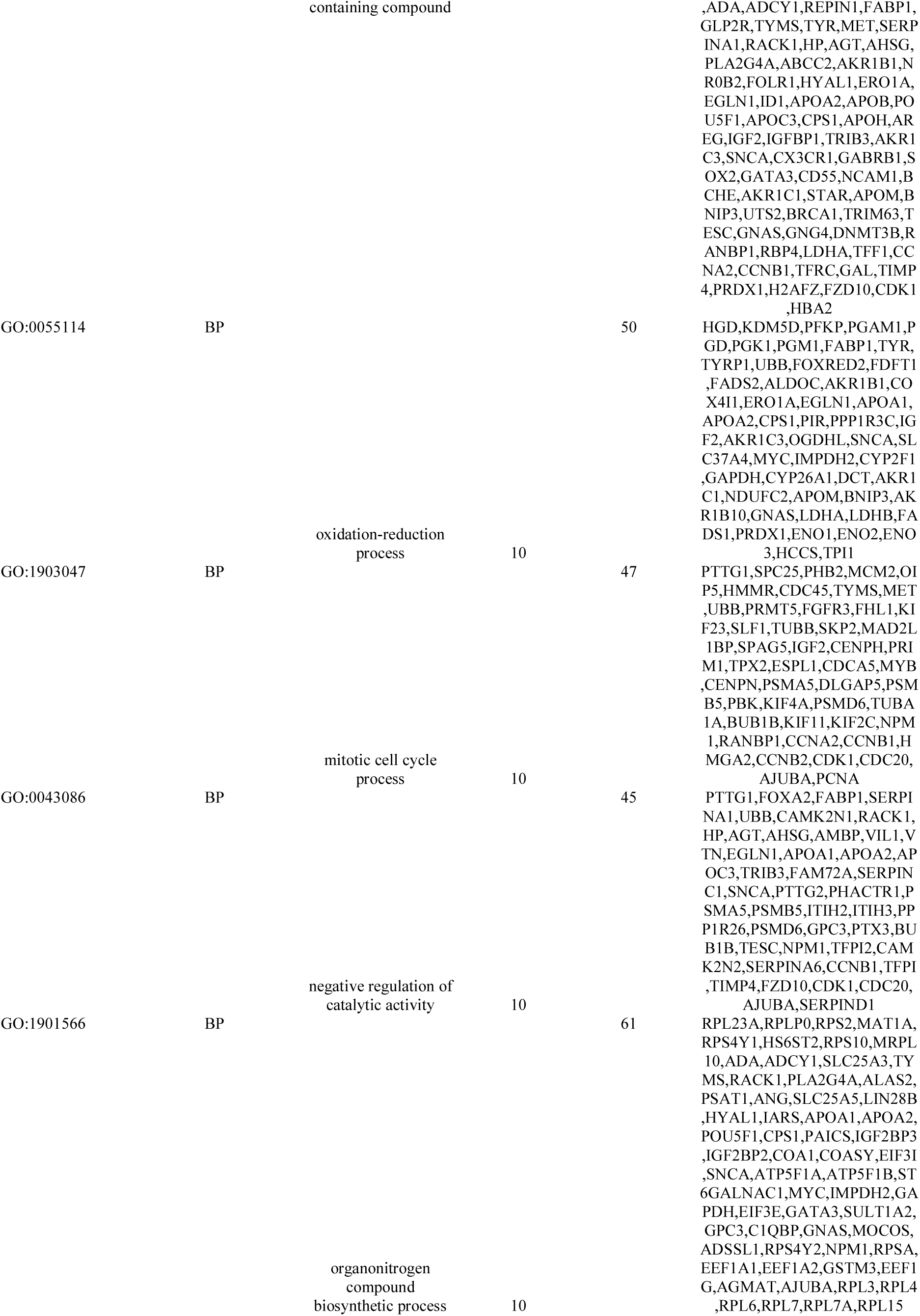
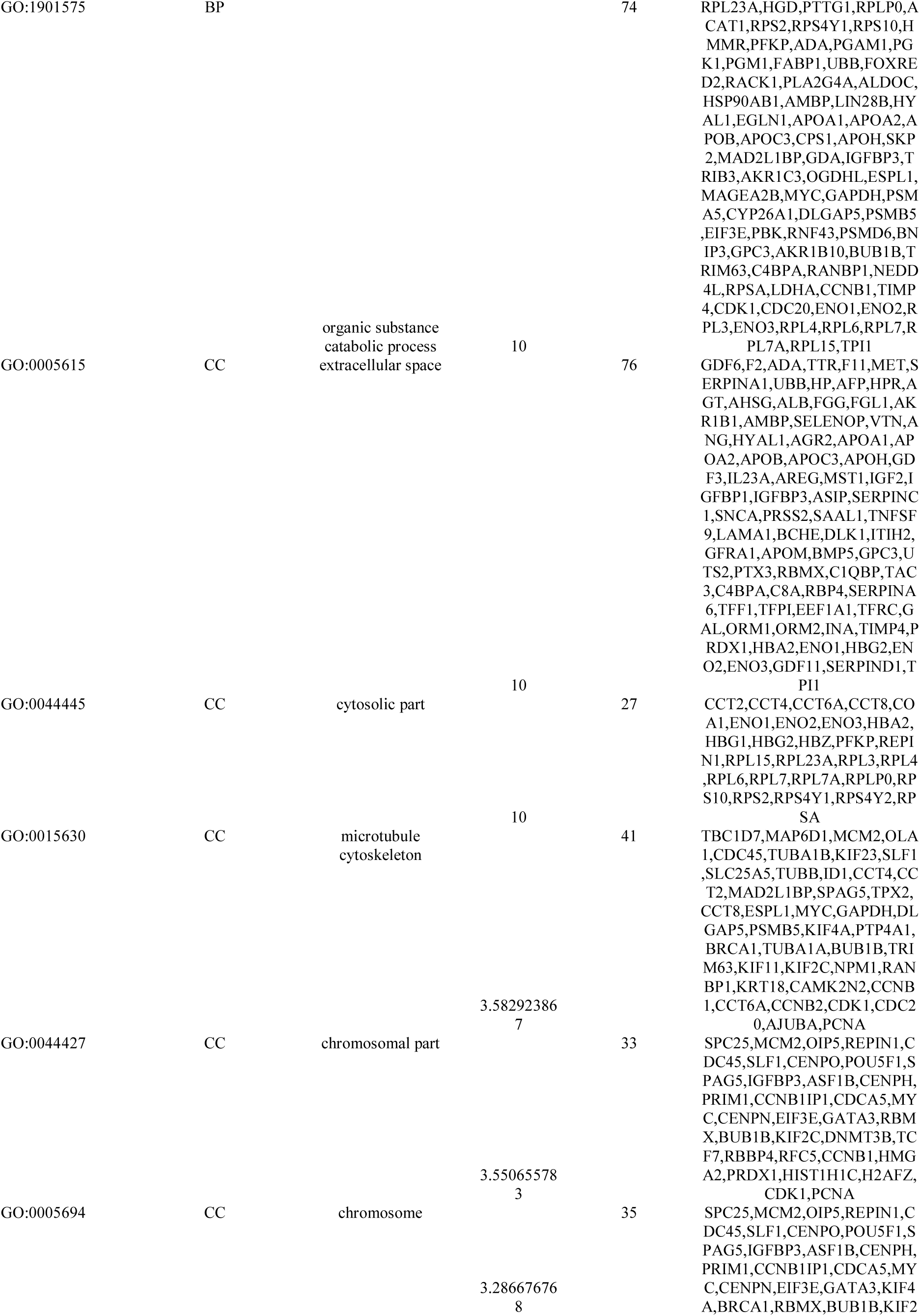
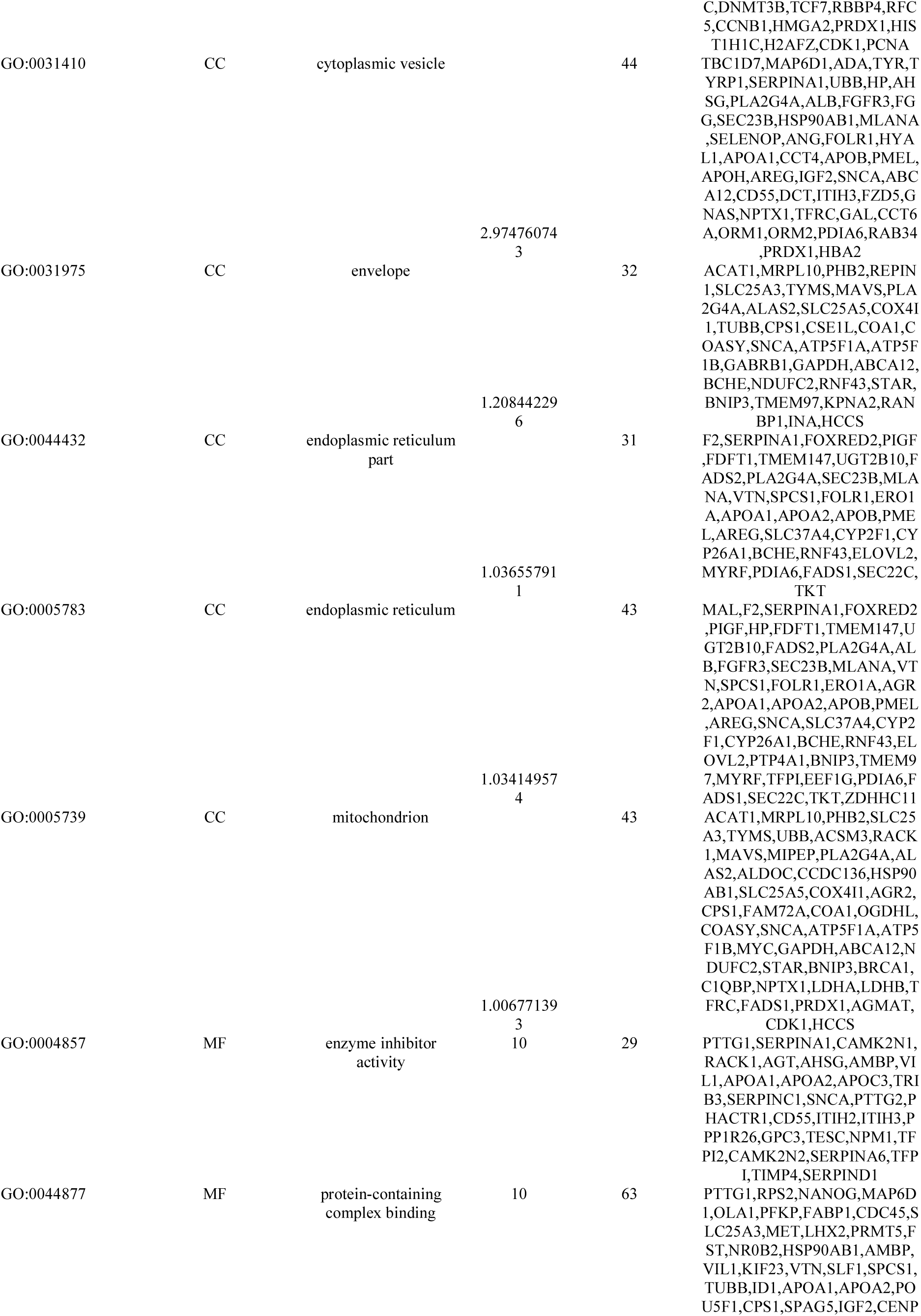
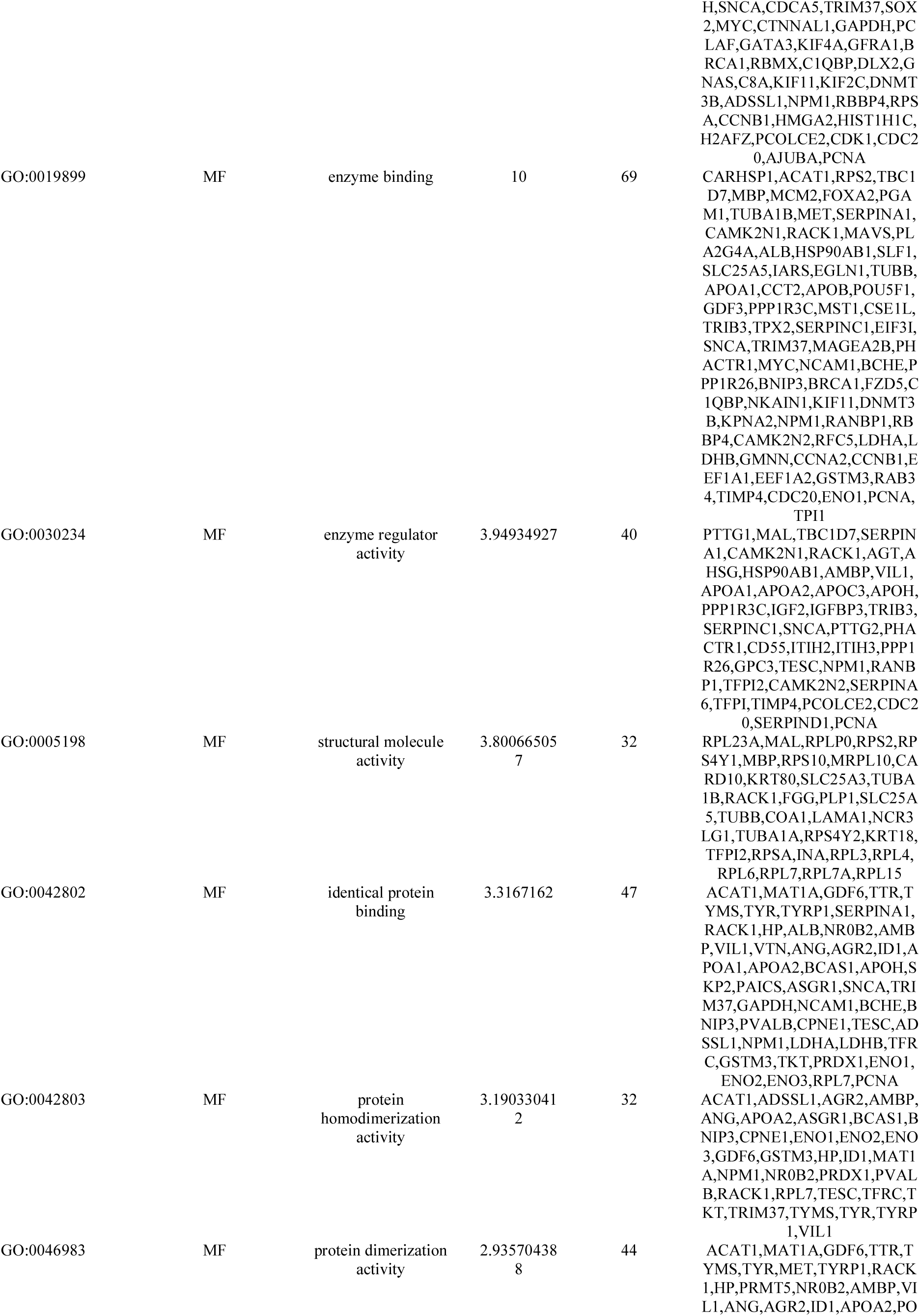
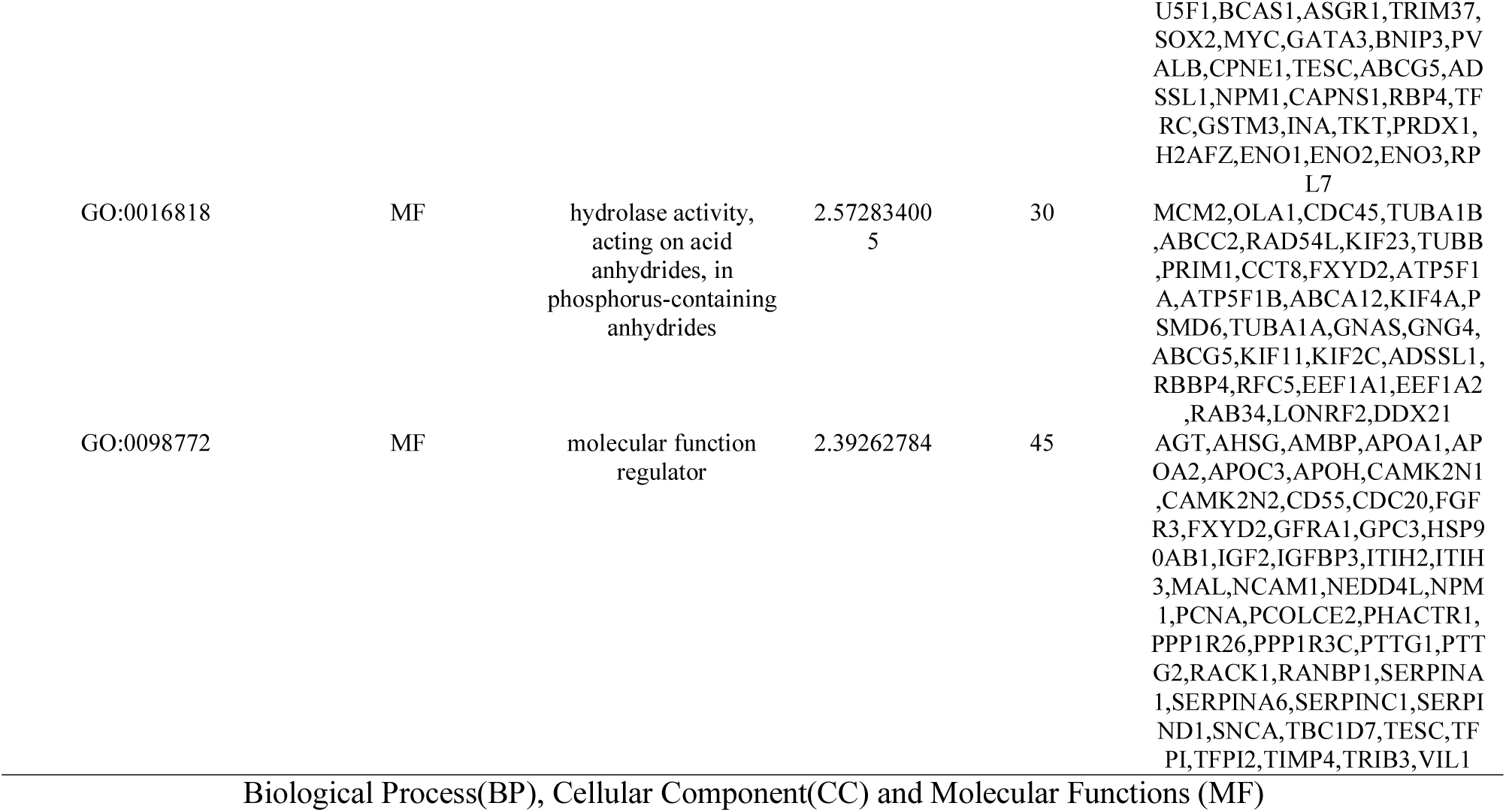
The enriched GO terms of the down regulated differentially expressed genes

### PPI network construction and module analysis

The InnateDb interactome database and Cytoscape were used to construct a PPI network of the potential interactions between the up and down regulated genes. As presented in Fig. 5, there were 2762 nodes and 4473 interactions found in the PPI network for up regulated genes. Hub genes with high node degree distribution, betweenness centrality, stress centrality, closeness centrality such as UBD, HLA-B, IL7R, CD4, PDGFRB, IRF8 and STAT2, and low clustering coefficient such as DERL3, IGFLR1, RAMP1, MARCO and ICOS are listed in Table 7. The statistical results and scatter plot for node degree distribution, betweenness centrality, stress centrality, closeness centrality and clustring coefficient for up regulated genes are shown in Fig. 6A - 6E. Pathway and GO enrichment analysis displayed that the top up regulated hub genes were primarily enriched in the immune response, antigen processing and presentation, PI3K-Akt signaling pathway, regulation of immune system process, HTLV-I infection, cytokine signaling in immune system, herpes simplex infection, endoplasmic reticulum part, cell surface, intrinsic component of plasma membrane and cell adhesion molecules (CAMs). Similarly, as presented in Fig. 7, there were 4710 nodes and 12360 interactions found in the PPI network for down regulated genes. Hub genes with high node degree such as high node degree distribution, betweenness centrality, stress centrality, closeness centrality such as MYC, HSP90AB1, BRCA1, EEF1A1, HNRNPA1, PCNA, RPSA, GAPDH, and low clustering coefficient such as C8A, GDF11, SERPINA1, TFRC and MET are listed in Table 7. The statistical results and scatter plot for node degree distribution, betweenness centrality, stress centrality, closeness centrality and clustring coefficient for down regulated genes are shown in Fig. 8A - 8E. Pathway and GO enrichment analysis displayed that the top down regulated hub genes were primarily enriched in the cell cycle, pathways in cancer, ubiquitin mediated proteolysis, peptide chain elongation, mitotic cell cycle process, ribosome, biosynthesis of amino acids, complement and coagulation cascades, ensemble of genes encoding extracellular matrix and extracellular matrix-associated proteins, negative regulation of protein metabolic process, FOXA2 and FOXA3 transcription factor networks and response to oxygen-containing compound.

**Fig. 5.**
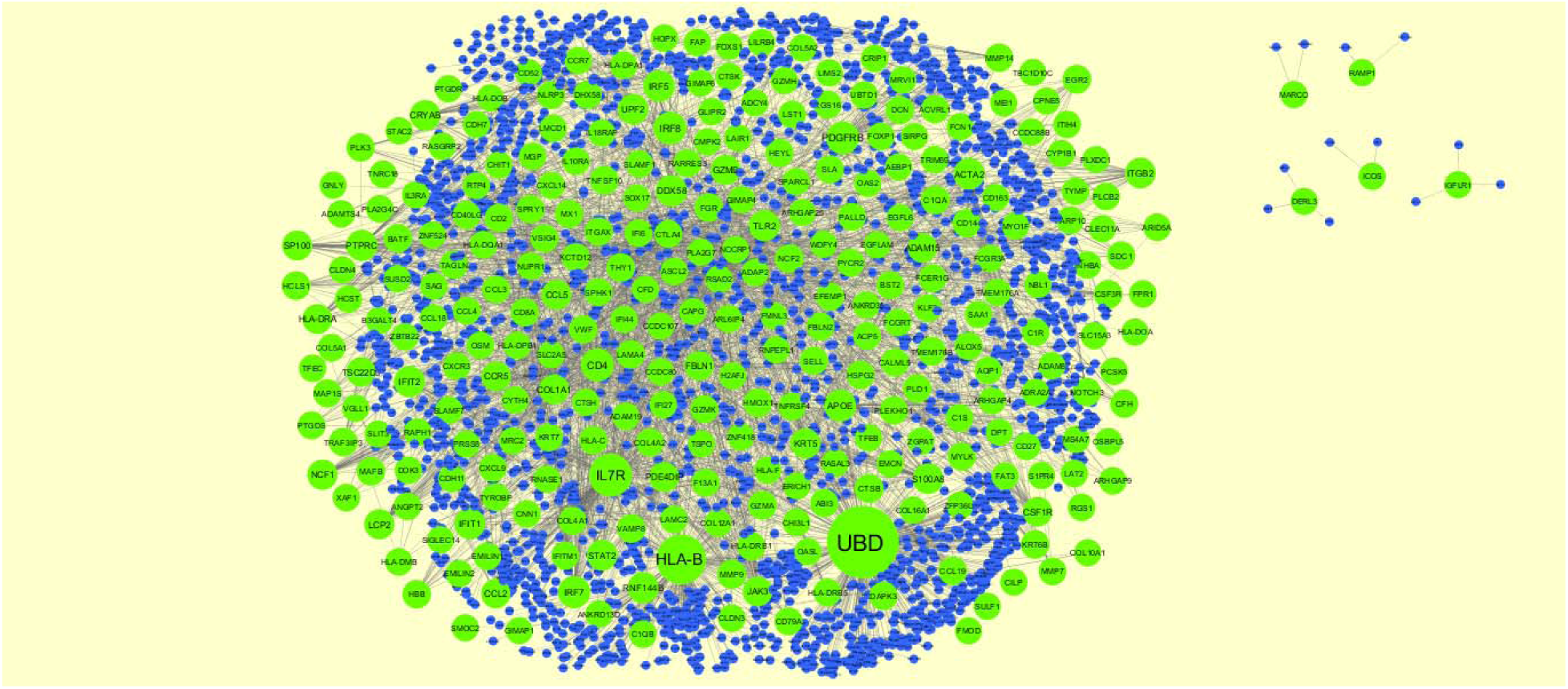
Protein–protein interaction network of up regulated genes. Green nodes denotes up regulated genes.

**Fig. 6.**
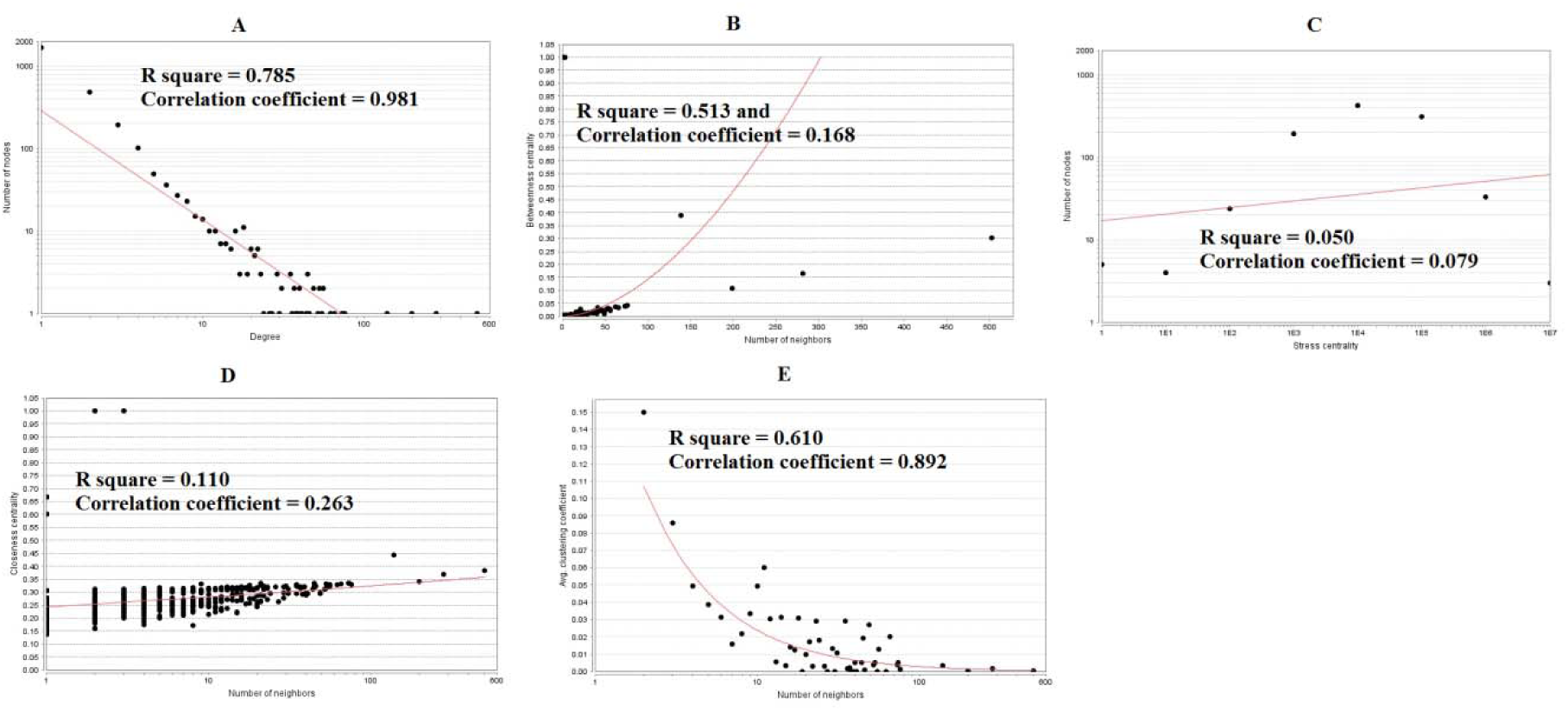
Scatter plot for up regulated genes. (A- Node degree; B- Betweenness centrality; C- Stress centrality; D- Closeness centrality; E- Clustering coefficient)

**Fig. 7.**
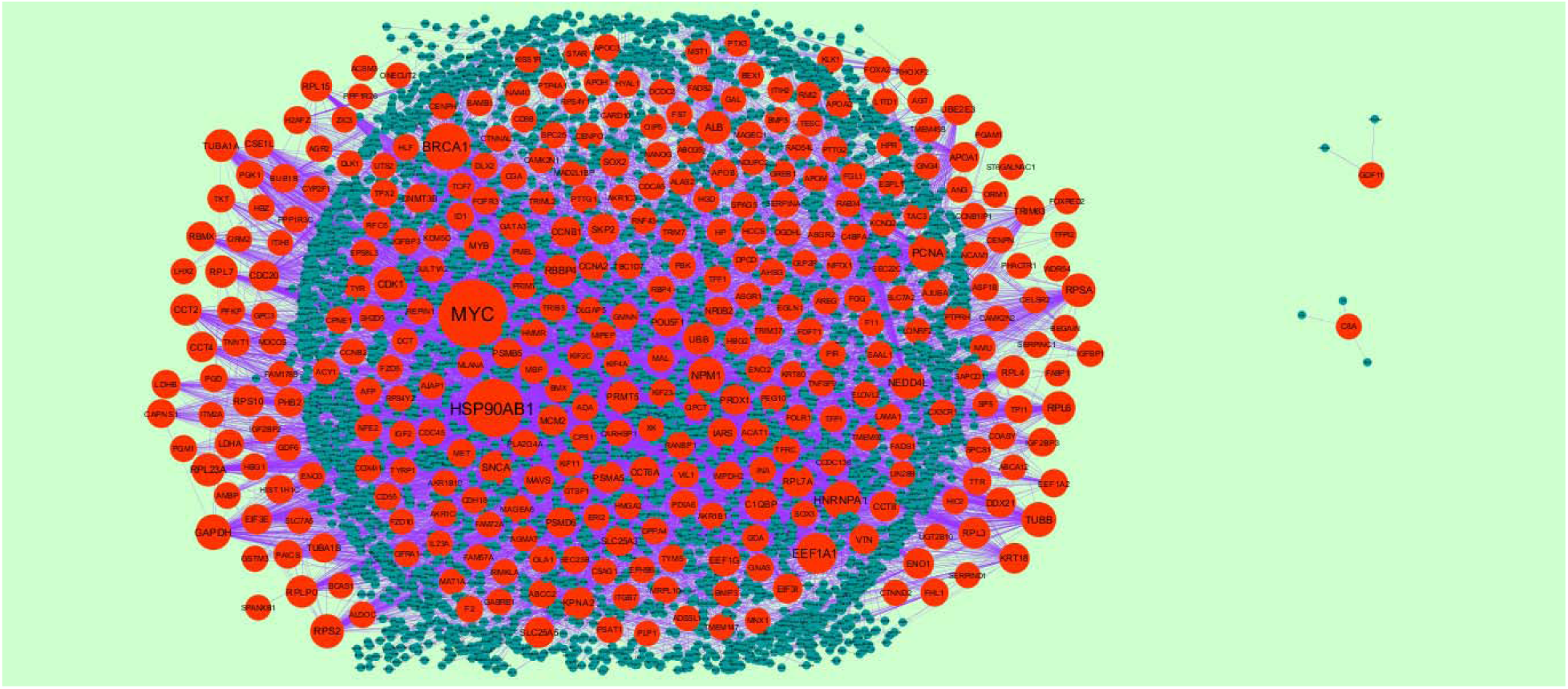
Protein–protein interaction network of down regulated genes. Red nodes denotes down regulated genes.

**Fig. 8.**
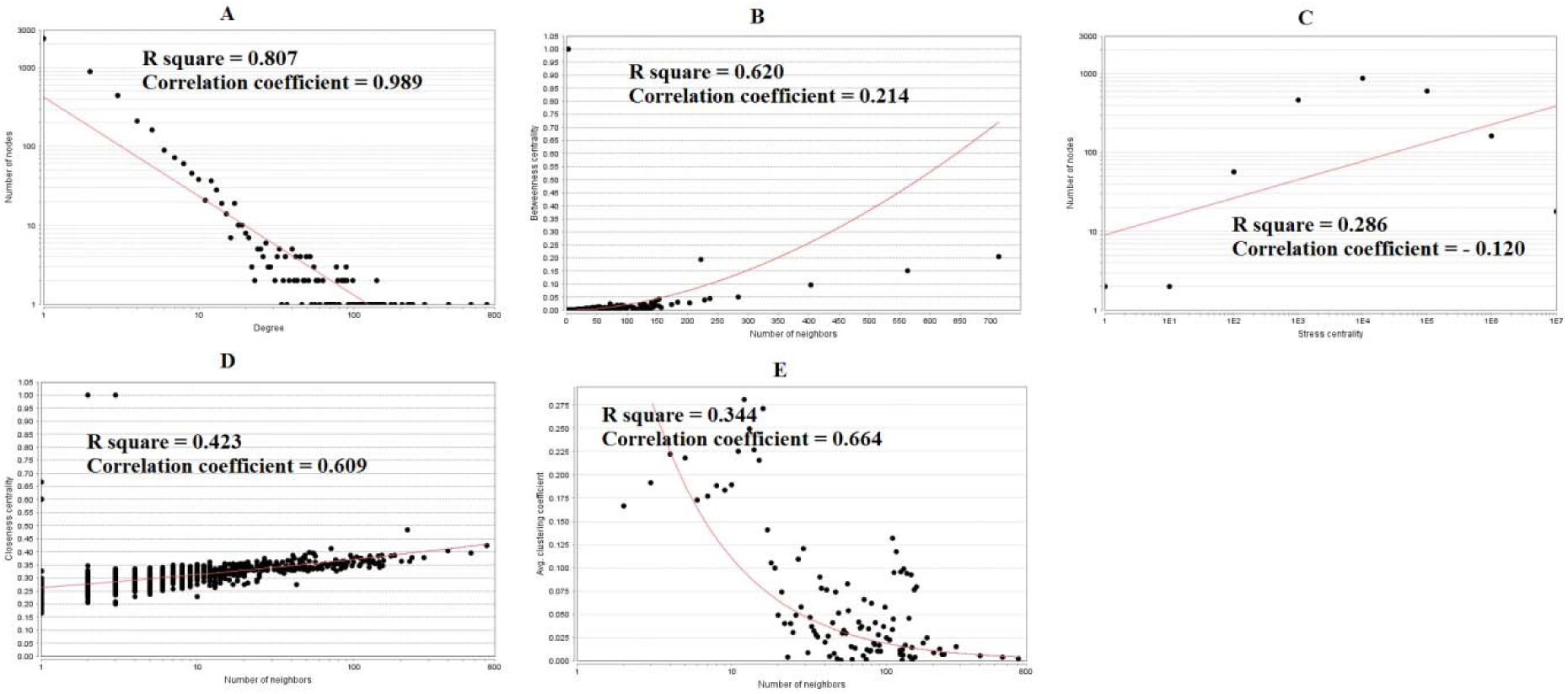
Scatter plot for down regulated genes. (A- Node degree; B- Betweenness centrality; C- Stress centrality; D- Closeness centrality; E- Clustering coefficient)

**Table 7.** Topology table for up and down regulated genes

| Regulation | Node | Degree | Betweenness | Stress | Closeness | Clustering Coefficient |
| --- | --- | --- | --- | --- | --- | --- |
| Up | UBD | 502 | 0.302275 | 16950660 | 0.384045 | 4.61E-04 |
| Up | HLA-B | 281 | 0.164738 | 9465596 | 0.368372 | 0.001525 |
| Up | IL7R | 199 | 0.107848 | 14949930 | 0.341804 | 4.57E-04 |
| Up | CD4 | 76 | 0.042133 | 3911440 | 0.330125 | 0.001404 |
| Up | PDGFRB | 74 | 0.040256 | 2681424 | 0.334145 | 0.005183 |
| Up | IRF8 | 73 | 0.038659 | 2565812 | 0.335945 | 0.003805 |
| Up | IRF7 | 66 | 0.0331 | 1989330 | 0.331681 | 0.020047 |
| Up | APOE | 62 | 0.035788 | 2025952 | 0.327798 | 0 |
| Up | DDX58 | 56 | 0.024544 | 2061120 | 0.324043 | 6.49E-04 |
| Up | IRF5 | 56 | 0.023574 | 1548406 | 0.330284 | 0.025325 |
| Up | ACTA2 | 55 | 0.029533 | 2038742 | 0.325852 | 0 |
| Up | PTPRC | 53 | 0.027721 | 1213282 | 0.332929 | 0.01016 |
| Up | RNF144B | 53 | 0.030673 | 1603846 | 0.315439 | 0 |
| Up | TLR2 | 52 | 0.029312 | 2006014 | 0.312173 | 0.003771 |
| Up | IFIT1 | 49 | 0.013024 | 1408454 | 0.296681 | 0.027211 |
| Up | IFIT2 | 49 | 0.009994 | 1218310 | 0.294959 | 0.026361 |
| Up | CRYAB | 46 | 0.02525 | 1470644 | 0.32328 | 9.66E-04 |
| Up | S100A8 | 45 | 0.021005 | 1360014 | 0.322217 | 0.00303 |
| Up | ITGB2 | 45 | 0.022099 | 1371900 | 0.318477 | 0.00303 |
| Up | STAT2 | 45 | 0.021537 | 1232394 | 0.334838 | 0.051515 |
| Up | UPF2 | 44 | 0.022875 | 1150530 | 0.313028 | 0.005285 |
| Up | LCP2 | 40 | 0.012169 | 2067080 | 0.288175 | 0 |
| Up | NCF1 | 40 | 0.015369 | 661004 | 0.298002 | 0.010256 |
| Up | KRT5 | 39 | 0.014715 | 1367672 | 0.321575 | 0 |
| Up | GZMB | 38 | 0.01355 | 602118 | 0.290925 | 0 |
| Up | PDE4DIP | 37 | 0.019054 | 1056154 | 0.315729 | 0 |
| Up | SP100 | 37 | 0.017696 | 646020 | 0.320261 | 0.004505 |
| Up | COL1A1 | 36 | 0.01797 | 1077814 | 0.32165 | 0.001587 |
| Up | HLA-DRA | 35 | 0.01681 | 871340 | 0.322634 | 0.045378 |
| Up | CCL5 | 35 | 0.013092 | 726240 | 0.29499 | 0.030252 |
| Up | JAK3 | 35 | 0.013511 | 614458 | 0.329056 | 0.011765 |
| Up | ADAM15 | 31 | 0.012973 | 902560 | 0.313314 | 0 |
| Up | CCR5 | 31 | 0.014936 | 419342 | 0.295212 | 0.021505 |
| Up | CSF1R | 30 | 0.011451 | 1285278 | 0.323738 | 0 |
| Up | CCL2 | 29 | 0.009028 | 456566 | 0.298943 | 0.039409 |
| Up | FBLN1 | 29 | 0.011988 | 450122 | 0.27706 | 0 |
| Up | TSC22D3 | 29 | 0.01097 | 1029688 | 0.320149 | 0 |
| Up | MMP9 | 27 | 0.01006 | 1963768 | 0.263289 | 0 |
| Up | CTSB | 26 | 0.01367 | 733408 | 0.320823 | 0.003077 |
| Up | TNFSF10 | 24 | 0.006552 | 280408 | 0.295562 | 0.018116 |
| Up | NCF2 | 23 | 0.015009 | 561080 | 0.319255 | 0.055336 |
| Up | C1QA | 23 | 0.010035 | 510722 | 0.271093 | 0.027668 |
| Up | EGR2 | 23 | 0.011552 | 535974 | 0.313887 | 0.003953 |
| Up | FGR | 22 | 0.009047 | 727656 | 0.32199 | 0.008658 |
| Up | HMOX1 | 22 | 0.008782 | 728174 | 0.318145 | 0 |
| Up | THY1 | 22 | 0.008748 | 501966 | 0.31157 | 0 |
| Up | VAMP8 | 22 | 0.011723 | 568706 | 0.312849 | 0 |
| Up | TRAF3IP3 | 22 | 0.011978 | 547354 | 0.311889 | 0 |
| Up | HCLS1 | 22 | 0.006302 | 236634 | 0.28655 | 0.008658 |
| Up | LAMA4 | 21 | 0.00523 | 726394 | 0.266072 | 0 |
| Up | KCTD12 | 21 | 0.009889 | 427674 | 0.327173 | 0.004762 |
| Up | GZMA | 21 | 0.006168 | 895510 | 0.263416 | 0 |
| Up | MAFB | 20 | 0.009604 | 296158 | 0.263669 | 0.021053 |
| Up | CSF3R | 20 | 0.008479 | 713904 | 0.317225 | 0 |
| Up | HBB | 20 | 0.006616 | 387328 | 0.313779 | 0.026316 |
| Up | BATF | 20 | 0.007018 | 778768 | 0.244476 | 0 |
| Up | CD14 | 20 | 0.012001 | 854950 | 0.251974 | 0.005263 |
| Up | KRT6B | 19 | 0.005782 | 496980 | 0.311288 | 0 |
| Up | MMP14 | 19 | 0.011862 | 460210 | 0.309392 | 0 |
| Up | SPRY1 | 19 | 0.004879 | 621564 | 0.274977 | 0 |
| Up | HLA-C | 18 | 0.006045 | 356646 | 0.314607 | 0.013072 |
| Up | CCL19 | 18 | 0.00516 | 306414 | 0.323967 | 0.130719 |
| Up | CD2 | 18 | 0.00815 | 407412 | 0.315947 | 0.019608 |
| Up | TAGLN | 18 | 0.007256 | 390648 | 0.309323 | 0 |
| Up | HLA-DRB1 | 18 | 0.004263 | 508188 | 0.315765 | 0.091503 |
| Up | CD8A | 18 | 0.008187 | 250414 | 0.327603 | 0.039216 |
| Up | DAPK3 | 18 | 0.005695 | 515936 | 0.31562 | 0 |
| Up | HSPG2 | 18 | 0.005625 | 647004 | 0.257943 | 0.026144 |
| Up | NOTCH3 | 18 | 0.011172 | 384008 | 0.315947 | 0 |
| Up | PLD1 | 18 | 0.006783 | 350082 | 0.309776 | 0 |
| Up | DHX58 | 17 | 0.016559 | 484556 | 0.314535 | 0.014706 |
| Up | VWF | 17 | 0.00802 | 288034 | 0.318551 | 0.007353 |
| Up | DCN | 17 | 0.007617 | 400960 | 0.255565 | 0.014706 |
| Up | COL4A1 | 16 | 0.005724 | 315884 | 0.316494 | 0.025 |
| Up | ITGAX | 16 | 0.003956 | 195814 | 0.269178 | 0.025 |
| Up | HEYL | 16 | 0.007673 | 481602 | 0.316238 | 0 |
| Up | SLA | 16 | 0.00585 | 357650 | 0.318773 | 0.033333 |
| Up | MX1 | 16 | 0.007186 | 401100 | 0.319255 | 0.016667 |
| Up | PLK3 | 16 | 0.006652 | 279404 | 0.316238 | 0.008333 |
| Up | SDC1 | 16 | 0.009469 | 410180 | 0.319963 | 0.008333 |
| Up | TCIRG1 | 16 | 0.006859 | 359880 | 0.309602 | 0 |
| Up | CFH | 15 | 0.00445 | 244882 | 0.295117 | 0.009524 |
| Up | ABI3 | 15 | 0.007402 | 731014 | 0.219889 | 0 |
| Up | SELL | 15 | 0.007166 | 346972 | 0.316567 | 0.009524 |
| Up | ALOX5 | 15 | 0.007838 | 873370 | 0.267316 | 0 |
| Up | ACVRL1 | 15 | 0.008005 | 1719772 | 0.223235 | 0 |
| Up | MAP1S | 15 | 0.005764 | 334716 | 0.309288 | 0 |
| Up | PLEKHO1 | 14 | 0.006138 | 280456 | 0.309288 | 0 |
| Up | MYLK | 14 | 0.005342 | 544002 | 0.315693 | 0 |
| Up | EFEMP1 | 14 | 0.004205 | 276612 | 0.311995 | 0 |
| Up | COL4A2 | 14 | 0.002731 | 112746 | 0.270318 | 0.032967 |
| Up | CD40LG | 14 | 0.005908 | 269538 | 0.318366 | 0.010989 |
| Up | FCER1G | 13 | 0.005848 | 305260 | 0.31504 | 0 |
| Up | PALLD | 13 | 0.005071 | 298560 | 0.308939 | 0 |
| Up | RARRES3 | 13 | 0.00331 | 406516 | 0.313493 | 0 |
| Up | RGS16 | 13 | 0.004116 | 729950 | 0.23737 | 0 |
| Up | SPHK1 | 13 | 0.004949 | 342988 | 0.320261 | 0.012821 |
| Up | CTLA4 | 13 | 0.002854 | 347870 | 0.271119 | 0 |
| Up | F13A1 | 13 | 0.003164 | 212990 | 0.272682 | 0.025641 |
| Up | NLRP3 | 12 | 0.004585 | 826154 | 0.232582 | 0 |
| Up | PTGDS | 12 | 0.004022 | 505956 | 0.231953 | 0 |
| Up | CD79A | 12 | 0.004848 | 216848 | 0.317519 | 0.030303 |
| Up | CALML5 | 12 | 0.004267 | 221012 | 0.310162 | 0 |
| Up | FBLN2 | 12 | 0.00218 | 194722 | 0.23836 | 0.060606 |
| Up | TYROBP | 12 | 0.006573 | 767160 | 0.228876 | 0 |
| Up | CXCR3 | 11 | 0.00241 | 88198 | 0.239944 | 0.018182 |
| Up | HLA-DQA1 | 11 | 0.003891 | 239548 | 0.310021 | 0 |
| Up | CCL3 | 11 | 5.10E-04 | 52576 | 0.236531 | 0.163636 |
| Up | PYCR2 | 11 | 0.00456 | 302658 | 0.3136 | 0 |
| Up | C1S | 11 | 0.003338 | 264818 | 0.244085 | 0.072727 |
| Up | EMILIN1 | 11 | 0.005448 | 372612 | 0.223617 | 0 |
| Up | CCL4 | 10 | 8.16E-04 | 80812 | 0.247185 | 0.2 |
| Up | MMP7 | 10 | 0.003914 | 319190 | 0.213308 | 0 |
| Up | KRT7 | 10 | 0.004334 | 239098 | 0.312635 | 0 |
| Up | KLF2 | 10 | 0.004907 | 265400 | 0.312422 | 0 |
| Up | GZMK | 10 | 0.001459 | 241170 | 0.244346 | 0 |
| Up | HLA-DRB5 | 10 | 0.001276 | 222656 | 0.313779 | 0.2 |
| Up | BST2 | 10 | 0.00352 | 149770 | 0.308905 | 0 |
| Up | IFITM1 | 10 | 0.002258 | 145372 | 0.315838 | 0.111111 |
| Up | FOXP1 | 10 | 0.003912 | 432154 | 0.254924 | 0 |
| Up | ACP5 | 10 | 0.005474 | 165884 | 0.312493 | 0.022222 |
| Up | C1QB | 9 | 0.001965 | 127342 | 0.250685 | 0.111111 |
| Up | LAMC2 | 9 | 0.003181 | 154346 | 0.309183 | 0 |
| Up | CXCL9 | 9 | 0.00301 | 229902 | 0.263289 | 0 |
| Up | ARID5A | 9 | 0.005318 | 173402 | 0.330842 | 0.027778 |
| Up | NUPR1 | 9 | 0.002524 | 298524 | 0.246585 | 0 |
| Up | CCL18 | 8 | 0.001778 | 235150 | 0.243998 | 0 |
| Up | PARP10 | 8 | 0.002967 | 169404 | 0.309079 | 0 |
| Up | CNN1 | 8 | 0.002758 | 130418 | 0.309253 | 0 |
| Up | CHI3L1 | 8 | 6.77E-04 | 76426 | 0.25 | 0 |
| Up | PLCB2 | 8 | 0.002583 | 109656 | 0.309288 | 0 |
| Up | INHBA | 8 | 0.002491 | 346390 | 0.172124 | 0 |
| Up | ANGPT2 | 8 | 0.001319 | 89508 | 0.310197 | 0 |
| Up | ARL6IP4 | 8 | 0.002341 | 184198 | 0.310724 | 0 |
| Up | IL21R | 8 | 0.001037 | 140942 | 0.244934 | 0 |
| Up | ASCL2 | 8 | 0.001824 | 174586 | 0.260119 | 0 |
| Up | IFI6 | 8 | 0.001077 | 201080 | 0.301572 | 0.178571 |
| Up | PLA2G4C | 8 | 9.57E-04 | 110798 | 0.256737 | 0 |
| Up | CLDN3 | 8 | 0.004348 | 386516 | 0.317152 | 0 |
| Up | HOPX | 8 | 0.00371 | 810748 | 0.221935 | 0 |
| Up | CD27 | 8 | 0.003403 | 725650 | 0.232286 | 0 |
| Up | CTSK | 7 | 0.002827 | 418038 | 0.212072 | 0 |
| Up | AQP1 | 7 | 0.002949 | 207146 | 0.21137 | 0 |
| Up | COL5A1 | 7 | 0.005883 | 223714 | 0.308592 | 0.047619 |
| Up | ADAMTS4 | 7 | 0.002492 | 142124 | 0.309288 | 0 |
| Up | RAPH1 | 7 | 0.002642 | 158114 | 0.3088 | 0 |
| Up | LST1 | 7 | 0.001581 | 36596 | 0.259603 | 0 |
| Up | VSIG4 | 7 | 0.001821 | 226500 | 0.243543 | 0 |
| Up | ARHGAP9 | 7 | 0.002225 | 279174 | 0.313815 | 0 |
| Up | XAF1 | 7 | 0.002586 | 312004 | 0.271253 | 0 |
| Up | OSM | 7 | 0.003147 | 352934 | 0.236858 | 0 |
| Up | RSAD2 | 7 | 2.78E-04 | 34162 | 0.259186 | 0 |
| Up | SLAMF1 | 7 | 0.001708 | 53706 | 0.263922 | 0 |
| Up | TNFRSF4 | 7 | 0.001983 | 164224 | 0.21882 | 0 |
| Up | ITIH4 | 7 | 0.001022 | 122020 | 0.25509 | 0 |
| Up | HLA-DPA1 | 6 | 0.001024 | 54710 | 0.307073 | 0 |
| Up | HLA-DPB1 | 6 | 9.69E-04 | 111958 | 0.230608 | 0 |
| Up | HLA-DMB | 6 | 0.003785 | 69950 | 0.265506 | 0.133333 |
| Up | LAT2 | 6 | 0.002505 | 169848 | 0.31097 | 0 |
| Up | IL10RA | 6 | 0.001329 | 67330 | 0.307244 | 0 |
| Up | UBTD1 | 6 | 0.003009 | 101750 | 0.307073 | 0 |
| Up | LMCD1 | 6 | 0.0023 | 312490 | 0.23403 | 0 |
| Up | RGS1 | 6 | 0.002401 | 302054 | 0.22947 | 0 |
| Up | ADRA2A | 6 | 0.002926 | 126184 | 0.212746 | 0 |
| Up | TYMP | 6 | 0.002067 | 209918 | 0.312173 | 0 |
| Up | ADAP2 | 6 | 3.68E-04 | 36340 | 0.252554 | 0 |
| Up | C1R | 6 | 9.13E-04 | 64610 | 0.248528 | 0.266667 |
| Up | SPARCL1 | 6 | 0.00209 | 117824 | 0.311041 | 0 |
| Up | TRIM69 | 6 | 0.001202 | 148848 | 0.232267 | 0 |
| Up | SLAMF7 | 5 | 0.002243 | 139618 | 0.2084 | 0 |
| Up | PCSK5 | 5 | 0.001951 | 113638 | 0.309148 | 0 |
| Up | CRIP1 | 5 | 0.002988 | 93620 | 0.307073 | 0 |
| Up | FCN1 | 5 | 0.00151 | 64158 | 0.201173 | 0 |
| Up | FCGRT | 5 | 9.49E-04 | 119148 | 0.261458 | 0 |
| Up | TMEM176A | 5 | 0.001809 | 192988 | 0.227642 | 0 |
| Up | COL5A2 | 5 | 0.003641 | 112224 | 0.236063 | 0.1 |
| Up | IFI27 | 5 | 2.46E-04 | 36552 | 0.314139 | 0.3 |
| Up | ADAM19 | 5 | 0.001837 | 81484 | 0.308038 | 0 |
| Up | LAIR1 | 5 | 0.001064 | 201568 | 0.314859 | 0 |
| Up | RASGRP2 | 5 | 0.00168 | 171754 | 0.312493 | 0 |
| Up | ANKRD13D | 5 | 0.001988 | 282066 | 0.316457 | 0 |
| Up | TFEC | 5 | 0.002391 | 253672 | 0.202794 | 0 |
| Up | OSBPL5 | 5 | 0.001471 | 182202 | 0.312493 | 0 |
| Up | GLIPR2 | 4 | 0.001552 | 87224 | 0.308453 | 0 |
| Up | VGLL1 | 4 | 0.002186 | 36318 | 0.173594 | 0 |
| Up | COL16A1 | 4 | 1.74E-04 | 11634 | 0.266046 | 0 |
| Up | ARHGAP4 | 4 | 0.002186 | 66450 | 0.306935 | 0 |
| Up | CDH11 | 4 | 1.14E-04 | 9298 | 0.232345 | 0 |
| Up | ZBTB22 | 4 | 0.002186 | 128490 | 0.197084 | 0 |
| Up | SAG | 4 | 0.001492 | 129474 | 0.210996 | 0 |
| Up | B3GALT4 | 4 | 0.002186 | 133788 | 0.213641 | 0 |
| Up | FAP | 4 | 7.48E-04 | 102422 | 0.210947 | 0 |
| Up | MYO1F | 4 | 5.58E-04 | 46160 | 0.307727 | 0 |
| Up | DOK3 | 4 | 1.51E-04 | 8916 | 0.2249 | 0 |
| Up | SAA1 | 4 | 0.001101 | 164528 | 0.229777 | 0 |
| Up | HLA-F | 4 | 4.18E-04 | 68162 | 0.308488 | 0 |
| Up | ADAM8 | 4 | 0.001983 | 39306 | 0.186514 | 0 |
| Up | HCST | 4 | 9.06E-04 | 82848 | 0.311217 | 0 |
| Up | OAS2 | 4 | 9.28E-04 | 194354 | 0.314607 | 0 |
| Up | FCGR3A | 4 | 0.001544 | 184974 | 0.227907 | 0 |
| Up | CYP1B1 | 4 | 5.56E-04 | 70438 | 0.308488 | 0 |
| Up | COL12A1 | 4 | 0.001193 | 72386 | 0.307485 | 0 |
| Up | FMNL3 | 4 | 0.001288 | 174096 | 0.312422 | 0 |
| Up | FPR1 | 4 | 7.79E-04 | 53162 | 0.239986 | 0 |
| Up | GZMH | 4 | 5.50E-05 | 4868 | 0.210139 | 0 |
| Up | H2AFJ | 3 | 1.16E-04 | 19544 | 0.307073 | 0 |
| Up | ZGPAT | 3 | 3.84E-05 | 3118 | 0.229355 | 0 |
| Up | RASAL3 | 3 | 1.12E-04 | 19378 | 0.237494 | 0 |
| Up | NCCRP1 | 3 | 1.57E-04 | 45650 | 0.308384 | 0 |
| Up | PLA2G7 | 3 | 3.16E-04 | 70568 | 0.31228 | 0 |
| Up | COL10A1 | 3 | 0.001457 | 166646 | 0.203199 | 0 |
| Up | ZFP36L2 | 3 | 9.25E-05 | 15144 | 0.307451 | 0 |
| Up | SLC2A5 | 3 | 2.76E-05 | 1624 | 0.241061 | 0.333333 |
| Up | CTSH | 3 | 3.51E-04 | 16788 | 0.306866 | 0 |
| Up | TNRC18 | 3 | 1.79E-04 | 47338 | 0.30887 | 0 |
| Up | CCDC80 | 3 | 1.95E-05 | 2102 | 0.216866 | 0 |
| Up | MGP | 3 | 7.32E-04 | 41846 | 0.207737 | 0 |
| Up | OASL | 3 | 8.88E-04 | 34710 | 0.306866 | 0 |
| Up | CFD | 3 | 7.62E-04 | 32594 | 0.307313 | 0 |
| Up | MRVI1 | 3 | 7.40E-04 | 53206 | 0.201468 | 0 |
| Up | LIMS2 | 3 | 7.78E-04 | 98670 | 0.228267 | 0 |
| Up | PTGDR | 3 | 0.001457 | 181566 | 0.22719 | 0 |
| Up | FOXS1 | 3 | 7.53E-04 | 101218 | 0.22719 | 0 |
| Up | IL18RAP | 3 | 7.60E-04 | 102434 | 0.232306 | 0 |
| Up | CCDC107 | 3 | 2.43E-04 | 15402 | 0.24236 | 0 |
| Up | TSPO | 3 | 9.03E-04 | 45462 | 0.307451 | 0 |
| Up | CCR7 | 3 | 7.29E-04 | 18830 | 0.246607 | 0.333333 |
| Up | DERL3 | 3 | 1 | 6 | 1 | 0 |
| Up | IGFLR1 | 3 | 1 | 6 | 1 | 0 |
| Up | MRC2 | 2 | 7.29E-04 | 22152 | 0.306798 | 0 |
| Up | RNPEPL1 | 2 | 2.21E-04 | 49220 | 0.308349 | 0 |
| Up | ZNF418 | 2 | 6.13E-04 | 75602 | 0.308211 | 0 |
| Up | IFI44 | 2 | 1.96E-04 | 30040 | 0.307348 | 0 |
| Up | LILRB4 | 2 | 1.14E-06 | 194 | 0.211126 | 0 |
| Up | ANKRD35 | 2 | 1.01E-05 | 1834 | 0.237967 | 0 |
| Up | ERICH1 | 2 | 9.17E-05 | 23458 | 0.307692 | 0 |
| Up | EGFLAM | 2 | 1.09E-04 | 10200 | 0.306798 | 0 |
| Up | PRSS8 | 2 | 7.29E-04 | 40016 | 0.217243 | 0 |
| Up | CHIT1 | 2 | 2.68E-05 | 1942 | 0.210526 | 0 |
| Up | NBL1 | 2 | 9.17E-05 | 6794 | 0.222402 | 0 |
| Up | HLA-DOB | 2 | 3.24E-06 | 238 | 0.202092 | 0 |
| Up | EMILIN2 | 2 | 3.05E-05 | 472 | 0.20403 | 0 |
| Up | EGFL6 | 2 | 1.44E-05 | 848 | 0.202749 | 0 |
| Up | TFEB | 2 | 4.31E-05 | 4334 | 0.230569 | 0 |
| Up | CYTH4 | 2 | 7.29E-04 | 22152 | 0.306798 | 0 |
| Up | SIRPG | 2 | 7.29E-04 | 22152 | 0.306798 | 0 |
| Up | STAC2 | 2 | 7.81E-06 | 490 | 0.205343 | 0 |
| Up | CCDC88B | 2 | 7.29E-04 | 22450 | 0.16043 | 0 |
| Up | MS4A7 | 2 | 7.29E-04 | 20644 | 0.192413 | 0 |
| Up | SOX17 | 2 | 6.16E-05 | 6626 | 0.233532 | 0 |
| Up | CDH7 | 2 | 7.29E-04 | 48342 | 0.207643 | 0 |
| Up | EMCN | 2 | 1.77E-05 | 356 | 0.241954 | 0 |
| Up | RNASE1 | 2 | 4.75E-05 | 2714 | 0.224018 | 0 |
| Up | SUSD2 | 2 | 7.29E-04 | 108014 | 0.211777 | 0 |
| Up | IL3RA | 2 | 7.29E-04 | 101316 | 0.215199 | 0 |
| Up | CXCL14 | 2 | 4.30E-05 | 1800 | 0.229643 | 0 |
| Up | SULF1 | 2 | 3.47E-05 | 5820 | 0.230356 | 0 |
| Up | GIMAP4 | 2 | 4.39E-04 | 130270 | 0.31228 | 0 |
| Up | CLDN4 | 2 | 9.08E-06 | 508 | 0.200175 | 0 |
| Up | FAT3 | 2 | 7.29E-04 | 35266 | 0.212236 | 0 |
| Up | S1PR4 | 2 | 7.29E-04 | 185770 | 0.158677 | 0 |
| Up | RAMP1 | 2 | 1 | 2 | 1 | 0 |
| Up | MARCO | 2 | 1 | 2 | 1 | 0 |
| Up | ICOS | 2 | 1 | 2 | 1 | 0 |
| Up | CLEC11A | 1 | 0 | 0 | 0.306729 | 0 |
| Up | CPNE5 | 1 | 0 | 0 | 0.306729 | 0 |
| Up | SMOC2 | 1 | 0 | 0 | 0.21217 | 0 |
| Up | GIMAP1 | 1 | 0 | 0 | 0.306729 | 0 |
| Up | ADCY4 | 1 | 0 | 0 | 0.306729 | 0 |
| Up | SLC15A3 | 1 | 0 | 0 | 0.164696 | 0 |
| Up | PLXDC1 | 1 | 0 | 0 | 0.306729 | 0 |
| Up | WDFY4 | 1 | 0 | 0 | 0.306729 | 0 |
| Up | MEI1 | 1 | 0 | 0 | 0.195762 | 0 |
| Up | CMPK2 | 1 | 0 | 0 | 0.306729 | 0 |
| Up | SIGLEC14 | 1 | 0 | 0 | 0.186261 | 0 |
| Up | DPT | 1 | 0 | 0 | 0.222366 | 0 |
| Up | FMOD | 1 | 0 | 0 | 0.181818 | 0 |
| Up | ARHGAP25 | 1 | 0 | 0 | 0.306729 | 0 |
| Up | CAPG | 1 | 0 | 0 | 0.306729 | 0 |
| Up | CD52 | 1 | 0 | 0 | 0.228781 | 0 |
| Up | HLA-DOA | 1 | 0 | 0 | 0.306729 | 0 |
| Up | CILP | 1 | 0 | 0 | 0.181818 | 0 |
| Up | SLIT3 | 1 | 0 | 0 | 0.196393 | 0 |
| Up | CD163 | 1 | 0 | 0 | 0.200864 | 0 |
| Up | AEBP1 | 1 | 0 | 0 | 0.222709 | 0 |
| Up | RTP4 | 1 | 0 | 0 | 0.208447 | 0 |
| Up | TMEM176B | 1 | 0 | 0 | 0.223708 | 0 |
| Up | ZNF524 | 1 | 0 | 0 | 0.2047 | 0 |
| Up | TBC1D10C | 1 | 0 | 0 | 0.227115 | 0 |
| Up | GIMAP6 | 1 | 0 | 0 | 0.227115 | 0 |
| Up | GNLY | 1 | 0 | 0 | 0.227115 | 0 |
| Down | MYC | 713 | 0.205435 | 87315356 | 0.42433 | 0.002565 |
| Down | HSP90AB1 | 563 | 0.150517 | 90042474 | 0.396225 | 0.003654 |
| Down | BRCA1 | 403 | 0.097657 | 34462280 | 0.40416 | 0.00537 |
| Down | EEF1A1 | 283 | 0.050657 | 28627038 | 0.37761 | 0.015488 |
| Down | HNRNPA1 | 237 | 0.04436 | 23745104 | 0.378339 | 0.00708 |
| Down | PCNA | 228 | 0.039118 | 25588738 | 0.362138 | 0.006994 |
| Down | NPM1 | 203 | 0.029258 | 23118776 | 0.364496 | 0.008974 |
| Down | GAPDH | 183 | 0.03126 | 13310882 | 0.386265 | 0.02462 |
| Down | TUBB | 173 | 0.0226 | 13312300 | 0.383805 | 0.019089 |
| Down | RPS2 | 156 | 0.010184 | 6684826 | 0.367143 | 0.079239 |
| Down | ALB | 153 | 0.043375 | 14319068 | 0.347267 | 0.003612 |
| Down | RPSA | 152 | 0.017762 | 6529218 | 0.386709 | 0.07642 |
| Down | NEDD4L | 147 | 0.035604 | 15156786 | 0.340774 | 2.80E-04 |
| Down | CDK1 | 146 | 0.020478 | 10576650 | 0.359453 | 0.014077 |
| Down | RPL6 | 144 | 0.01277 | 6222386 | 0.384559 | 0.092852 |
| Down | RBBP4 | 141 | 0.022261 | 13515558 | 0.35396 | 0.004965 |
| Down | RPL23A | 140 | 0.007195 | 5887530 | 0.353826 | 0.085098 |
| Down | SNCA | 140 | 0.029059 | 10160206 | 0.355216 | 0.005755 |
| Down | RPL4 | 136 | 0.006183 | 5067606 | 0.355512 | 0.093682 |
| Down | TUBA1A | 133 | 0.015507 | 8639510 | 0.365403 | 0.016632 |
| Down | RPL7 | 129 | 0.005207 | 4435368 | 0.360279 | 0.098595 |
| Down | C1QBP | 128 | 0.025126 | 11254578 | 0.364553 | 0.006029 |
| Down | KPNA2 | 127 | 0.024291 | 8493838 | 0.383868 | 0.012123 |
| Down | PRMT5 | 126 | 0.017703 | 13494440 | 0.341417 | 8.89E-04 |
| Down | RPLP0 | 124 | 0.006228 | 5001824 | 0.360334 | 0.095463 |
| Down | UBB | 123 | 0.0168 | 9698826 | 0.368495 | 0.007464 |
| Down | EEF1G | 122 | 0.021689 | 9177548 | 0.3689 | 0.011245 |
| Down | RPL7A | 116 | 0.005444 | 3815738 | 0.364807 | 0.117091 |
| Down | RPL3 | 112 | 0.008048 | 4666876 | 0.374572 | 0.094916 |
| Down | CCT2 | 111 | 0.011227 | 6217422 | 0.374214 | 0.044717 |
| Down | RPL15 | 110 | 0.002252 | 2856172 | 0.351289 | 0.13161 |
| Down | RPS10 | 109 | 0.007854 | 6158830 | 0.346168 | 0.033639 |
| Down | SKP2 | 105 | 0.01603 | 5513256 | 0.380482 | 0.022711 |
| Down | PSMA5 | 100 | 0.011632 | 6978802 | 0.359948 | 0.024646 |
| Down | SLC25A5 | 98 | 0.004847 | 4657438 | 0.346551 | 0.072586 |
| Down | CCNB1 | 98 | 0.011903 | 3857234 | 0.374006 | 0.042499 |
| Down | ENO1 | 96 | 0.0154 | 5859714 | 0.385853 | 0.036842 |
| Down | MYB | 92 | 0.018296 | 4267036 | 0.360003 | 0.010511 |
| Down | UBE2E3 | 90 | 0.019166 | 4652570 | 0.358986 | 0.007241 |
| Down | DDX21 | 90 | 0.007728 | 4881518 | 0.348891 | 0.026217 |
| Down | CDC20 | 89 | 0.012468 | 4329128 | 0.343688 | 0.028345 |
| Down | CCT4 | 89 | 0.00519 | 4277996 | 0.360666 | 0.052094 |
| Down | KRT18 | 89 | 0.013561 | 8188838 | 0.34179 | 0.002554 |
| Down | TRIM63 | 88 | 0.017034 | 4536234 | 0.340799 | 0.00627 |
| Down | PRDX1 | 88 | 0.012588 | 6795082 | 0.362669 | 0.015152 |
| Down | CCT8 | 85 | 0.004891 | 4403926 | 0.358931 | 0.054342 |
| Down | EIF3E | 85 | 0.01015 | 3943706 | 0.36365 | 0.028291 |
| Down | MCM2 | 84 | 0.011649 | 5324180 | 0.351999 | 0.012622 |
| Down | PSMD6 | 84 | 0.006594 | 3711384 | 0.337061 | 0.022662 |
| Down | RBMX | 83 | 0.009945 | 5139612 | 0.350086 | 0.021746 |
| Down | APOA1 | 83 | 0.019519 | 4233830 | 0.333807 | 0.016456 |
| Down | CCT6A | 80 | 0.005059 | 3370390 | 0.360472 | 0.062025 |
| Down | EIF3I | 79 | 0.00966 | 4532448 | 0.355351 | 0.011035 |
| Down | VTN | 78 | 0.013278 | 4330074 | 0.326732 | 0.001665 |
| Down | CSE1L | 78 | 0.00907 | 4428662 | 0.358712 | 0.018648 |
| Down | PHB2 | 77 | 0.011294 | 5386698 | 0.351814 | 0.007177 |
| Down | CCNA2 | 77 | 0.005002 | 2685430 | 0.355835 | 0.053999 |
| Down | PSMB5 | 77 | 0.009234 | 3457918 | 0.370499 | 0.042379 |
| Down | DNMT3B | 75 | 0.01232 | 3955286 | 0.35428 | 0.011892 |
| Down | MAVS | 74 | 0.01191 | 4009030 | 0.336819 | 0.001111 |
| Down | SLC25A3 | 71 | 0.012487 | 4850904 | 0.35372 | 0.005634 |
| Down | TUBA1B | 69 | 0.005616 | 3262686 | 0.345608 | 0.036658 |
| Down | IARS | 68 | 0.004923 | 3192328 | 0.342462 | 0.035119 |
| Down | LDHA | 66 | 0.005682 | 3384634 | 0.355916 | 0.041492 |
| Down | POU5F1 | 63 | 0.009683 | 2709306 | 0.351867 | 0.011265 |
| Down | SOX2 | 63 | 0.011281 | 2027862 | 0.334519 | 0.016385 |
| Down | NR0B2 | 60 | 0.007839 | 3384446 | 0.342886 | 0.001695 |
| Down | CAPNS1 | 59 | 0.008438 | 3550596 | 0.341145 | 0.009351 |
| Down | LDHB | 59 | 0.003764 | 3344120 | 0.33545 | 0.021625 |
| Down | IGF2BP3 | 57 | 0.007333 | 2298220 | 0.360003 | 0.039474 |
| Down | HIST1H1C | 55 | 0.004697 | 3059682 | 0.338419 | 0.016162 |
| Down | PAICS | 55 | 0.006134 | 3579222 | 0.333901 | 0.002694 |
| Down | BUB1B | 53 | 0.007217 | 1568390 | 0.340281 | 0.032656 |
| Down | PDIA6 | 52 | 0.006665 | 4090230 | 0.337812 | 0.010558 |
| Down | MBP | 51 | 0.005193 | 1684416 | 0.327415 | 0.001569 |
| Down | MET | 51 | 0.010548 | 3284818 | 0.333192 | 0 |
| Down | TTR | 49 | 0.008087 | 1488760 | 0.334448 | 0.010204 |
| Down | TNNT1 | 49 | 0.010948 | 1536610 | 0.350425 | 0.008503 |
| Down | EEF1A2 | 48 | 0.002097 | 1982718 | 0.332932 | 0 |
| Down | ABCC2 | 48 | 0.010493 | 1568726 | 0.341269 | 0.005319 |
| Down | PGK1 | 47 | 0.004818 | 1903066 | 0.356969 | 0.07123 |
| Down | TRIM37 | 46 | 0.007518 | 1753190 | 0.348632 | 0.007729 |
| Down | FHL1 | 45 | 0.006795 | 2104596 | 0.33533 | 0.00404 |
| Down | TFRC | 45 | 0.006068 | 3188390 | 0.333475 | 0 |
| Down | RHOXF2 | 43 | 0.011517 | 1750118 | 0.274457 | 0 |
| Down | APOB | 43 | 0.006945 | 2271724 | 0.33786 | 0.009967 |
| Down | PFKP | 42 | 0.003266 | 1640188 | 0.339544 | 0.00813 |
| Down | ENO2 | 42 | 0.0052 | 2675544 | 0.332956 | 0.003484 |
| Down | H2AFZ | 42 | 0.005388 | 1884076 | 0.329757 | 0.001161 |
| Down | TKT | 41 | 0.003583 | 1462950 | 0.353401 | 0.028049 |
| Down | MAGEA6 | 40 | 0.005724 | 1721840 | 0.329687 | 0 |
| Down | GATA3 | 40 | 0.008796 | 1432218 | 0.349669 | 0.014103 |
| Down | ACAT1 | 40 | 0.004855 | 2293968 | 0.332086 | 0.00641 |
| Down | TYMS | 40 | 0.007021 | 1967804 | 0.337279 | 0.007692 |
| Down | PGD | 38 | 0.00319 | 1432784 | 0.331454 | 0.036984 |
| Down | TPI1 | 37 | 0.003586 | 1296678 | 0.345862 | 0.09009 |
| Down | IMPDH2 | 36 | 0.002276 | 1522866 | 0.335426 | 0.01746 |
| Down | FGFR3 | 36 | 0.007363 | 1436054 | 0.317103 | 0.007937 |
| Down | PTTG1 | 36 | 0.002181 | 635618 | 0.337545 | 0.053968 |
| Down | ASF1B | 36 | 0.004296 | 935530 | 0.336001 | 0.02381 |
| Down | RFC5 | 35 | 0.003115 | 1013766 | 0.36147 | 0.055462 |
| Down | PLA2G4A | 35 | 0.003901 | 1353946 | 0.331664 | 0 |
| Down | GNAS | 33 | 0.006431 | 1316640 | 0.330964 | 0.007576 |
| Down | TRIB3 | 33 | 0.00524 | 1451550 | 0.340084 | 0.011364 |
| Down | CDC45 | 33 | 0.002712 | 1142452 | 0.338468 | 0.034091 |
| Down | BMX | 33 | 0.004466 | 1042152 | 0.34915 | 0.017045 |
| Down | SERPINA1 | 32 | 0.005464 | 1480486 | 0.334305 | 0 |
| Down | TPX2 | 32 | 0.003838 | 1028888 | 0.344545 | 0.016129 |
| Down | RANBP1 | 31 | 0.002458 | 1306090 | 0.332979 | 0 |
| Down | KIF23 | 28 | 0.002728 | 1417572 | 0.327825 | 0 |
| Down | ID1 | 27 | 0.003239 | 771330 | 0.332438 | 0.008547 |
| Down | OLA1 | 27 | 0.003068 | 727676 | 0.338956 | 0.031339 |
| Down | SEC23B | 26 | 0.004471 | 1519614 | 0.329872 | 0 |
| Down | IGF2BP2 | 26 | 0.001427 | 660150 | 0.355324 | 0.058462 |
| Down | ENO3 | 25 | 0.002857 | 559136 | 0.343262 | 0.066667 |
| Down | IGFBP3 | 25 | 0.002823 | 1764080 | 0.31355 | 0.003333 |
| Down | F2 | 24 | 0.004813 | 1568314 | 0.291651 | 0.043478 |
| Down | NFE2 | 24 | 0.003738 | 2099820 | 0.285091 | 0 |
| Down | GMNN | 24 | 0.002113 | 741708 | 0.332508 | 0.028986 |
| Down | PSAT1 | 24 | 0.003089 | 738494 | 0.3484 | 0.021739 |
| Down | EGLN1 | 23 | 0.003385 | 798698 | 0.328146 | 0 |
| Down | PGM1 | 23 | 0.00253 | 753590 | 0.333357 | 0.007905 |
| Down | CD55 | 22 | 0.004056 | 696952 | 0.33129 | 0 |
| Down | FOXA2 | 21 | 0.004992 | 872700 | 0.281405 | 0.019048 |
| Down | PBK | 21 | 0.00267 | 546972 | 0.354386 | 0.047619 |
| Down | AJUBA | 21 | 0.001964 | 617438 | 0.335833 | 0.02381 |
| Down | HP | 20 | 0.001845 | 455036 | 0.314221 | 0.031579 |
| Down | TFF1 | 20 | 0.002374 | 1520442 | 0.305285 | 0 |
| Down | BEGAIN | 20 | 0.004469 | 651432 | 0.330452 | 0 |
| Down | GDA | 20 | 0.00349 | 588498 | 0.329826 | 0.036842 |
| Down | MAD2L1BP | 19 | 0.002165 | 392308 | 0.339912 | 0.052632 |
| Down | KIF11 | 19 | 0.001601 | 423324 | 0.333475 | 0.011696 |
| Down | HBG1 | 19 | 0.00152 | 803574 | 0.311845 | 0 |
| Down | CPS1 | 19 | 0.001046 | 359790 | 0.348013 | 0.05848 |
| Down | ITGB7 | 19 | 0.004777 | 992014 | 0.329226 | 0 |
| Down | AMBP | 18 | 0.003475 | 642842 | 0.320956 | 0.045752 |
| Down | MRPL10 | 18 | 0.003042 | 444244 | 0.334162 | 0.03268 |
| Down | CGA | 18 | 0.003848 | 592198 | 0.306479 | 0.006536 |
| Down | AFP | 18 | 0.001988 | 422106 | 0.2893 | 0 |
| Down | IGFBP1 | 18 | 0.001208 | 551500 | 0.288927 | 0.006536 |
| Down | CENPO | 18 | 0.002863 | 489612 | 0.350922 | 0.091503 |
| Down | CENPH | 17 | 0.001891 | 359996 | 0.330777 | 0.095588 |
| Down | CDCA5 | 17 | 0.001718 | 456258 | 0.330801 | 0 |
| Down | PGAM1 | 17 | 0.00118 | 280570 | 0.342487 | 0.080882 |
| Down | IGF2 | 17 | 0.003588 | 782808 | 0.275486 | 0.014706 |
| Down | AKR1B1 | 17 | 0.001528 | 523568 | 0.331664 | 0 |
| Down | FDFT1 | 17 | 0.002414 | 651582 | 0.330243 | 0 |
| Down | HBG2 | 17 | 0.001113 | 334894 | 0.320671 | 0.066176 |
| Down | HMMR | 17 | 0.00107 | 246928 | 0.340379 | 0.058824 |
| Down | AHSG | 17 | 0.002005 | 497564 | 0.313258 | 0.022059 |
| Down | APOA2 | 16 | 5.79E-04 | 137538 | 0.29176 | 0.1 |
| Down | APOH | 15 | 0.00198 | 1212586 | 0.297576 | 0 |
| Down | KIF2C | 15 | 0.001078 | 393602 | 0.331617 | 0.009524 |
| Down | RNF43 | 15 | 0.001084 | 220414 | 0.334495 | 0.047619 |
| Down | ALDOC | 15 | 0.001343 | 400682 | 0.329387 | 0.009524 |
| Down | APOC3 | 14 | 0.001271 | 431288 | 0.33803 | 0.087912 |
| Down | CPNE1 | 14 | 0.001438 | 530794 | 0.329549 | 0 |
| Down | HMGA2 | 14 | 0.001654 | 271978 | 0.334757 | 0.032967 |
| Down | ASGR1 | 14 | 9.33E-04 | 506086 | 0.294852 | 0 |
| Down | DPPA4 | 14 | 9.72E-04 | 483504 | 0.281321 | 0 |
| Down | AGR2 | 14 | 0.002379 | 249434 | 0.30724 | 0 |
| Down | PEG10 | 14 | 0.001408 | 303552 | 0.327323 | 0 |
| Down | ADA | 14 | 0.002493 | 415608 | 0.332414 | 0.032967 |
| Down | CENPN | 13 | 0.001246 | 216110 | 0.32918 | 0.179487 |
| Down | BNIP3 | 13 | 0.002082 | 358706 | 0.327917 | 0 |
| Down | PIR | 13 | 0.0018 | 907788 | 0.277077 | 0 |
| Down | PRIM1 | 13 | 3.21E-04 | 124150 | 0.327437 | 0 |
| Down | HBZ | 13 | 0.001144 | 489382 | 0.315211 | 0 |
| Down | COX4I1 | 13 | 0.002122 | 373360 | 0.330847 | 0 |
| Down | TFPI | 13 | 9.35E-04 | 374928 | 0.288697 | 0 |
| Down | FGG | 13 | 0.001136 | 470062 | 0.282114 | 0.038462 |
| Down | FST | 13 | 0.002668 | 735238 | 0.272738 | 0 |
| Down | LAMA1 | 13 | 0.002341 | 300044 | 0.328375 | 0 |
| Down | COASY | 13 | 0.00221 | 434498 | 0.333664 | 0.038462 |
| Down | QPCT | 12 | 1.58E-05 | 41260 | 0.327711 | 0 |
| Down | CTNNAL1 | 12 | 0.002284 | 342522 | 0.328077 | 0 |
| Down | STAR | 12 | 0.001173 | 546324 | 0.287338 | 0 |
| Down | NCAM1 | 12 | 6.47E-04 | 190710 | 0.334257 | 0.030303 |
| Down | CCNB2 | 12 | 2.10E-04 | 97062 | 0.334424 | 0.151515 |
| Down | HLF | 12 | 0.001817 | 256830 | 0.288572 | 0 |
| Down | PTP4A1 | 12 | 0.002403 | 431426 | 0.3315 | 0 |
| Down | GSTM3 | 12 | 3.03E-04 | 186268 | 0.328123 | 0 |
| Down | ASGR2 | 12 | 4.32E-04 | 277066 | 0.270432 | 0 |
| Down | CTNND2 | 12 | 0.001233 | 215538 | 0.331547 | 0 |
| Down | CARHSP1 | 12 | 7.10E-04 | 191384 | 0.327757 | 0 |
| Down | INA | 11 | 1.07E-04 | 136268 | 0.328558 | 0 |
| Down | ESPL1 | 11 | 0.001031 | 164318 | 0.3315 | 0.090909 |
| Down | KIF4A | 11 | 9.35E-04 | 224848 | 0.332273 | 0.036364 |
| Down | SPC25 | 11 | 0.001423 | 244986 | 0.327734 | 0 |
| Down | REPIN1 | 11 | 2.04E-04 | 68744 | 0.302516 | 0.054545 |
| Down | TCF7 | 11 | 0.001496 | 290812 | 0.327643 | 0 |
| Down | AGMAT | 10 | 0.001337 | 920392 | 0.276052 | 0 |
| Down | BAMBI | 10 | 0.001928 | 278690 | 0.326914 | 0 |
| Down | AGT | 10 | 6.54E-04 | 346990 | 0.294778 | 0 |
| Down | ANG | 10 | 0.001664 | 579520 | 0.291995 | 0 |
| Down | TBC1D7 | 10 | 8.03E-04 | 167126 | 0.334662 | 0.022222 |
| Down | ACY1 | 10 | 3.04E-04 | 182552 | 0.338225 | 0.044444 |
| Down | OIP5 | 10 | 0.001463 | 256898 | 0.326619 | 0 |
| Down | TAC3 | 10 | 0.001551 | 205706 | 0.327232 | 0 |
| Down | PTX3 | 10 | 0.002565 | 1004098 | 0.227512 | 0 |
| Down | C4BPA | 9 | 0.001339 | 472358 | 0.266327 | 0 |
| Down | ORM1 | 9 | 0.00101 | 549272 | 0.270634 | 0 |
| Down | RMI2 | 9 | 0.002581 | 293792 | 0.327096 | 0 |
| Down | SERPINC1 | 9 | 9.68E-04 | 241966 | 0.289158 | 0.111111 |
| Down | AKR1C3 | 9 | 4.82E-04 | 211076 | 0.330684 | 0 |
| Down | PPP1R3C | 9 | 5.97E-04 | 244542 | 0.332744 | 0 |
| Down | SAAL1 | 9 | 0.001925 | 225340 | 0.327118 | 0 |
| Down | MAGEC1 | 9 | 0.001886 | 221860 | 0.327688 | 0.027778 |
| Down | MAL | 9 | 1.21E-04 | 32686 | 0.277045 | 0 |
| Down | CELSR2 | 8 | 6.27E-04 | 152870 | 0.327141 | 0 |
| Down | DLX2 | 8 | 0.00179 | 236210 | 0.326959 | 0 |
| Down | FZD5 | 8 | 9.48E-04 | 155326 | 0.326505 | 0 |
| Down | CCDC136 | 8 | 4.60E-04 | 192812 | 0.263167 | 0 |
| Down | IL23A | 8 | 0.001304 | 477832 | 0.260167 | 0 |
| Down | HGD | 8 | 7.64E-04 | 393294 | 0.278522 | 0 |
| Down | ALAS2 | 8 | 5.41E-04 | 228948 | 0.278258 | 0 |
| Down | TYRP1 | 8 | 0.001307 | 605734 | 0.270339 | 0 |
| Down | NPTX1 | 8 | 0.001488 | 278684 | 0.327073 | 0 |
| Down | DLGAP5 | 8 | 5.16E-04 | 103456 | 0.335139 | 0.142857 |
| Down | GNG4 | 7 | 0.00132 | 384244 | 0.258849 | 0 |
| Down | RBP4 | 7 | 1.27E-04 | 36580 | 0.295389 | 0 |
| Down | TRIM7 | 7 | 4.98E-04 | 91946 | 0.308814 | 0.047619 |
| Down | EPHB6 | 7 | 5.70E-04 | 124884 | 0.303668 | 0.047619 |
| Down | CCNB1IP1 | 7 | 4.58E-04 | 83072 | 0.335258 | 0.238095 |
| Down | AREG | 7 | 0.001309 | 192106 | 0.330104 | 0 |
| Down | FOXRED2 | 7 | 5.95E-04 | 132040 | 0.326528 | 0 |
| Down | FOLR1 | 7 | 5.33E-04 | 101898 | 0.328146 | 0.095238 |
| Down | SPAG5 | 7 | 9.69E-04 | 105098 | 0.332179 | 0.047619 |
| Down | VIL1 | 6 | 4.50E-04 | 233426 | 0.274297 | 0 |
| Down | MIPEP | 6 | 1.47E-04 | 64910 | 0.326664 | 0 |
| Down | MST1 | 6 | 8.72E-04 | 564148 | 0.261179 | 0 |
| Down | SOX3 | 6 | 2.61E-05 | 23258 | 0.292486 | 0 |
| Down | TESC | 6 | 0.001863 | 181202 | 0.32655 | 0 |
| Down | NANOG | 6 | 0.003614 | 321302 | 0.328513 | 0 |
| Down | TMEM147 | 6 | 0.001022 | 145542 | 0.326369 | 0 |
| Down | ORM2 | 5 | 4.70E-04 | 298836 | 0.258735 | 0 |
| Down | GABRB1 | 5 | 8.64E-04 | 130460 | 0.326482 | 0 |
| Down | GFRA1 | 5 | 8.53E-04 | 697882 | 0.244616 | 0 |
| Down | EPS8L3 | 5 | 8.87E-04 | 307116 | 0.254162 | 0 |
| Down | TYR | 5 | 2.18E-04 | 103128 | 0.3268 | 0 |
| Down | RNF182 | 5 | 4.70E-04 | 84816 | 0.326936 | 0 |
| Down | MLANA | 5 | 5.45E-04 | 118846 | 0.326914 | 0 |
| Down | SLC7A5 | 5 | 8.06E-05 | 59374 | 0.327825 | 0 |
| Down | HIC2 | 5 | 4.48E-04 | 177072 | 0.280599 | 0 |
| Down | LHX2 | 5 | 8.66E-04 | 103008 | 0.236365 | 0 |
| Down | PTPRH | 4 | 4.46E-04 | 238256 | 0.262535 | 0 |
| Down | RAD54L | 4 | 8.58E-04 | 93666 | 0.326165 | 0 |
| Down | PLP1 | 4 | 4.52E-04 | 105346 | 0.248902 | 0 |
| Down | GPC3 | 4 | 2.27E-04 | 90714 | 0.32826 | 0 |
| Down | APOM | 4 | 4.43E-04 | 365960 | 0.252714 | 0 |
| Down | TNFSF9 | 4 | 4.35E-04 | 55354 | 0.326165 | 0 |
| Down | CX3CR1 | 4 | 8.51E-04 | 696212 | 0.22759 | 0 |
| Down | FABP1 | 4 | 2.03E-04 | 38604 | 0.256282 | 0 |
| Down | MAT1A | 4 | 5.34E-04 | 74956 | 0.326165 | 0 |
| Down | ABCG5 | 4 | 1.07E-04 | 78982 | 0.259765 | 0 |
| Down | CDH18 | 4 | 4.91E-04 | 88100 | 0.326392 | 0 |
| Down | TFPI2 | 4 | 1.62E-05 | 6890 | 0.254575 | 0 |
| Down | TMEM97 | 4 | 4.65E-05 | 62730 | 0.330173 | 0 |
| Down | DPCD | 4 | 2.98E-05 | 30284 | 0.330173 | 0 |
| Down | F11 | 4 | 6.78E-05 | 20816 | 0.25463 | 0 |
| Down | KCNQ2 | 4 | 4.36E-04 | 124824 | 0.257714 | 0 |
| Down | CARD10 | 4 | 8.52E-04 | 100582 | 0.326324 | 0 |
| Down | NAA40 | 3 | 4.79E-04 | 56384 | 0.32612 | 0 |
| Down | AKR1C1 | 3 | 4.27E-04 | 194386 | 0.296937 | 0 |
| Down | XK | 3 | 4.26E-04 | 291070 | 0.274169 | 0 |
| Down | NDUFC2 | 3 | 4.36E-04 | 66938 | 0.3268 | 0 |
| Down | TMEM45B | 3 | 8.51E-04 | 629030 | 0.208459 | 0 |
| Down | KDM5D | 3 | 3.94E-05 | 21036 | 0.32612 | 0 |
| Down | NMU | 3 | 8.51E-04 | 108454 | 0.199635 | 0 |
| Down | AKR1B10 | 3 | 1.77E-05 | 13768 | 0.264098 | 0 |
| Down | BEX1 | 3 | 4.29E-04 | 48934 | 0.32612 | 0 |
| Down | MOCOS | 3 | 2.35E-04 | 96724 | 0.333688 | 0.333333 |
| Down | ITIH2 | 3 | 4.28E-04 | 256524 | 0.240807 | 0 |
| Down | FADS1 | 3 | 5.00E-04 | 63558 | 0.32612 | 0 |
| Down | KRT80 | 3 | 9.19E-06 | 14894 | 0.328375 | 0 |
| Down | RAB34 | 3 | 4.33E-04 | 51810 | 0.326505 | 0 |
| Down | FAM57A | 3 | 6.22E-05 | 25536 | 0.32612 | 0 |
| Down | CSAG1 | 3 | 2.16E-05 | 10552 | 0.274746 | 0 |
| Down | SERPIND1 | 3 | 4.86E-05 | 24834 | 0.273547 | 0.333333 |
| Down | PMEL | 3 | 2.91E-05 | 11556 | 0.250413 | 0 |
| Down | DCT | 3 | 9.41E-06 | 4328 | 0.242109 | 0 |
| Down | FGL1 | 3 | 4.30E-04 | 429502 | 0.234222 | 0 |
| Down | ADSSL1 | 3 | 5.63E-05 | 10700 | 0.274505 | 0 |
| Down | GDF6 | 3 | 3.75E-05 | 7180 | 0.247578 | 0 |
| Down | C8A | 3 | 1 | 6 | 1 | 0 |
| Down | OGDHL | 2 | 8.69E-06 | 10524 | 0.326959 | 0 |
| Down | AJAP1 | 2 | 3.70E-07 | 498 | 0.267585 | 0 |
| Down | PHACTR1 | 2 | 1.50E-06 | 1436 | 0.255335 | 0 |
| Down | CAMK2N1 | 2 | 1.67E-06 | 1268 | 0.26021 | 0 |
| Down | SH2D5 | 2 | 7.10E-07 | 384 | 0.264961 | 0 |
| Down | LIN28B | 2 | 3.66E-06 | 3450 | 0.261994 | 0 |
| Down | PPP1R26 | 2 | 4.25E-04 | 274936 | 0.245164 | 0 |
| Down | HYAL1 | 2 | 4.25E-04 | 41178 | 0.326075 | 0 |
| Down | ITIH3 | 2 | 4.70E-06 | 6466 | 0.267691 | 0 |
| Down | HCCS | 2 | 1.05E-05 | 19668 | 0.326755 | 0 |
| Down | SULT1A2 | 2 | 0 | 0 | 0.34627 | 1 |
| Down | CD8B | 2 | 4.08E-06 | 2652 | 0.257616 | 0 |
| Down | HPR | 2 | 1.33E-06 | 218 | 0.20844 | 0 |
| Down | BCAS1 | 2 | 7.50E-07 | 916 | 0.251041 | 0 |
| Down | LONRF2 | 2 | 4.25E-04 | 41178 | 0.326075 | 0 |
| Down | FAM178B | 2 | 3.49E-06 | 1036 | 0.224546 | 0 |
| Down | CYP2F1 | 2 | 1.09E-06 | 670 | 0.26559 | 0 |
| Down | GLP2R | 2 | 9.00E-06 | 6666 | 0.326324 | 0 |
| Down | RPS4Y1 | 2 | 9.00E-06 | 6666 | 0.326324 | 0 |
| Down | DLK1 | 2 | 4.25E-04 | 234974 | 0.205894 | 0 |
| Down | KISS1R | 2 | 4.25E-04 | 315046 | 0.231022 | 0 |
| Down | FADS2 | 2 | 4.25E-04 | 41178 | 0.326075 | 0 |
| Down | KLK1 | 2 | 7.31E-06 | 1102 | 0.21277 | 0 |
| Down | DCDC2 | 2 | 6.64E-06 | 1740 | 0.272185 | 0 |
| Down | PTTG2 | 2 | 2.46E-05 | 7724 | 0.269332 | 0 |
| Down | GREB1 | 2 | 3.97E-06 | 3266 | 0.283219 | 0 |
| Down | GDF11 | 2 | 1 | 2 | 1 | 0 |
| Down | SPCS1 | 1 | 0 | 0 | 0.32603 | 0 |
| Down | FZD10 | 1 | 0 | 0 | 0.246552 | 0 |
| Down | ELOVL2 | 1 | 0 | 0 | 0.279349 | 0 |
| Down | ST6GALNAC1 | 1 | 0 | 0 | 0.32603 | 0 |
| Down | SPANXB1 | 1 | 0 | 0 | 0.283629 | 0 |
| Down | WDR54 | 1 | 0 | 0 | 0.32603 | 0 |
| Down | SEC22C | 1 | 0 | 0 | 0.32603 | 0 |
| Down | CAMK2N2 | 1 | 0 | 0 | 0.231045 | 0 |
| Down | GTSF1 | 1 | 0 | 0 | 0.32603 | 0 |
| Down | RPS4Y2 | 1 | 0 | 0 | 0.32603 | 0 |
| Down | RIMKLA | 1 | 0 | 0 | 0.32603 | 0 |
| Down | ABCA12 | 1 | 0 | 0 | 0.32603 | 0 |
| Down | SP5 | 1 | 0 | 0 | 0.218099 | 0 |
| Down | FAM72A | 1 | 0 | 0 | 0.32603 | 0 |
| Down | L1TD1 | 1 | 0 | 0 | 0.197854 | 0 |
| Down | SAPCD1 | 1 | 0 | 0 | 0.231045 | 0 |
| Down | MNX1 | 1 | 0 | 0 | 0.261964 | 0 |
| Down | GAL | 1 | 0 | 0 | 0.257432 | 0 |
| Down | BMP5 | 1 | 0 | 0 | 0.248783 | 0 |
| Down | UTS2 | 1 | 0 | 0 | 0.282335 | 0 |
| Down | ZIC3 | 1 | 0 | 0 | 0.20046 | 0 |
| Down | ITM2A | 1 | 0 | 0 | 0.32603 | 0 |
| Down | ONECUT2 | 1 | 0 | 0 | 0.234362 | 0 |
| Down | ACSM3 | 1 | 0 | 0 | 0.235241 | 0 |
| Down | UGT2B10 | 1 | 0 | 0 | 0.229724 | 0 |
| Down | TRIML2 | 1 | 0 | 0 | 0.283629 | 0 |
| Down | ERI2 | 1 | 0 | 0 | 0.248783 | 0 |
| Down | SLC7A2 | 1 | 0 | 0 | 0.32603 | 0 |

Module 4 (13 nodes and 43 edges), module 5 (12 nodes and 29 edges), module 6 (12 nodes and 21 edges) and module 15 (8 nodes and 16 edges) were key modules in this PPI network for up regulated genes (Fig.9). The results of patway and GO enricment analysis presented that up regulated hub genes were mainly enriched in herpes simplex infection, toxoplasmosis, chemokine signaling pathway, PI3K-Akt signaling pathway, immune response, regulation of immune system process, positive regulation of cell motility and hematopoietic or lymphoid organ development. Module 11 (64 nodes and 274 edges), module 18 (42 nodes and 116 edges), module 19 (42 nodes and 116 edges) and module 26 (32 nodes and 95 edges) were key modules in this PPI network for down regulated genes (Fig.10). The results of patway and GO enricment analysis presented that down regulated hub genes were mainly enriched in cell cycle, metabolism of amino acids and derivatives, ubiquitin mediated proteolysis, HIF-1-alpha transcription factor network, negative regulation of protein metabolic process, carboxylic acid metabolic process, mitotic cell cycle process and organonitrogen compound biosynthetic process.

**Fig. 9.**
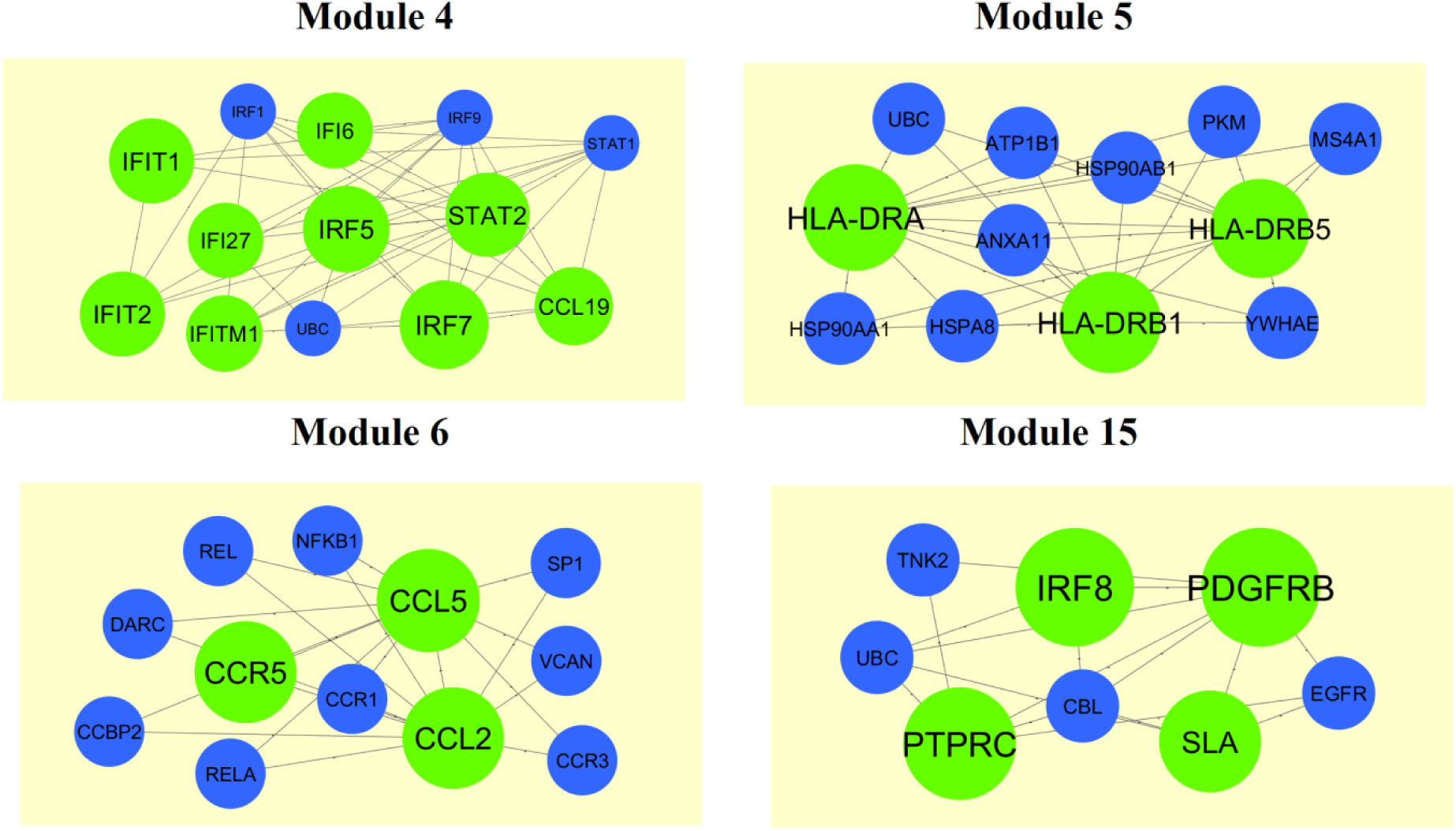
Modules in PPI network. The green nodes denote the up regulated genes

**Fig. 10.**
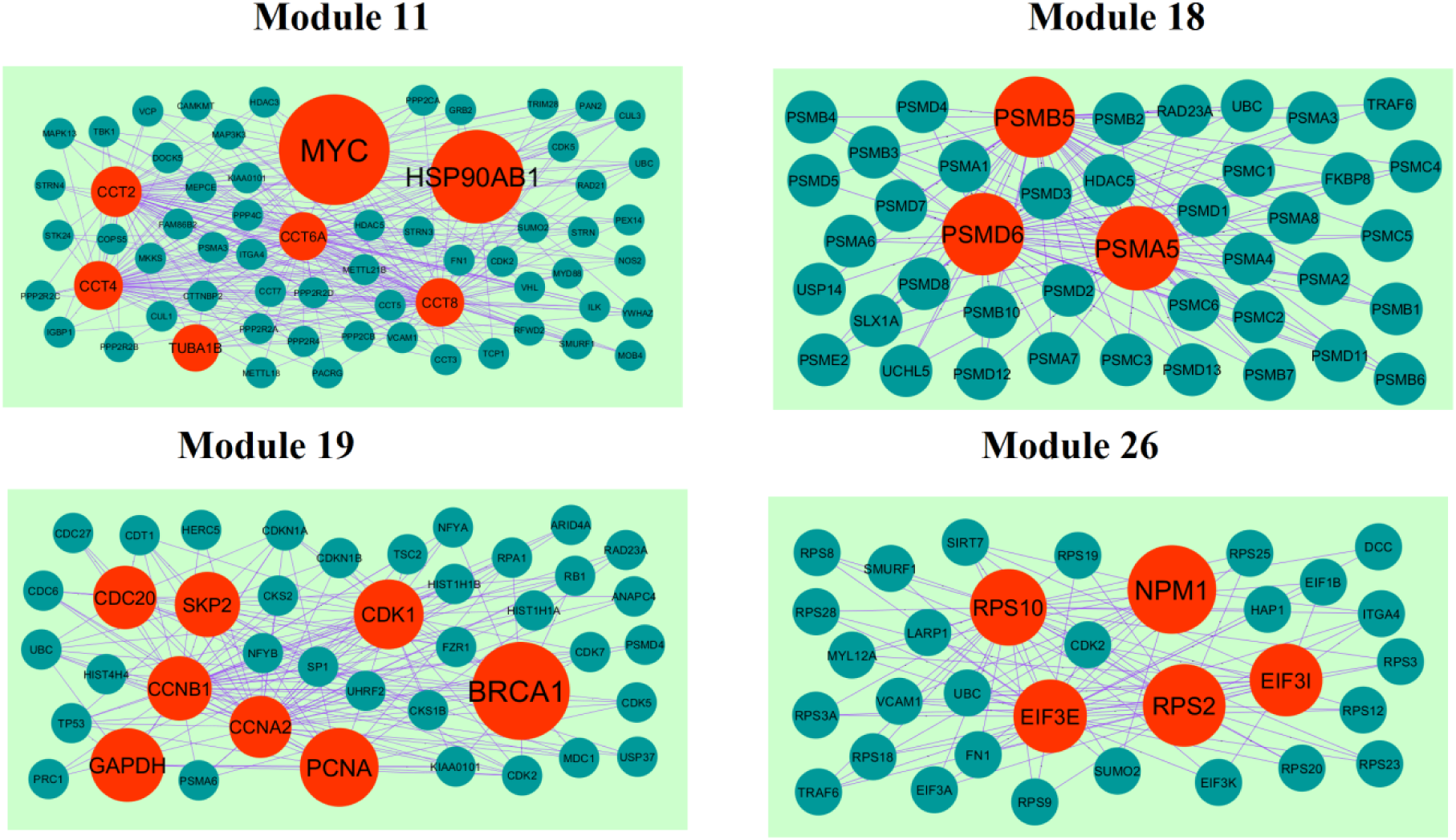
Modules in PPI network. The red nodes denote the down regulated genes.

### Construction of target gene - miRNA regulatory network

Expression of miRNAs was linked with pathogenesis of TNBC [Radojicic et al. 2011]. An up regulated target gene - miRNA regulatory network is shown in Fig. 11. Top five up regulated target genes such as CCDC80 interacts with 135 miRNAs (ex, hsa-mir-6499-3p), MYO1F interacts with 104 miRNAs (ex, hsa-mir-6882-3p), CDH7 interacts with 88 miRNAs (ex, hsa-mir-7845-5p), KLF2 interacts with 85 miRNAs (ex, hsa-mir-548as-3p) and FPR1 interacts with 85 miRNAs (ex, hsa-mir-4722-3p) are listed in Table 8. Pathway and GO enrichment analysis displayed that the top up regulated targate genes were primarily enriched in the extracellular structure organization, immune response, hematopoietic or lymphoid organ development and innate immune system. Similarly, down regulated target gene - miRNA regulatory network is shown in Fig. 12. Top five down regulated target genes such as PEG10 interacts with 139 miRNAs (ex, hsa-mir-4443), HNRNPA1 interacts with 139 miRNAs (ex, hsa-mir-6504-5p), PTP4A1 interacts with 132 miRNAs (ex, hsa-mir-548aq-5p), SLC7A5 interacts with 130 miRNAs (ex, hsa-mir-4434) and MYC interacts with 103 miRNAs (ex, hsa-mir-6507-3p) are listed in Table 8. Pathway and GO enrichment analysis displayed that the top down regulated targate genes were primarily enriched in the validated targets of C-MYC transcriptional activation, signaling events mediated by PRL, metabolism of amino acids and derivatives and carboxylic acid metabolic process.

**Fig. 11.**
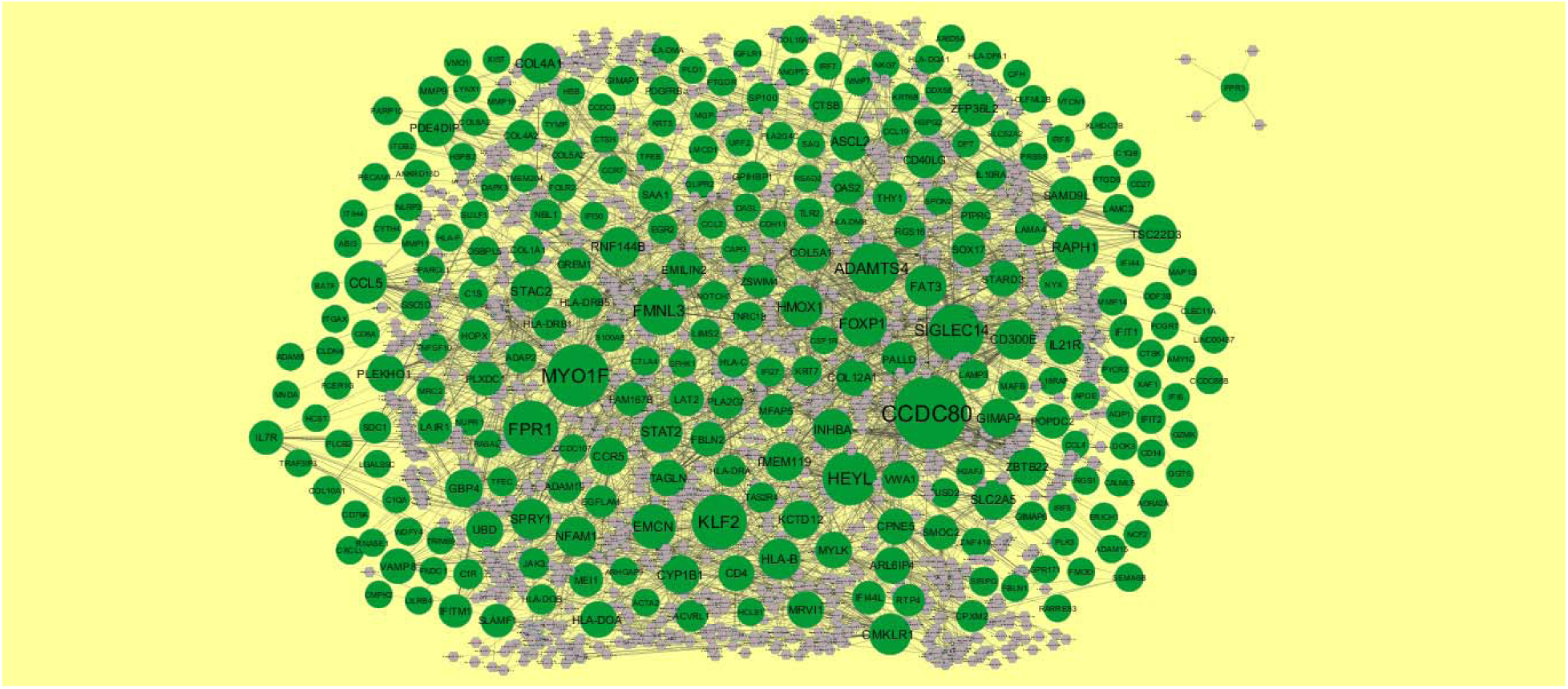
The network of up regulated genes and their related miRNAs. The green circles nodes are the up regulated DEGs, and gray diamond nodes are the miRNAs

**Fig. 12.**
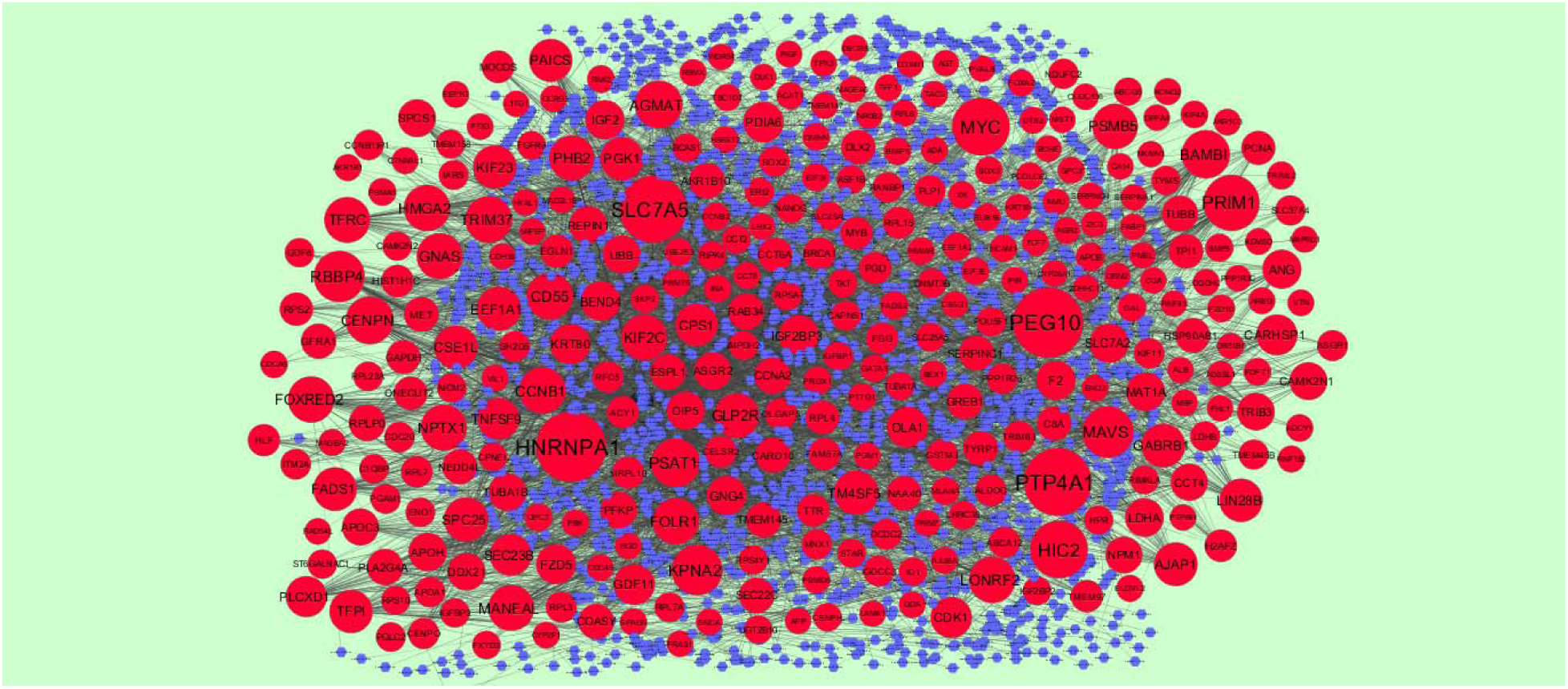
The network of down regulated genes and their related miRNAs. The red circles nodes are the down regulated DEGs, and blue diamond nodes are the miRNAs

**Table 8.**
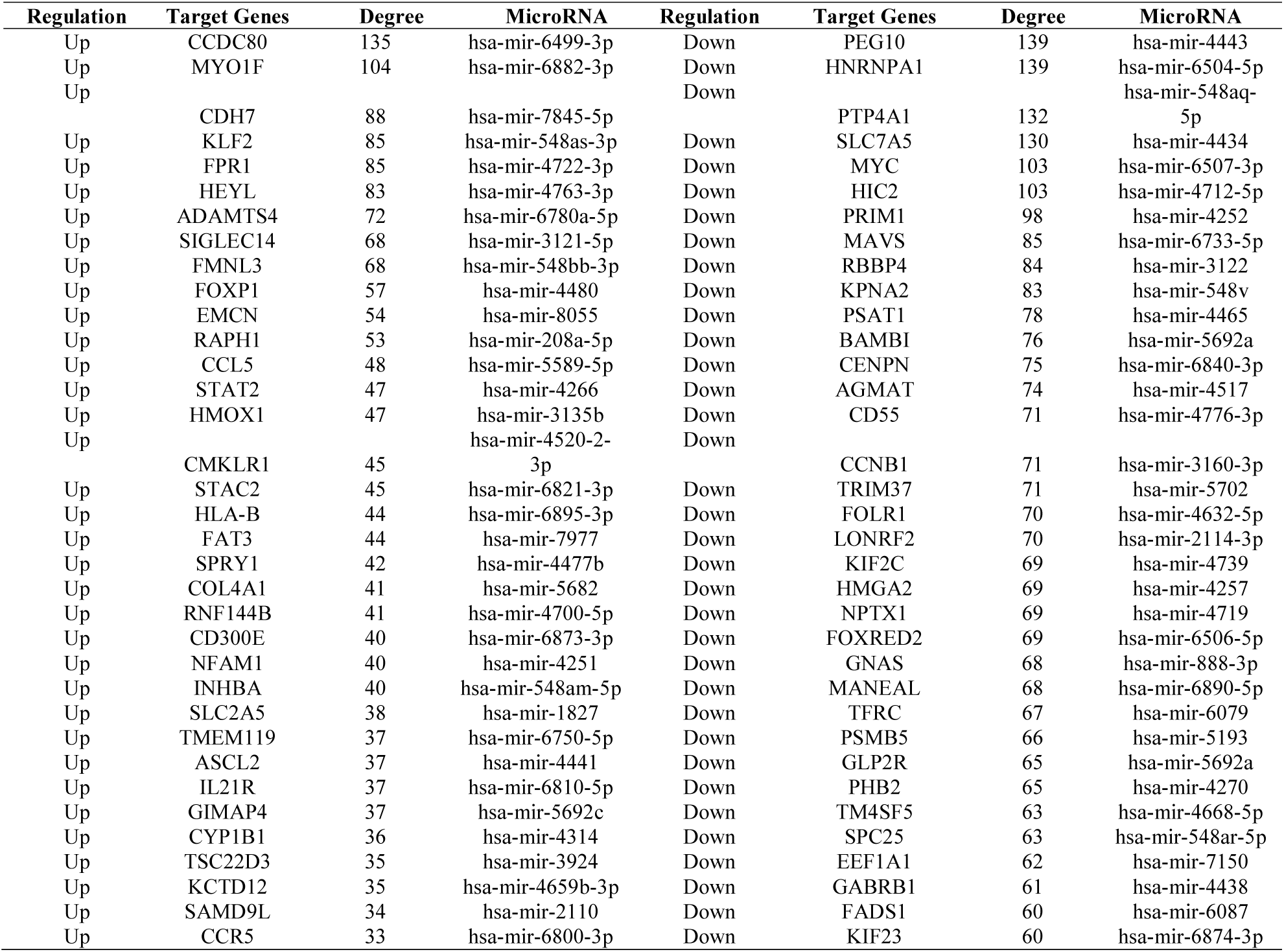
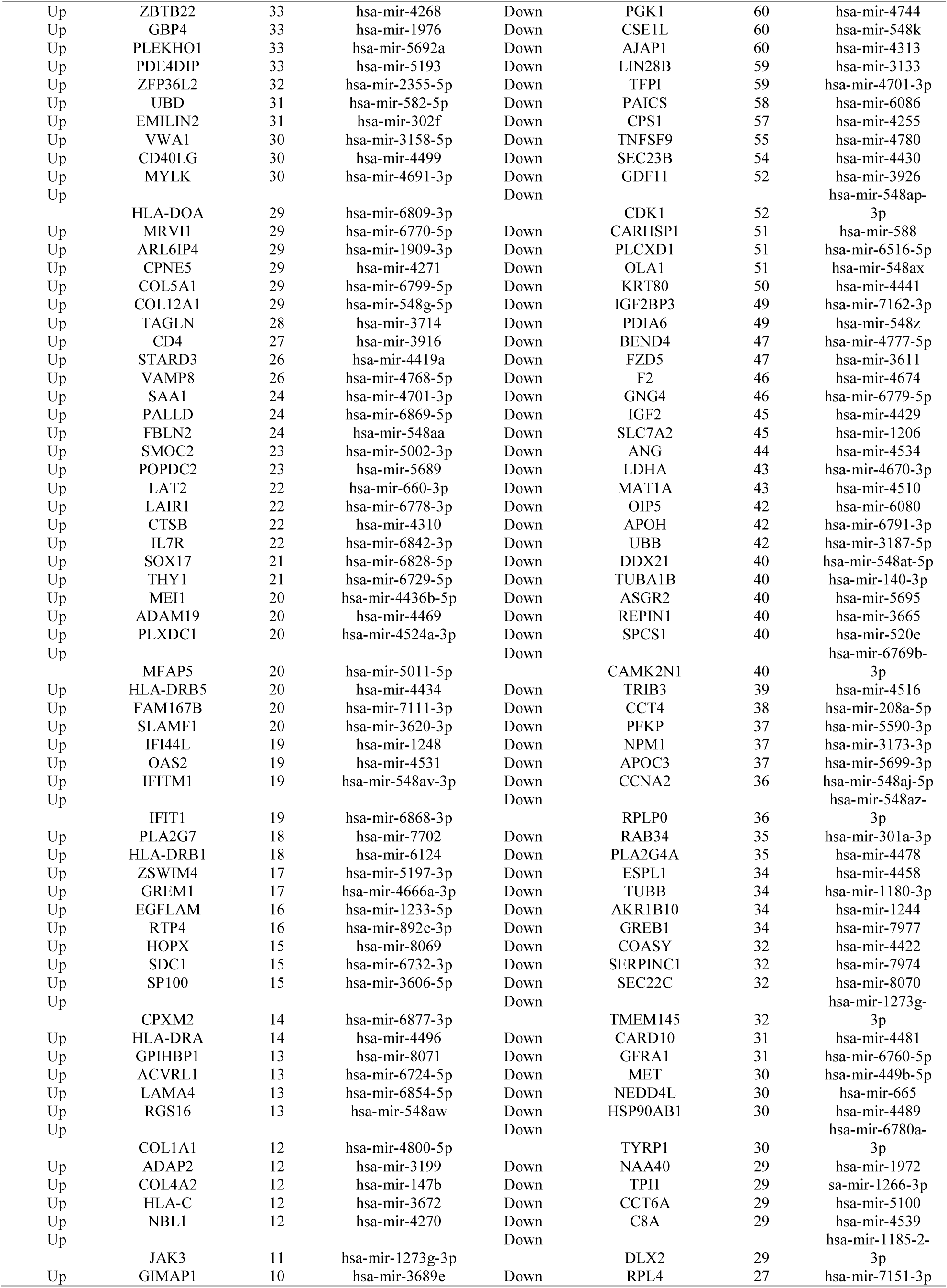
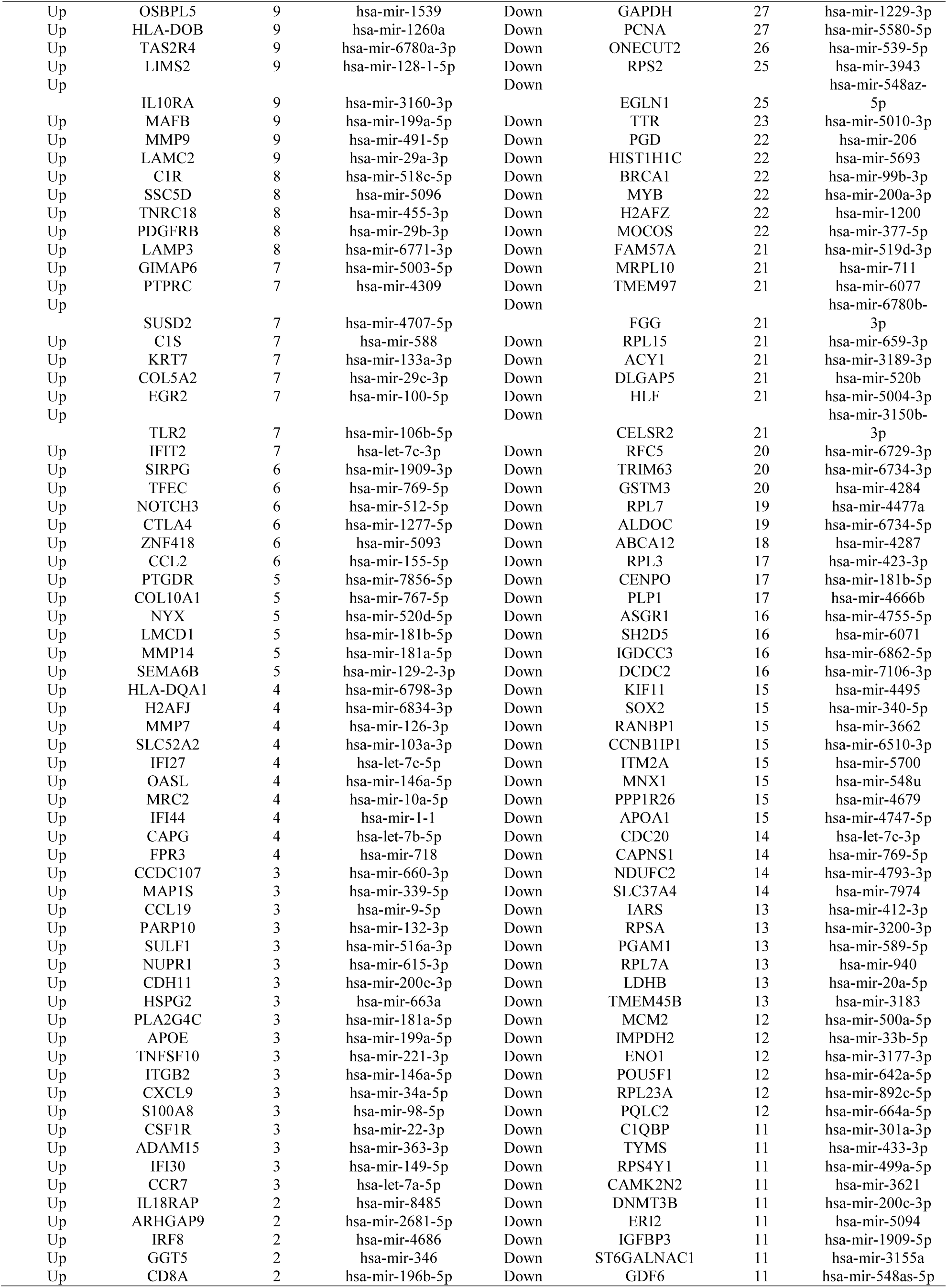
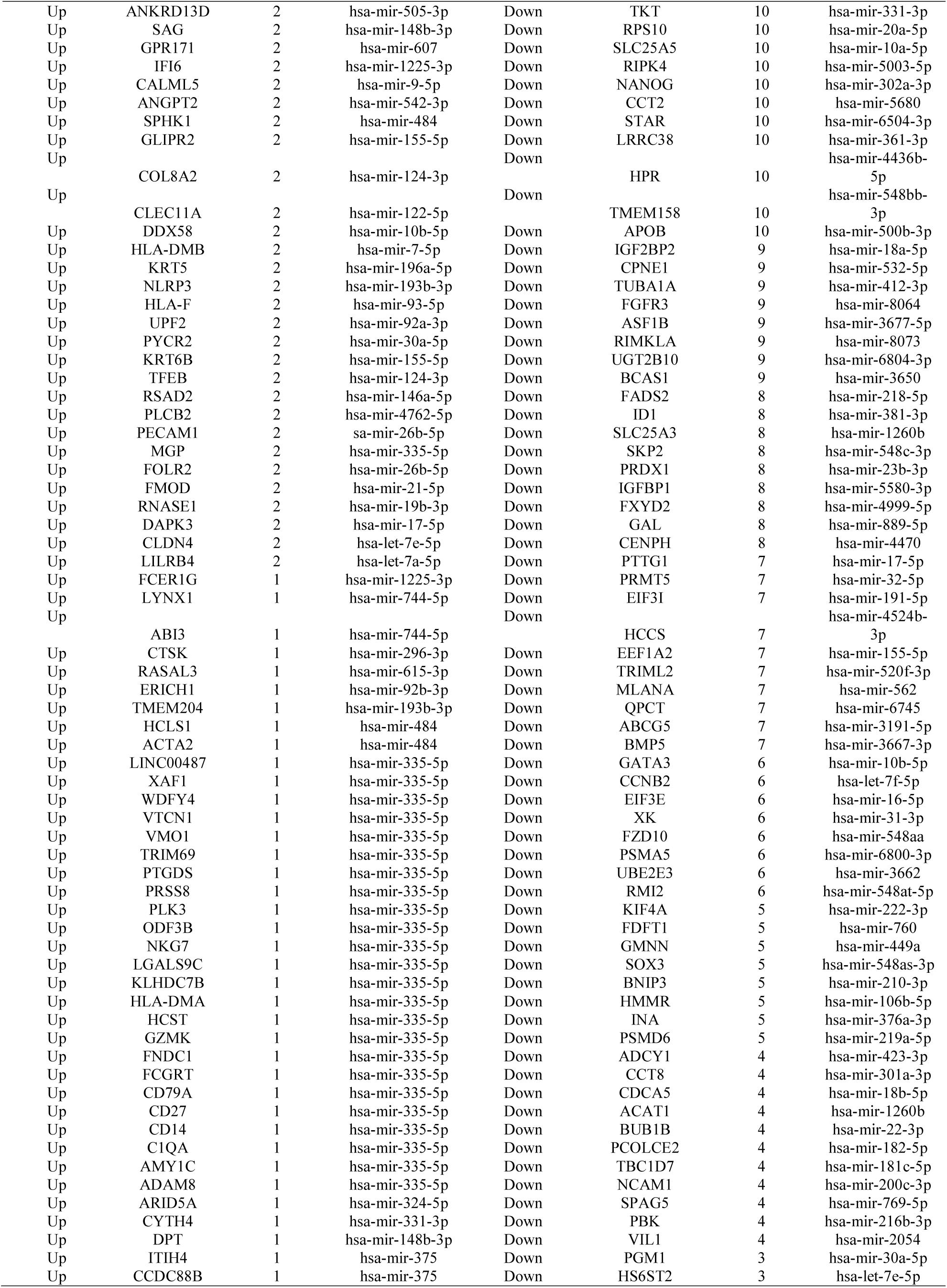
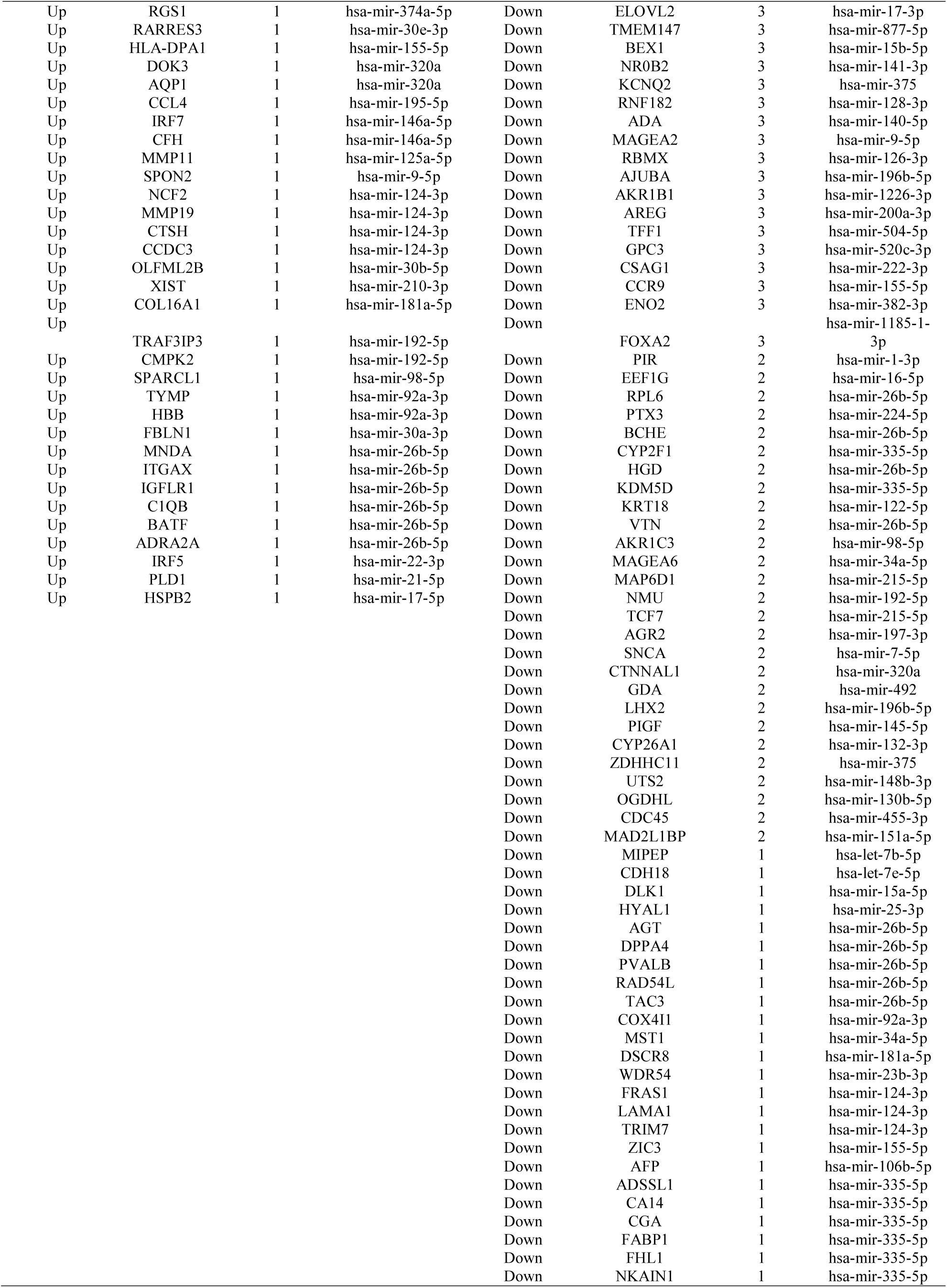
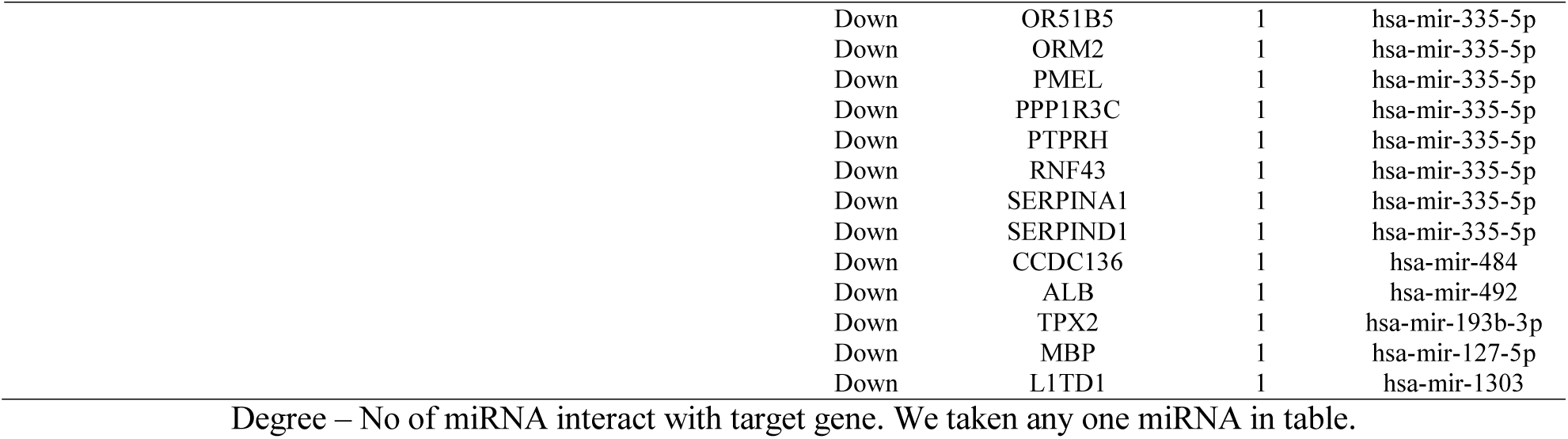
miRNA - target gene interaction table

### Construction of target gene - TF regulatory network

Expressions of TFs were important for progression of TNBC [Lawrence et al. 2015]. An up regulated target gene - TF regulatory network is shown in Fig. 13. Top five up regulated target genes such as HOPX interacts with 218 TFs (ex, FOXC1), NYX interacts with 163 TFs (ex, GATA2), CXCL9 interacts with 110 TFs (ex, YY1), ACTA2 interacts with 100 TFs (ex, FOXL1) and CYP1B1 interacts with 91 TFs (ex, PPARG) are listed in Table 9. Pathway and GO enrichment analysis displayed that the top up regulated targate genes were primarily enriched in the ensemble of genes encoding core extracellular matrix including ECM glycoproteins, collagens and proteoglycans, chemokine signaling pathway, inflammation mediated by chemokine and cytokine signaling pathway, and positive regulation of cell motility. Similarly, down regulated target gene - TF regulatory network is shown in Fig. 14. Top five down regulated target genes such as CCNA2 interacts with 207 TFs (ex, FOXC1), NR0B2 interacts with 167 TFs (ex, GATA2), CPS1 interacts with 109 TFs (ex, YY1), CCNB2 interacts with 93 TFs (ex, FOXL1) and STAR interacts with 87 TFs (ex, E2F1) are listed in Table 9. Pathway and GO enrichment analysis displayed that the top down regulated targate genes were primarily enriched in the cell cycle, mechanism of gene regulation by peroxisome proliferators via PPARa (alpha), biosynthesis of amino acids, mitotic cell cycle process and carboxylic acid metabolic process.

**Fig. 13.**
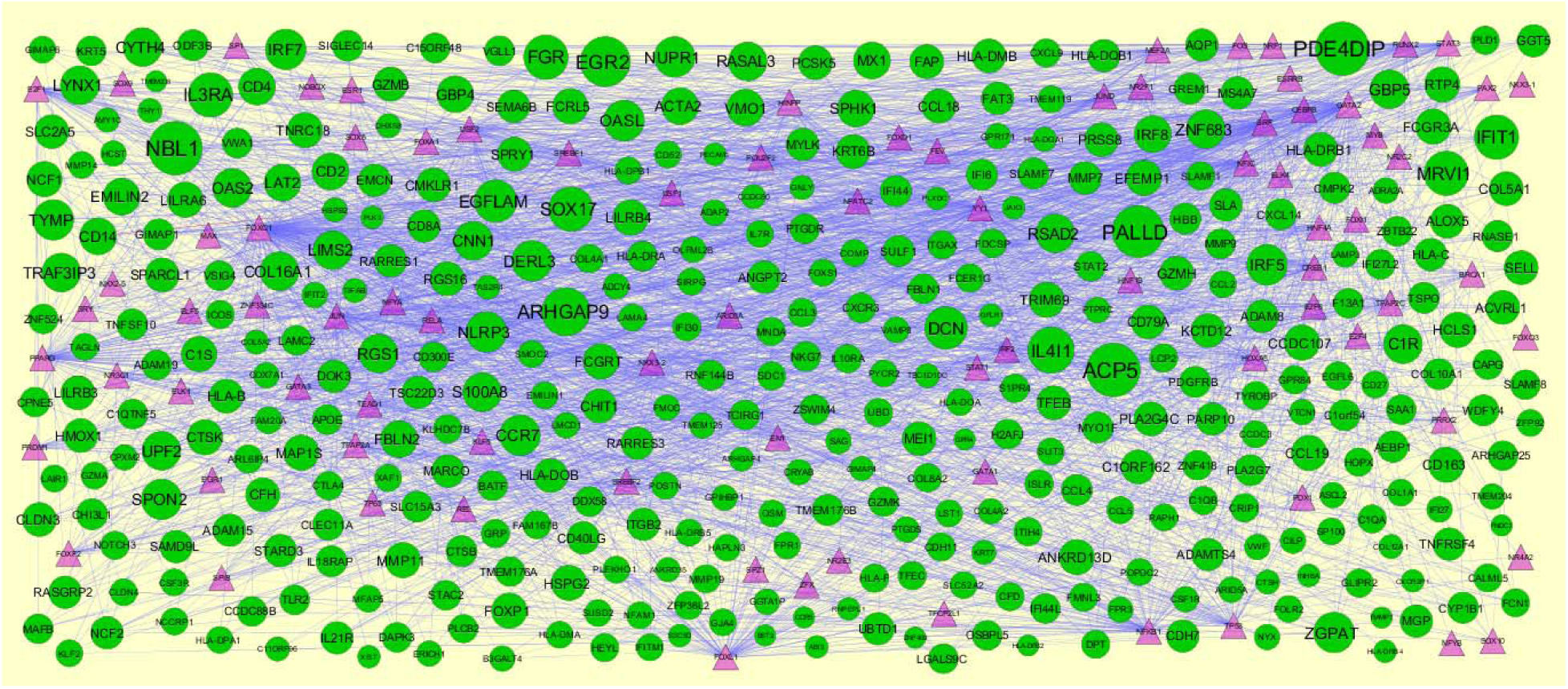
TF-gene network of predicted target up regulated genes. (Lavender triangle - TFs and green circles-target up regulated genes)

**Fig. 14.**
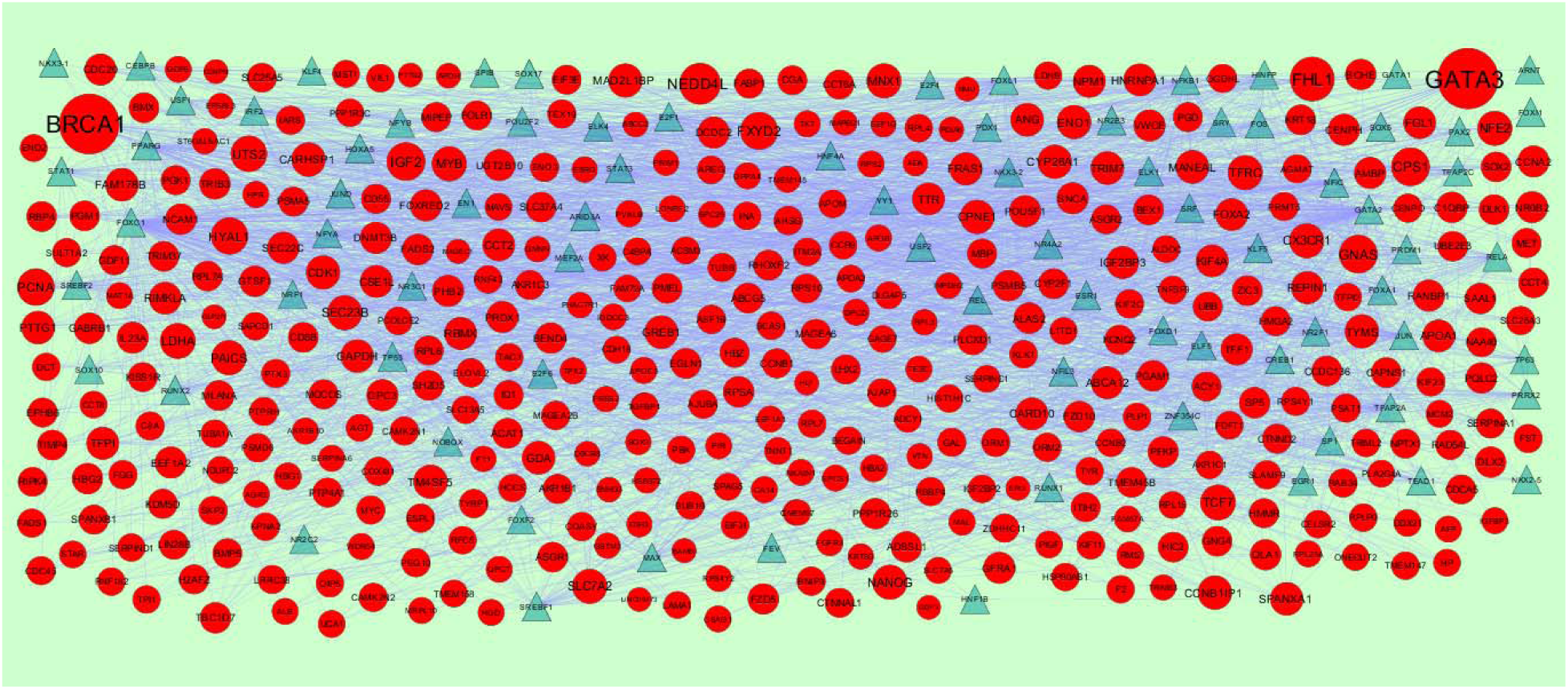
TF-gene network of predicted target down regulated genes. (Blue triangle - TFs and red circles-target up regulated genes)

**Table 9.**
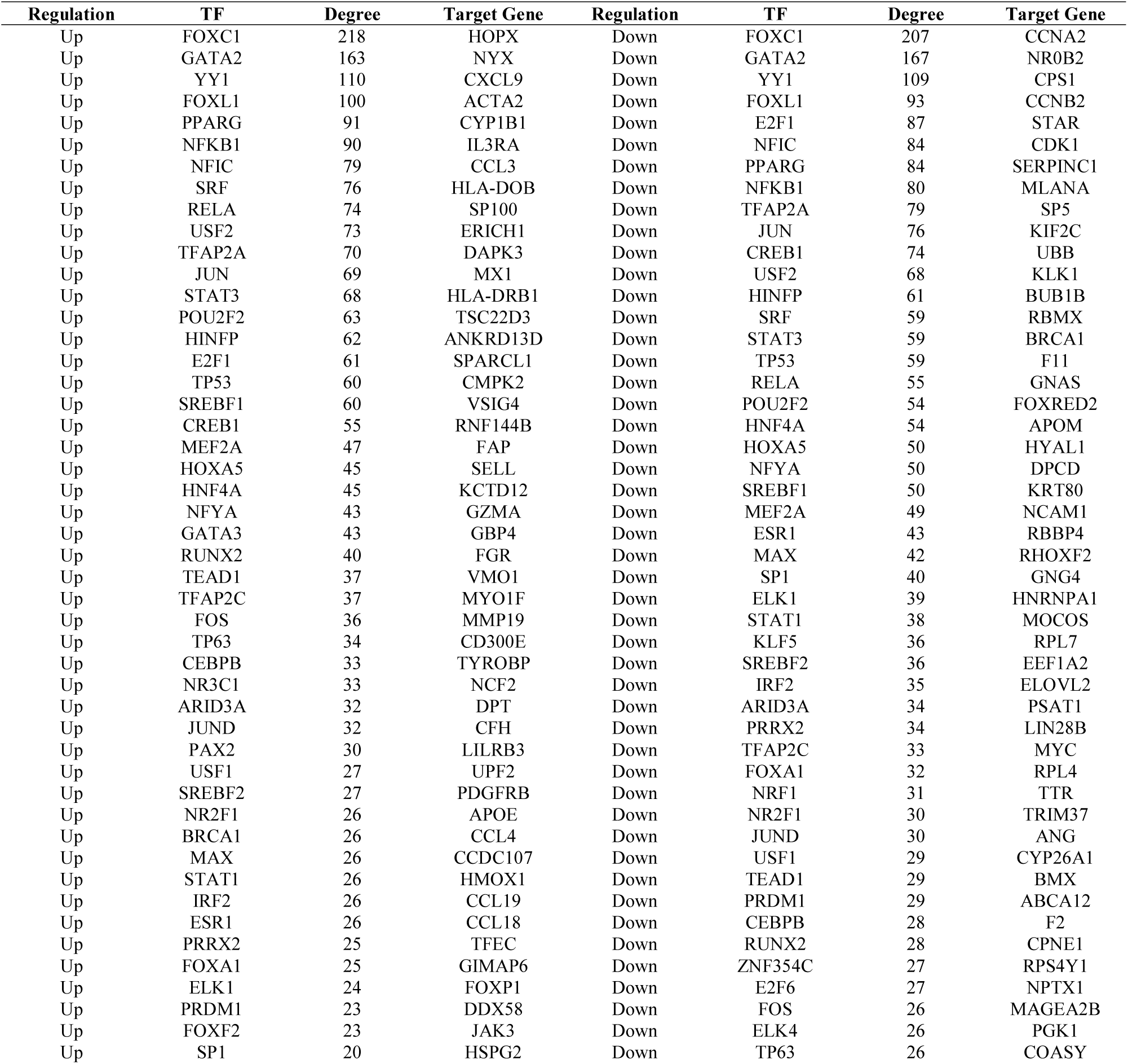
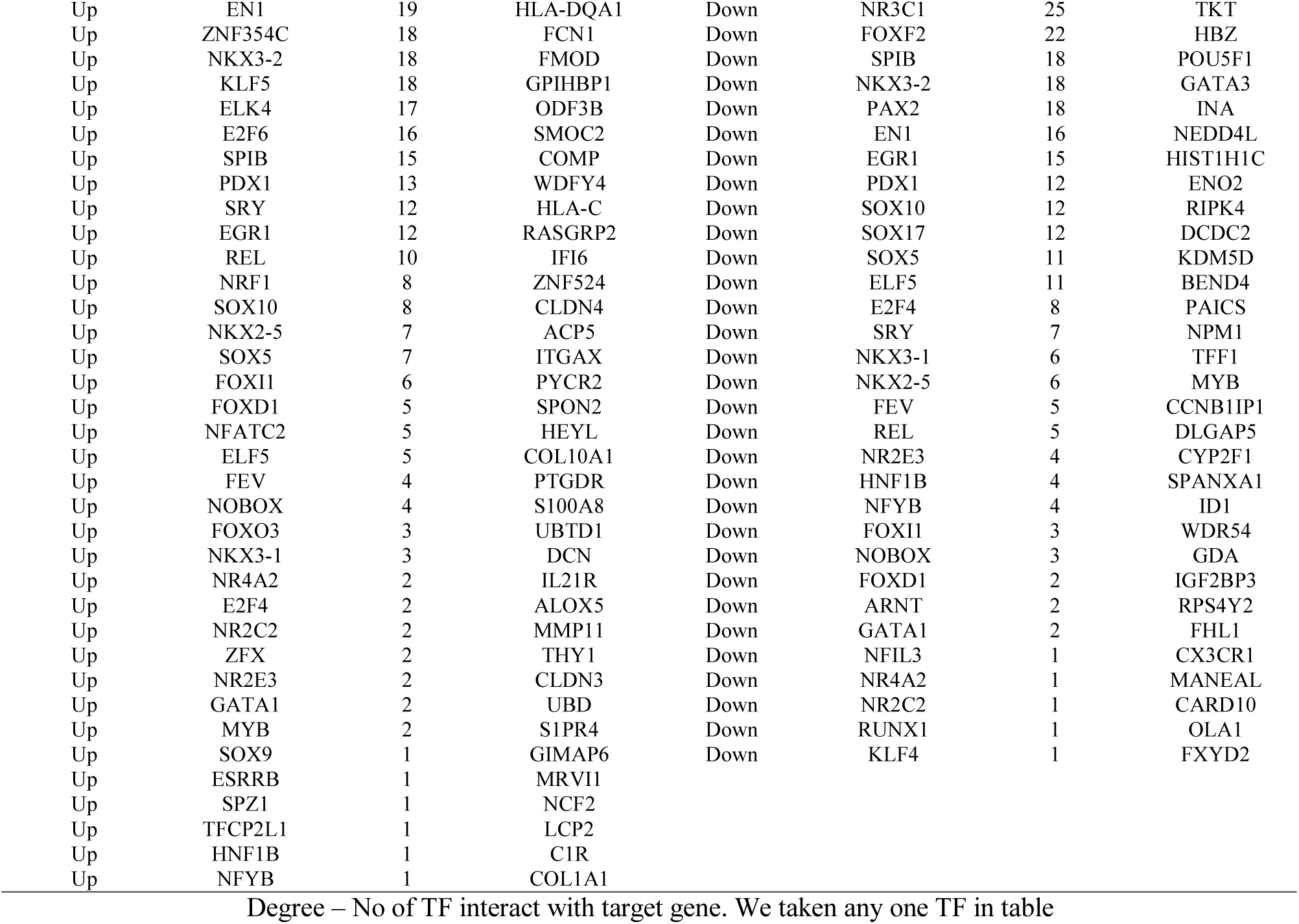
TF - target gene interaction table

### Validation of hub genes

The prognostic information of the 10 hub genes was available in the free online UALCAN database. As shown in Fig 15, the increase expression levels of the six hub genes such as ADAM15 (P = 0.0028), BATF (P = 0.002), NOTCH3 (P = 0.02), SDC1 (P = 0.00044), RBMX (P = 0.029) and ABCC2 (P = 0.0046) were associated with a worst OS for TNBC, while low expression levels of the four hub genes such ITGAX (P = 0.043), RPL4 (P = 0.048), EEF1G (P = 0.025) and RPL3 (P = 0.011) were associated with worst OS for TNBC (Fig. 16). Based on the UALCAN database, it was identified that the mRNA expression levels of ADAM15, BATF, NOTCH3, ITGAX and SDC1 were significantly increased in TNBC samples compared with normal breast samples (Fig. 17), while mRNA expression levels of RPL4, EEF1G, RPL3, RBMX and ABCC2 were significantly decreased in TNBC samples compared with normal breast samples (Fig. 18). In addition, UALCAN box plots of gene expression by pathological stages of breast cancer based on the TCGA database results shows that ADAM15, BATF, NOTCH3, ITGAX and SDC1 were more expressed in all stages of breast cancer compared to normal breast samples (Fig. 19), while RPL4, EEF1G, RPL3, RBMX and ABCC2 less expressed in all stages of breast cancer compared to normal breast samples (Fig. 20). We used cBioportal tool to explore the specific modification of hub genes. Percentages of modification in hub genes such as 10% ADAM15 (missense mutation, truncating mutation and amplification), 0.6% BATF (missense mutation, amplification and deep deletion), 3% NOTCH3 (missense mutation, truncating mutation, fusion and amplification), 5% ITGAX (missense mutation, truncating mutation and amplification), 0.4% SDC1 (truncating mutation and amplification), 0.7% RPL4 (missense mutation, truncating mutation and amplification), 1.1% EEF1G (missense mutation, truncating mutation and amplification), 0.5% RPL3 (fusion, amplification and deep deletion), 1.4% RBMX (missense mutation, truncating mutation, amplification and deep deletion) and 1.2 % ABCC2 (missense mutation, truncating mutation, fusion, amplification and deep deletion) (Fig.21). The Human Protein Atlas and IHC assays were performed to validate the expression level of hub genes. Meanwhile, IHC staining exhibited that the protein expression of ADAM15, BATF, NOTCH3, ITGAX and SDC1 were elevated in TNBC tissue compared with normal breast samples (Fig. 22A-22E), while protein expression of RPL4, EEF1G, RPL3, RBMX and ABCC2 were decreased in TNBC tissue compared with normal breast samples (Fig. 22F-22J) . In order to detect the discriminatory ability of common DEGs abovementioned among TNBC patients and healthy individuals, the ROC analyses in GSE88715 dataset was performed. The AUC under ROC curve were 0.887, 0.933, 0.824, 0.925, 0.833, 0.786, 0.876, 0.834, 0.861 and 0.918 for ADAM15, BATF, NOTCH3, ITGAX, SDC1, RPL4, EEF1G, RPL3, RBMX and ABCC2, respectively (Fig. 23). To validate these major results, we selectively performed RT-PCR analysis of ten hub genes including ADAM15, BATF, NOTCH3, ITGAX, SDC1, RPL4, EEF1G, RPL3, RBMX and ABCC2 in TNBC patients and healthy control. The relative mRNA expression levels of ADAM15, BATF, NOTCH3, ITGAX and SDC1 were significantly higher in TNBC patients when compared with those from healthy controls (Fig. 24A-E). The expression levels of RPL4, EEF1G, RPL3, RBMX and ABCC2 were significantly lower in TNBC patients than healthy control group (Fig. 24F-J). In the TNBC, the high expression levels of the ADAM15, BATF, NOTCH3, ITGA and SDC1 all negatively associated with tumor purity (Fig. 25 A-E), whereas low expression of RPL4, EEF1G, RPL3, RBMX and ABCC2 were all positively associated with tumor purity (Fig. 25 F-J). It can be inferred from this result that all ten hub genes are probably expressed not in the microenvironment, but expressed in the tumor cells.

**Fig. 15.**
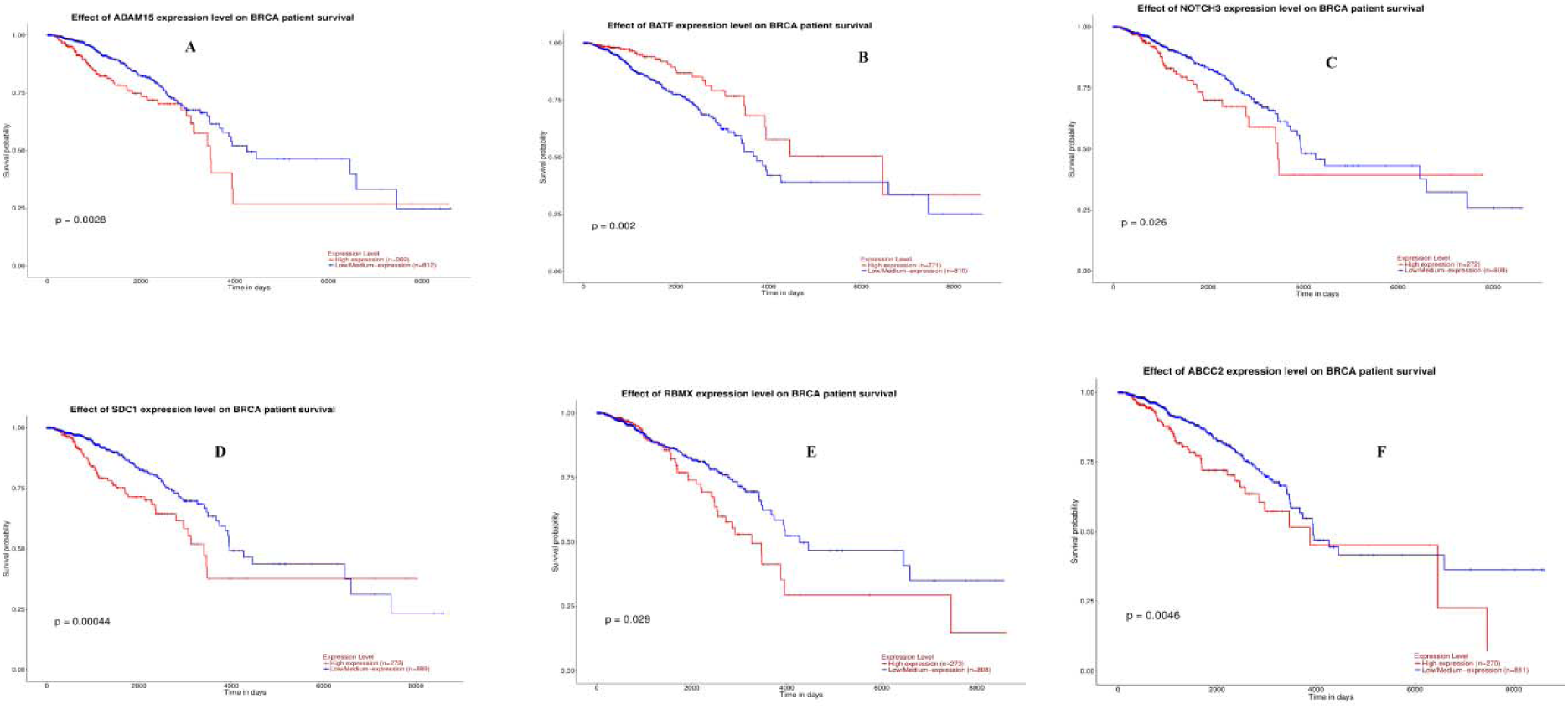
Overall survival analysis of hub genes . Overall survival analyses were performed using the UALCAN online platform. Red indicates - high expression group; Blue indicates – low expression group A) ADAM15 B) BATF C) NOTCH3 D) SDC1 E) RBMX F) ABCC2

**Fig. 16.**
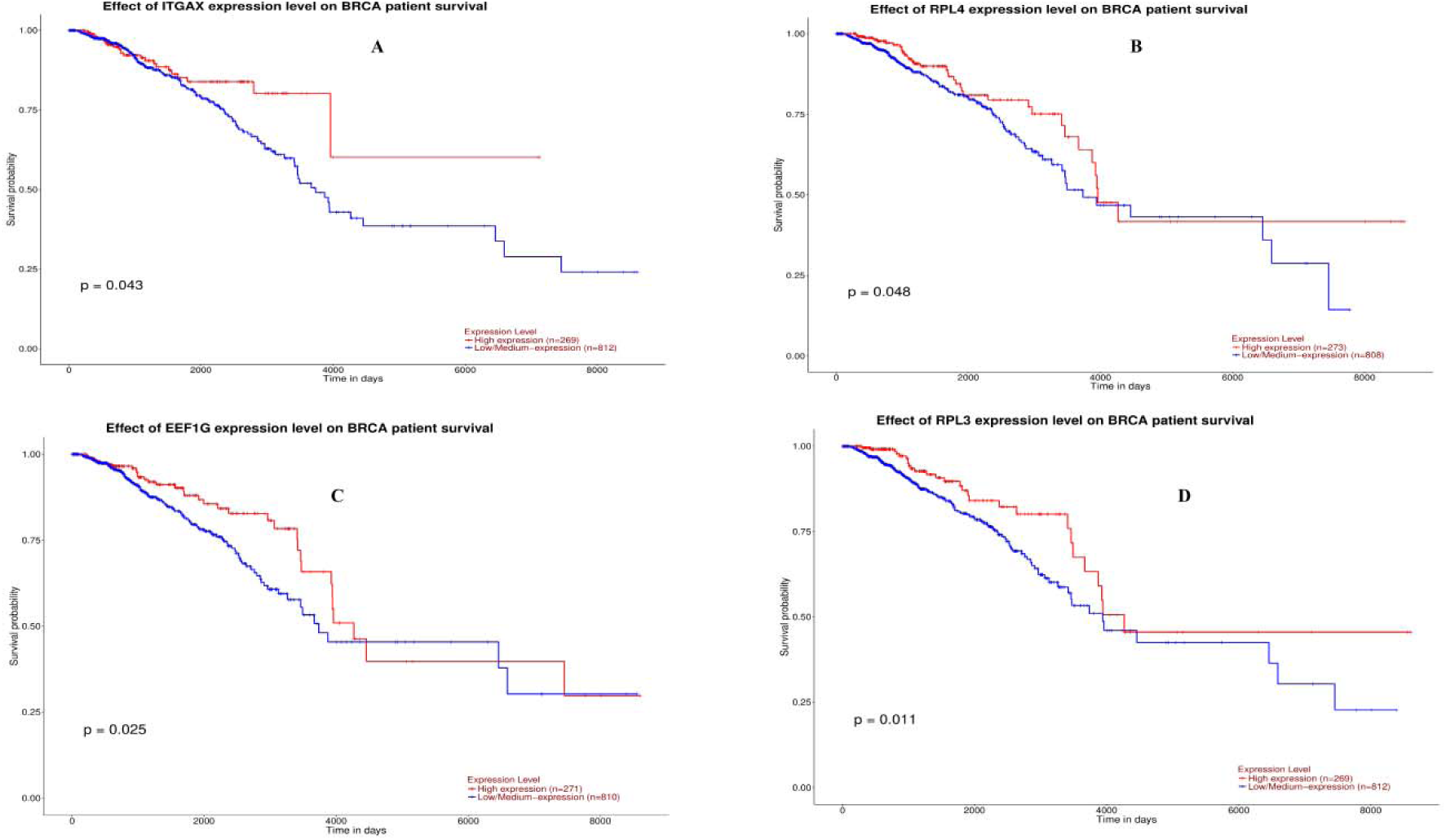
Overall survival analysis of hub genes. Overall survival analyses were performed using the UALCAN online platform. Red indicates - high expression group; Blue indicates – low expression group A) ITGAX B) RPL4 C) EEF1G D) RPL3

**Fig. 17.**
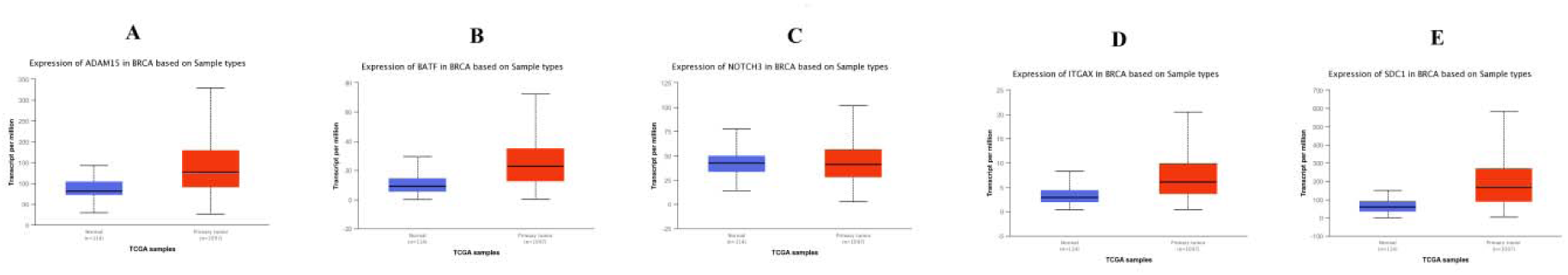
Box plots (expression analysis) hub genes (up regulated genes) were produced using the UALCAN platform. A) ADAM15 B) BATF C) NOTCH3 D) ITGAX E) SDC1

**Fig. 18.**
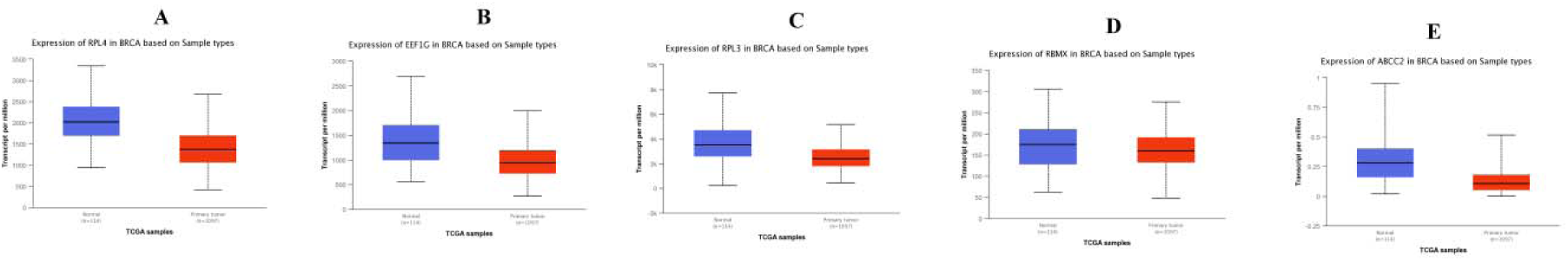
Box plots (expression analysis) hub genes (down regulated genes) were produced using the UALCAN platform. A) RPL4 B) EEF1G C) RPL3 D) RBMX E) ABCC2

**Fig. 19.**
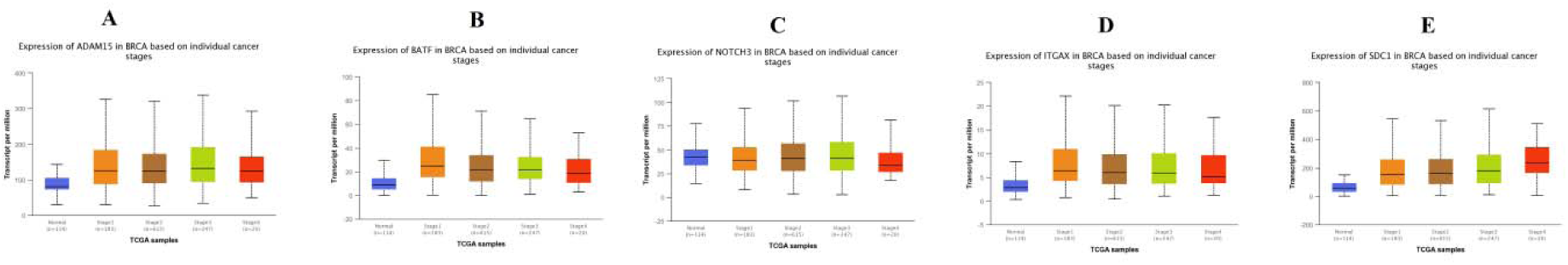
Box plots (stage analysis) of hub genes (up regulated) were produced using the UALCAN platform. A) ADAM15 B) BATF C) NOTCH3 D) ITGAX E) SDC1

**Fig. 20.**
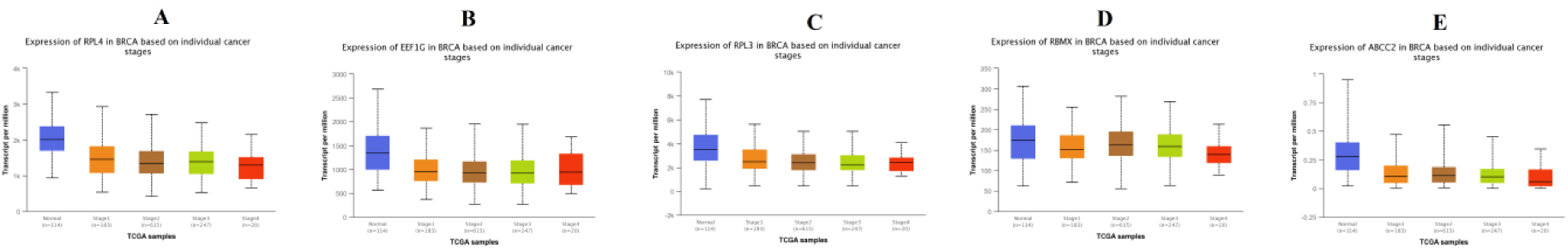
Box plots (stage analysis) of hub genes (down regulated) were produced using the UALCAN platform. A) RPL4 B) EEF1G C) RPL3 D) RBMX E) ABCC2

**Fig. 21.**
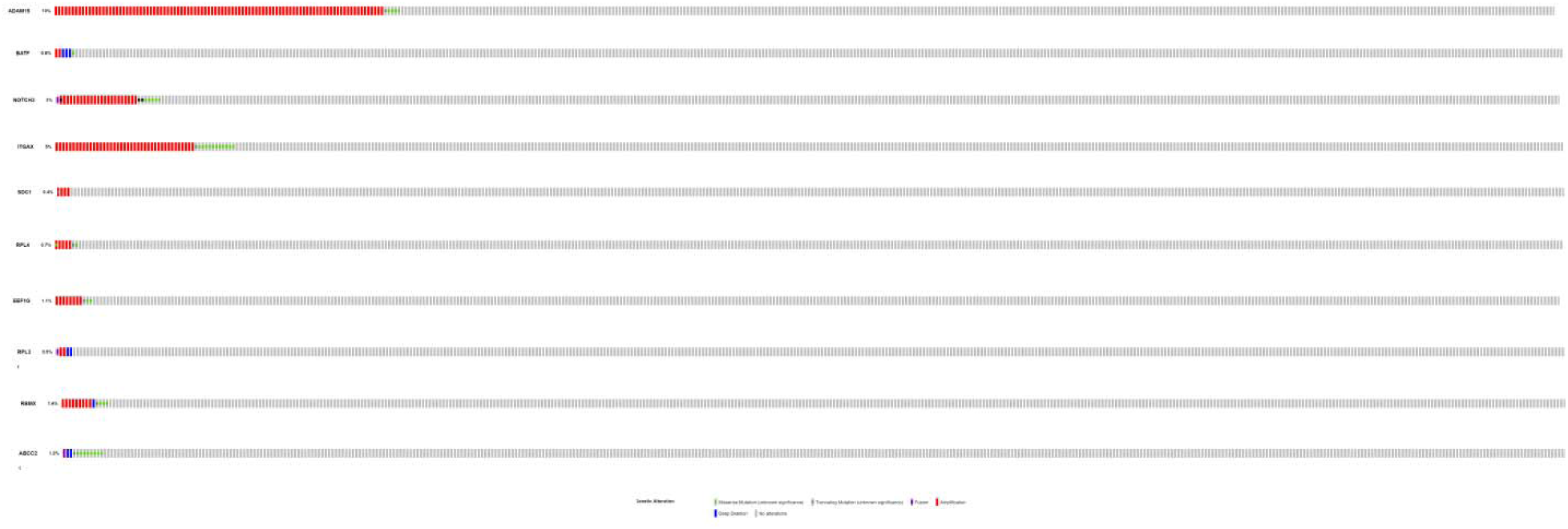
Mutation analyses of hub genes were produced using the CbioPortal online platform

**Fig. 22.**
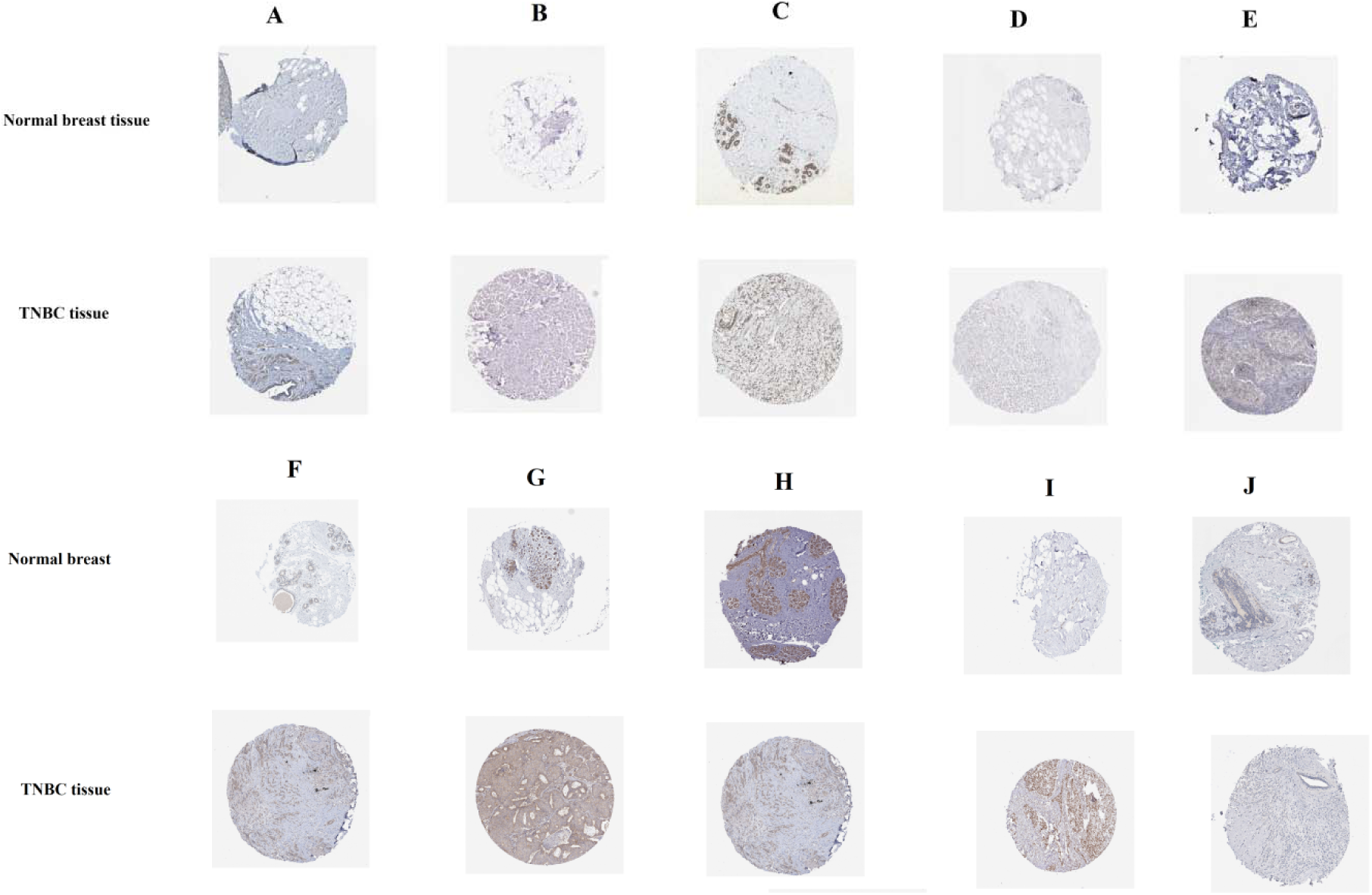
Immune histochemical analyses of hub genes were produced using the human protein atlas (HPA) online platform. A) ADAM15 B) BATF C) NOTCH3 D) ITGAX E) SDC1 F) RPL4 G) EEF1G H) RPL3 I) RBMX J) ABCC2

**Fig. 23.**
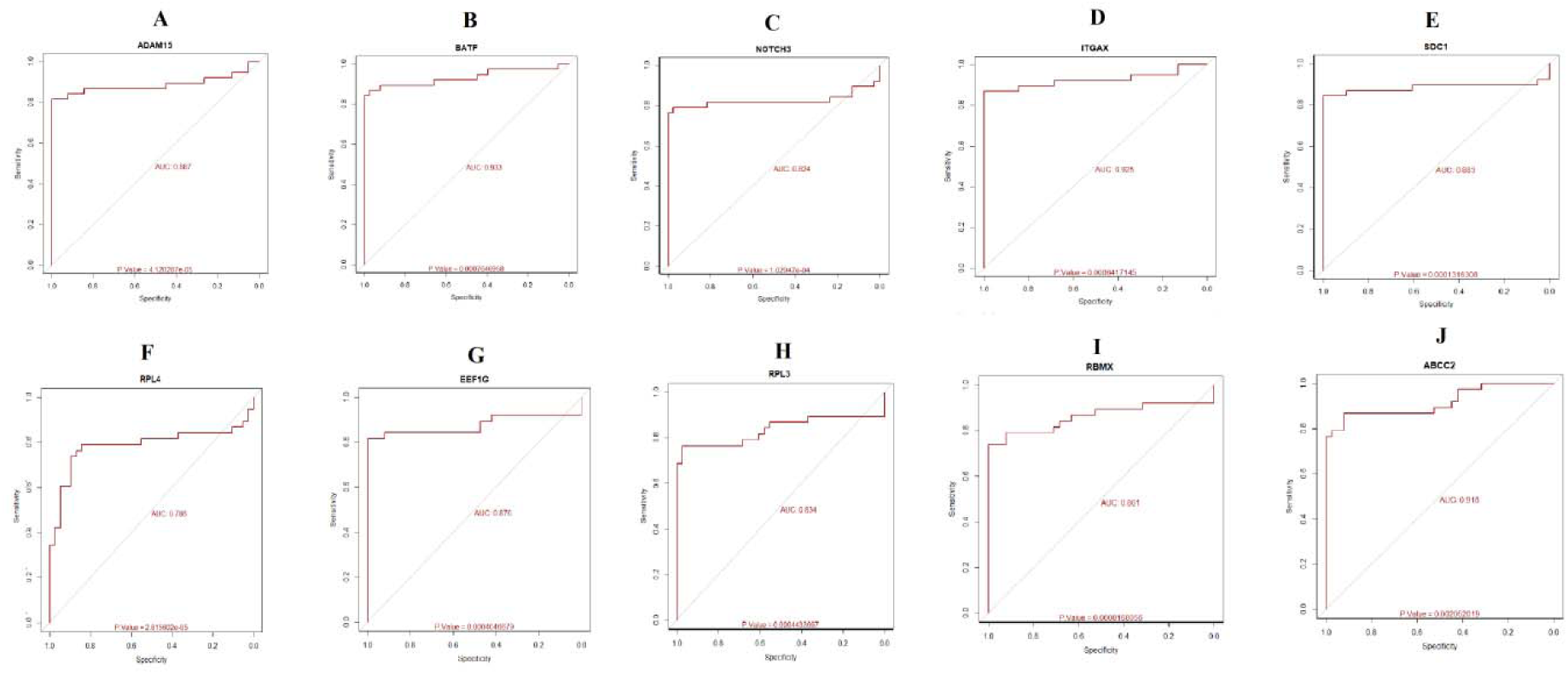
ROC curve validated the sensitivity, specificity of hub genes as a predictive biomarker for TNBC prognosis. A) ADAM15 B) BATF C) NOTCH3 D) ITGAX E) SDC1 F) RPL4 G) EEF1G H) RPL3 I) RBMX J) ABCC2

**Fig. 24.**
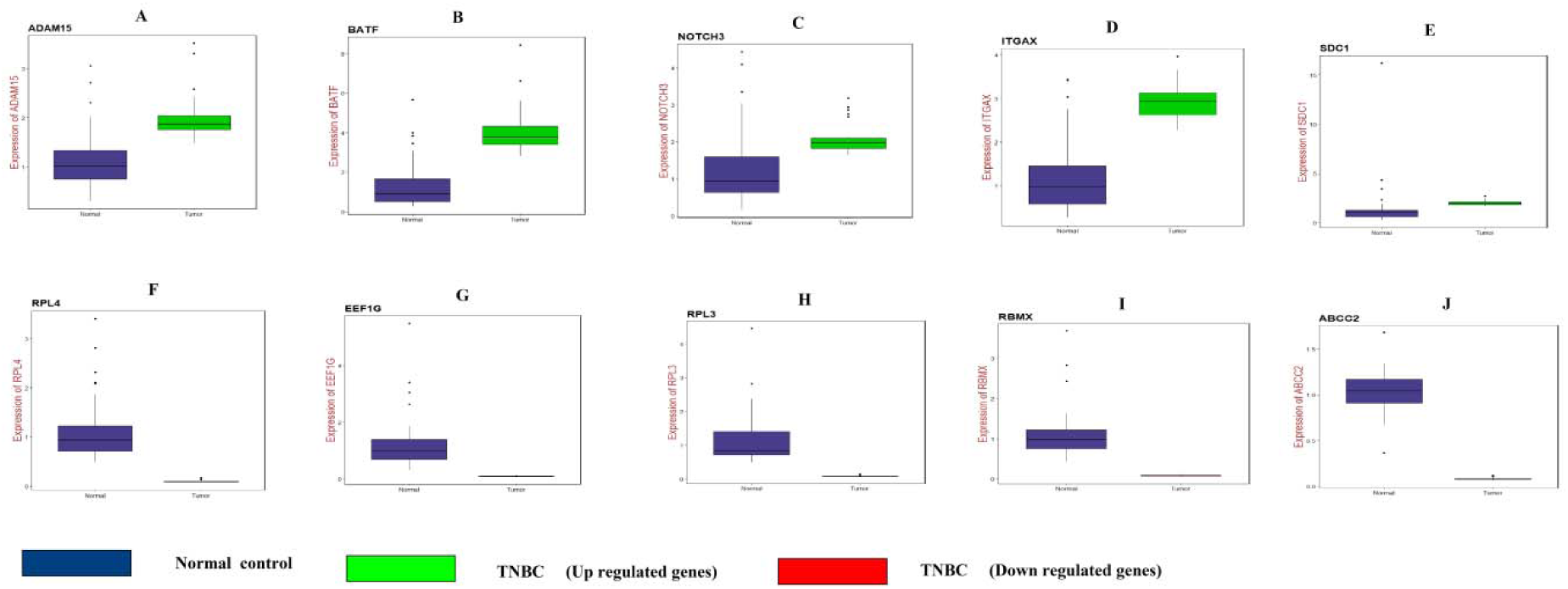
Validation of microarray data by quantitative PCR. A) ADAM15 B) BATF C) NOTCH3 D) ITGAX E) SDC1 F) RPL4 G) EEF1G H) RPL3 I) RBMX J) ABCC2

**Fig. 25.**
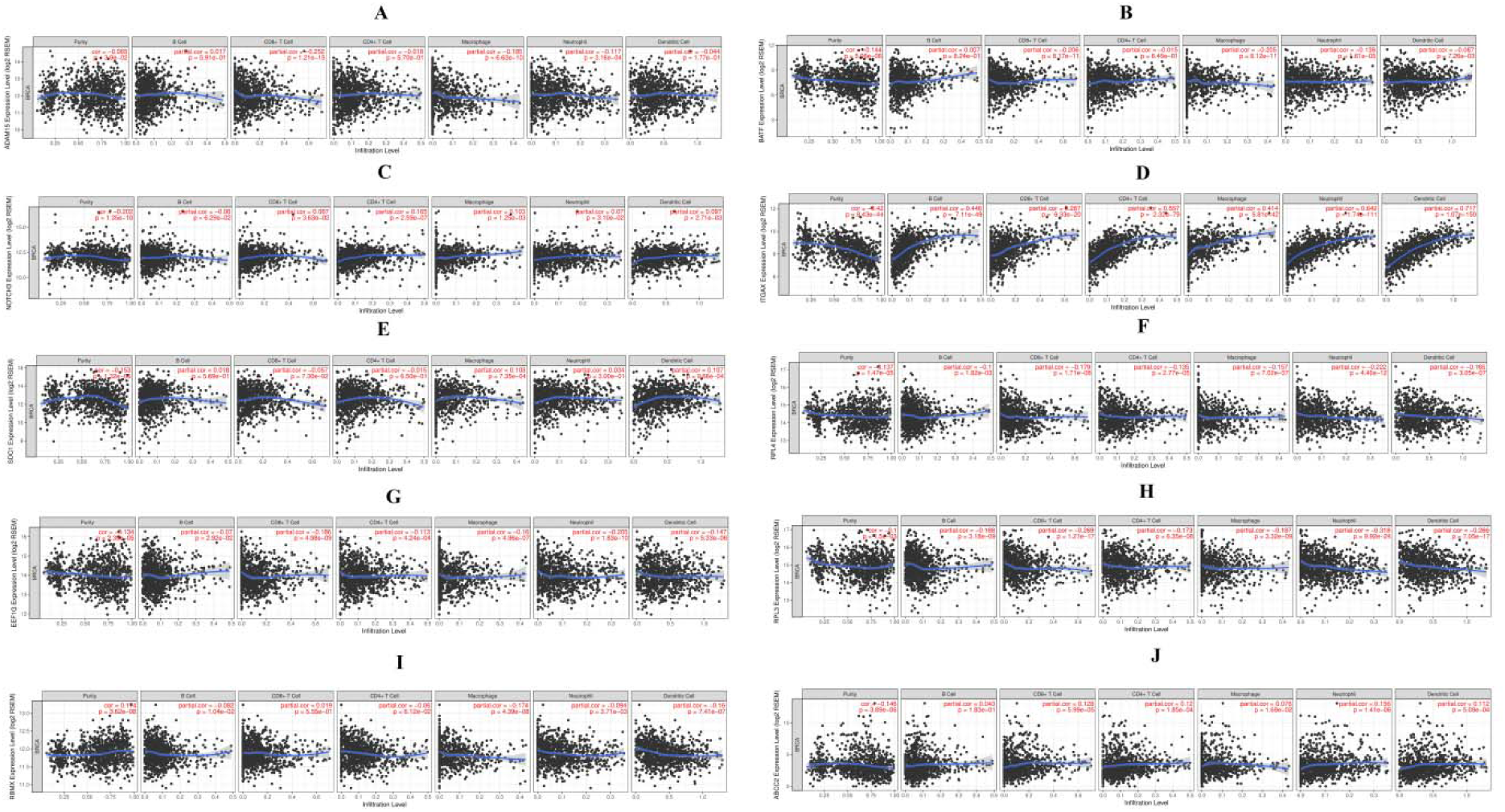
Scatter plot for Immune infiltration of A) ADAM15 B) BATF C) NOTCH3 D) ITGAX E) SDC1 F) RPL4 G) EEF1G H) RPL3 I) RBMX J) ABCC2 in breast cancer

## Discussion

Although many basic and clinical examinations have been conducted to illuminate the molecular origins of TNBC, the process has been slow and unsatisfying due to the absence of stable and effective biomarkers [Lehmann et al. 2011]. In this study, we diagnosed significant DEGs between TNBC (tumor stroma) and tissue samples from normal tissue (normal stroma). Furthermore, we implemented a series of bioinformatics analysis to screen important genes and pathways.

A total of 949 DEGs were found, consisting of 469 up regulated genes and 480 down regulated genes. CSF1R was associated with tumor-infiltrating macrophages in pancreatic cancer [Zhu et al. 2014], but this gene may be linked tumor-infiltrating macrophages in TNBC. Genes such as CD14 [Lobba et al. 2012], AHSG (alpha 2-HS glycoprotein) [Yi et al. 2009] and AFP (alpha fetoprotein) [Parikh et al. 2005] were responsible for pathogenesis of hormonal-receptor positive breast cancer, but these genes may be liable for progression of TNBC. Methylation inactivation of tumor suppressor PLEKHO1 was involved in progression of gastric cancer [Ma et al. 2016], but loss of this gene may be important for advancement of TNBC. Gene such as CCL4 [Sasaki et al. 2016], TYMP (thymidine phosphorylase) [Marangoni et al. 2018] and IGFBP1 [Zhang and Yee, 2000] were responsible for development of TNBC. ALB (albumin) was linked with advancement of colorectal cancer [Dixon et al. 2003], but this gene may be associated with pathogenesis of TNBC. FABP1 was liable for invasion of gastric cancer cells [Satoh et al. 2012], but this gene may be important for invasion of TNBC cells.

In pathway enrichment analysis, up regulated genes were enriched in numerous pathways from various pathway databases. Single nucleotide polymorphism (SNP) in enriched genes such as PLA2G4C [Olsen et al. 2016], HLA-DPB1 [Ivansson et al. 2011], HLA-DRB1 [Mahmoodi et al. 2012], C1QA [Azzato et al. 2010], CFH (complement factor H) [Zhang et al. 2012], CHIT1 [Li et al. 2016], FCGR3A [Gavin et al. 2017] and POSTN (periostin) [Försti et al. 2007] were important for pathogenesis of various cancers, but these polymorphic genes may be linked with progression of TNBC. High expression of enriched genes such as PLD1 [Gozgit et al. 2007], HLA-DRA (major histocompatibility complex, class II, DR alpha) [Rangel et al. 2004], IL10RA [Zadka et al. 2018], LAMC2 [Xu et al. 2017], TLR2 [Xin et al. 2014], GZMA (granzyme A) [Legrand et al. 2010], FCER1G [Chen et al. 2017], FGR (FGR proto-oncogene, Src family tyrosine kinase) [Kim et al. 2011], ICOS (inducible T cell costimulator) [Tiriveedhi et al. 2013], LAT2 [Barollo et al. 2016], SAA1 [Kim et al. 2015], VWF (von Willebrand factor) [Röhsig et al. 2001], EFEMP1 [Chen et al. 2013], FMOD (fibromodulin) [Dawoody Nejad et al. 2017] and FNDC1 [Das and Ogunwobi, 2017] were involved in advancement of various cancers, but elevated expression of these genes may be linked with development of TNBC. ALOX5 was linked with proliferation of colorectal cancer cells [Wang et al. 2018], but this gene may be responsible for proliferation of TNBC cells. Enriched gene such as CCR5 [Norton et al. 2017], LAMA4 [Yang et al. 2018], CD4 [Matsumoto et al. 2016], ADAM8 [Das et al. 2016], BST2 [Woodman et al. 2016], CTSB (cathepsin B) [Yang et al. 2017], FPR1 [Baracco et al. 2016], MMP9 [Mehner et al. 2014], S100A8 [Parris et al. 2014], COL4A2 [JingSong et al. 2017], DCN (decorin) [Yang et al. 2015], HAPLN3 [Kuo et al. 2010] and MFAP5 [Chen et al. 2019] were associated with pathogenesis of TNBC. Enriched genes such as CCL3 [Zhang et al. 2016], ADA2 [Aghaei et al. 2005], ARHGAP9 [Sun et al. 2019], CHI3L1 [Chen et al. 2017], CTSH (cathepsin H) [Jevnikar et al. 2013], CTSK (cathepsin K) [Kleer et al. 2008], DDX58 [Chang et al. 2017], IRF7 [Li et al. 2015], ITGB2 [Liu et al. 2018], JAK3 [Ye et al. 2013], MNDA (myeloid cell nuclear differentiation antigen) [Sun et al. 2014], NCF2 [Zhang et al. 2018], NLRP3 [Wang et al. 2016], PDGFRB (platelet derived growth factor receptor beta) [Schito et al. 2012], SLC2A5 [Weng et al. 2018], PTGDS (prostaglandin D2 synthase) [Shyu et al. 2013], AEBP1 [Liu et al. 2018], COL10A1 [Li et al. 2018], COL12A1 [Duan et al. 2018], COL1A1 [Zhang et al. 2018], COL4A1 [Huang et al. 2018], COL5A1 [Liu et al. 2018], COL5A2 [Zeng et al. 2018], COMP (cartilage oligomeric matrix protein) [Englund et al. 2016], DPT (dermatopontin) [Yamatoji et al. 2012], EMILIN2 [Marastoni et al. 2014],MGP (matrix Gla protein) [Tiago et al. 2016], SMOC2 [Shvab et al. 2016], SPARCL1 [Cao et al. 2013] and SPON2 [Chandrasinghe et al. 2017] were responsible for invasion of various cancer cells, but these genes may be liable for invasion of TNBC cells. IGHG1 was important for inhibition of apoptosis in prostate cancer [Pan et al. 2013], but this gene may be liable for inhibition of apoptosis in TNBC. Methylation inactivation of tumor suppressors enriched genes such as FBLN1 [Xu et al. 2015], SLIT3 [Kim et al. 2016] and LIMS2 [Kim et al. 2006] were important for progression of various cancers, but inactivation of these genes may be associated with development of TNBC. HSPG2 was involved in drug resistance in hormonal-receptor positive breast cancer [Hu et al. 2013], but this gene may be associated with chemo resistance in TNBC. Our study found that PLCB2, CD40LG, HLA-DMA (major histocompatibility complex, class II, DM alpha), HLA-DMB (major histocompatibility complex, class II, DM beta), HLA-DOA (major histocompatibility complex, class II, DO alpha), HLA-DOB (major histocompatibility complex, class II, DO beta), HLA-DPA1, HLA-DRB4, HLA-DRB5, CD8A, GZMB (granzyme B), IL18RAP, ADCY4, C1QB, C1R, C1S, CALML5, CD300E, CFD (complement factor D), DOK3, FCN1, GNLY (granulysin), GPR84, HBB (hemoglobin subunit beta), HLA-B (major histocompatibility complex, class I, B), HLA-C (major histocompatibility complex, class I, C), IL3RA, ITGAX (integrin subunit alpha X), LAIR1, LCP2, LILRB3, NCF1, NFAM1, PECAM1, PTPRC (protein tyrosine phosphatase, receptor type C), RASAL3, RASGRP2, SELL (selectin L), SIGLEC14, TBC1D10C, TCIRG1, TYROBP (TYRO protein tyrosine kinase binding protein), VAMP8, CILP (cartilage intermediate layer protein), COL16A1, COL8A2, EGFLAM (EGF like, fibronectin type III and laminin G domains), EMILIN1, FBLN2, NYX (nyctalopin), VWA1 and GGT5 are up regulated in TNBC and has potential as a novel diagnostic and prognostic biomarker, and therapeutic target. Similarly, down regulated genes were enriched in numerous pathways from various pathway databases. Enriched genes such as ALDOC (aldolase, fructose-bisphosphate C) [Li et al. 2016], CPS1 [Zhang et al. 2018], ENO1 [Principe et al. 2017], TKT (transketolase) [Tseng et al. 2018], TFRC (transferrin receptor) [Wada et al. 2006] and RPL7A [Zhu et al. 2001] were involved in invasion of various cancer cells, but these genes may be liable for invasion of TNBC cells. Enriched genes such as ENO2 [Soh et al. 2011], ACY1 [Cook et al. 1993], APOA1 [Cine et al. 2014] and RPL7 [Boleij et al. 2010] were associated with pathogenesis of various cancers. Enriched genes such as PFKP (phosphofructokinase, platelet) [Moon et al. 2011] and PGK1 [Shashni et al. 2013] were linked with proliferation of hormonal-receptor positive breast cancer cells, but these genes may be liable for proliferation of TNBC cells. Enriched genes such as GAPDH (glyceraldehyde-3-phosphate dehydrogenase) [Higashimura et al. 2011], PGAM1 [Peng et al. 2016] and RPS2 [Wang et al. 2009] were important for pathogenesis of various cancers, but these genes may be identified with progression of TNBC. SNP in MAT1A was linked with advancement of hormonal-receptor positive breast cancer [Mumbrekar et al. 2017], but this polymorphic gene may be important for progression of TNBC. Enriched genes such as PSAT1 [Gao et al. 2017], DLK1 [Nueda et al. 2017], FOXA2 [Perez-Balaguer et al. 2015], LDHA (lactate dehydrogenase A) [Dong et al. 2017] and LDHB (lactate dehydrogenase B) [McCleland et al. 2012] were associated with advancement of TNBC. Low expression of enriched genes such as TTR (transthyretin) [Liu et al. 2007], RPL15 [Yan et al. 2015], RPL6 [Wu et al. 2011], RPLP0 [Teller et al. 2015] and RPSA (ribosomal protein SA) [Song et al. 2013] were liable for advancement of various cancers, but less expression of these genes may be identified with growth of TNBC. Enriched genes such as EEF1A1 [Lin et al. 2018] and RPL3 [Pagliara et al. 2016] were associated with control of cell cycle in various cancers, but these genes may be linked with control cell cycle in TNBC. Our study found that ENO3, TPI1, F2 (coagulation factor II, thrombin), RPL23A, RPL4, RPS10, RPS4Y1, RPS4Y2 and PGM1 are down regulated in TNBC and has potential as a novel diagnostic and prognostic biomarker, and therapeutic target.

In GO enrichment analysis, up regulated genes were enriched in all GO categories. Enriched genes such as CXCL9 [Tokunaga et al. 2018], CCL18 [Wang et al. 2016], IFI6 [Cheriyath et al. 2018], VTCN1 [Gao et al. 2015], ADAM19 [Zhang et al. 2015], MRC2 [Gai et al. 2014], CXCR3 [Bronger et al. 2012] and MMP19 [Djonov et al. 2001] were important invasion of various cancer cells, but these genes may be associated with invasion of TNBC cells. SNP in enriched genes such as HLA-DQA1 [Spraggs et al. 2011], HLA-DQB1 [Madeleine et al. 2008], IGKC (immunoglobulin kappa constant) [Pandey et al. 2014], TNFSF10 [Charbonneau et al. 2014], ADAM15 [Maretzky et al. 2009], CD27 [Xu et al. 2012], APOE (apolipoprotein E) [Saadat and Apolipoprotein, 2012] and SULF1 [Zhou et al. 2016] were liable for pathogenesis of various cancers, but these polymorphic genes may be identified with progression of TNBC. VSIG4 was inhibiting T-cell activation results in progression of lung cancer [Liao et al. 2014], but this gene may be inhibiting T-cell activation in TNBC. Enriched genes such as HMOX1 [Zhu et al. 2015], LAMP3 [Nagelkerke et al. 2014] and MX1 [Bajo et al. 2003] were liable for chemo resistance in hormonal-receptor positive breast cancer, but these genes may be associated with drug resistance in TNBC. Enriched genes such as CCL2 [Fang et al. 2016], CCL5 [Araujo et al. 2018], CCR7 [Li et al. 2017], MMP7 [Han et al. 2018], UBD (ubiquitin D) [Han et al. 2015], IFITM1 [Ogony et al. 2016], CXCL14 [Sjöberg et al. 2016], IFIT2 [Koh et al. 2019], IFIT1 [Danish et al. 2013], CTLA4 [Liu et al. 2018], IRF5 [Pimenta et al. 2015], STAT2 [Ogony et al. 2016], OSM (oncostatin M) [Bottai et al. 2017], SUSD2 [Hultgren et al. 2017], SDC1 [Cui et al. 2017], CRYAB (crystallin alpha B) [Chen et al. 2014], MYLK (myosin light chain kinase) [Choi et al. 2016], MMP11 [Kwon et al. 2011] and MMP14 [Ling et al. 2017] were responsible for advancement of TNBC. Over expression of genes such as CCL19 [Zhang et al. 2017], IGHA2 [Kang et al. 2012], TFEB (transcription factor EB) [Giatromanolaki et al. 2017], THY1 [Lu et al. 2011], RAMP1 [Logan et al. 2013], ACVRL1 [Chien et al. 2013] and CPXM2 [Niu et al. 2019] were involved in pathogenesis of various cancers, but high expression of these genes may be liable for advancement of TNBC. IRF8 was associated with immune surveillance in hormonal-receptor positive breast cancer and pancreatic cancer [Meyer et al. 2018], but this gene may be responsible for immune surveillance in TNBC. Enriched genes such as CRIP1 [Ludyga et al. 2013], XAF1 [Iravani et al. 2017], IFI44L [Huang et al. 2018], GZMH (granzyme H) [Tahbaz-Lahafi et al. 2014] and PRSS8 [Zhang et al. 2016] were important for development of various cancers, but low expression of these genes may be associated with development of TNBC. Genes such as MYO1F [Diquigiovanni et al. 2018], FOXP1 [Chiang et al. 2017], DAPK3 [Tan et al. 2017], FAP (fibroblast activation protein alpha) [Jia et al. 2014] and FOLR2 [Xu et al. 2018] were linked with proliferation of various cancer cells, but these genes may be associated with proliferation of TNBC cells. Methylation inactivation of tumarsupressor GREM1 was associated with advancement of renal cancer [Vlodrop et al. 2017], but loss of this gene may be linked with progression of TNBC. Our study found that HLA-F (major histocompatibility complex, class I, F), TRAC (T cell receptor alpha constant), TNFRSF4, FCGRT (Fc fragment of IgG receptor and transporter), IFI30, CMKLR1, DHX58, BATF (basic leucine zipper ATF-like transcription factor), LILRA6, ACKR4, IFI27, SLAMF1, RSAD2, IGHA1, IGHM (immunoglobulin heavy constant mu), OASL (2’-5’-oligoadenylate synthetase like), FCRL5, SLAMF7, MARCO (macrophage receptor with collagenous structure), IL7R, S1PR4, SP100, JAML (junction adhesion molecule like), HCST (hematopoietic cell signal transducer), SSC5D, GBP4, GBP5, LILRB4, LST1, OAS2, RGS1, CD79A, ADGRF5, GPIHBP1, GGTA1P, CD2, PCSK5 and GZMK (granzyme K) are up regulated in TNBC and has potential as a novel diagnostic and prognostic biomarker, and therapeutic target. Similarly, down regulated genes were enriched in all GO categories. Enriched genes such as ACAT1 [Antalis et al. 2010], PGD (phosphogluconate dehydrogenase) [Yang et al. 2018], FOLR1 [Necela et al. 2015], EGLN1 [Briggs et al. 2016], TRIB3 [Wennemers et al. 2011], MYC (MYC proto-oncogene, bHLH transcription factor) [Camarda et al. 2016], PSMB5 [Wei et al. 2018], GATA3 [Krings et al. 2014], AKR1C1 [Wenners et al. 2016], BRCA1 [Greenup et al. 2013], SLC7A5 [El Ansari et al. 2018], MET (MET proto-oncogene, receptor tyrosine kinase) [Mueller et al. 2012], HP (haptoglobin) [Tabassum et al. 2012], AGR2 [Ondrouskova et al. 2017], MST1 [Ercolani et al. 2017], IGFBP3 [Marzec et al. 2015], GFRA1 [He et al. 2017], SERPINA6 [Ronde et al. 2013], TIMP4 [Liss et al. 2009] and GDF11 [Bajikar et al. 2017] were linked with progression of TNBC. Increase expression of genes such as TYMS (thymidylatesynthetase) [Lee et al. 2013], ELOVL2 [González-Bengtsson et al. 2016], ADA (adenosine deaminase) [Aghaei et al. 2010], SERPINA1 [Chan et al. 2015], FGG (fibrinogen gamma chain) [Zhu et al. 2009], AKR1B1 [Reddy et al. 2017], AMBP (alpha-1-microglobulin/bikunin precursor) [Huang et al. 2013], VTN (vitronectin) [Niu et al. 2016], HYAL1 [Tan et al. 2011], AREG (amphiregulin) [Wang et al. 2008], C1QBP [Amamoto et al. 2011], C4BPA [Sogawa et al. 2016], RBP4 [Fei et al. 2017], ORM2 [Zhang et al. 2012], PRDX1 [O’Leary et al. 2014], PTTG1 [Ghayad et al. 2009], CD55 [Cui et al. 2012], ITIH3 [Chong et al. 2010] and NPM1 [Zhu et al. 2015] were responsible for development of various cancers, but high expression of these genes may be liable for development of TNBC. Genes such as TYRP1 [Agarwal et al. 2003], CYP26A1 [Osanai and Lee, 2015], STAR (steroidogenic acute regulatory protein) [Moog-Lutz et al. 1997], HPR (haptoglobin-related protein) [Shurbaji et al. 1991], ANG (angiogenin) [Dutta et al. 2014], IGF2 [Gebeshuber and Martinez, 2013], GPC3 [Castillo et al. 2016], PTX3 [Chang et al. 2015], TFF1 [Prest et al. 2002], TFPI (tissue factor pathway inhibitor) [Sierko et al. 2015], RACK1 [Cao et al. 2011], PHACTR1 [Fils-Aimé et al. 2013] and TESC (tescalcin) [Kang et al. 2016] were important for invasion of various cancer cells, but these genes may be identified with invasion of TNBC cells. Decrease expression of enriched genes such as ACSM3 [Ruan et al. 2017], PLA2G4A [Gosenca et al. 2012], OGDHL (oxoglutarate dehydrogenase like) [Fedorova et al. 2015], DNMT3B [Roscigno et al. 2016], GDF3 [Li et al. 2012], APOM (apolipoprotein M) [Hu et al. 2015] and TFPI2 [Xu et al. 2013] were associated with advancement of various cancers, but low expression of these genes may be liable for pathogenesis of TNBC. SNP in enriched genes such AGT (angiotensinogen) [González-Zuloeta et al. 2007], APOC3 [Launonen et al. 1999], FADS1 [Chen et al. 2017], SELENOP (selenoprotein P) [Ekoue et al. 2017], APOB (apolipoprotein B) [Liu et al. 2013], BCHE (butyrylcholinesterase) [Boberg et al. 2013], BMP5 [Huang et al. 2012] and UTS2 [Yumrutas et al. 2015] were involved in development of various cancers, but these polymorphic genes may be linked with development of TNBC. AKR1C3 was linked with drug resistance in hormonal-receptor positive breast cancer [Zhong et al. 2015], but this gene may be associated with chemo resistance in TNBC. LAMA1 was associated with proliferation of gastric cancer cells [Meng et al. 2016], but this gene may be involved in proliferation of TNBC cells. Methylation inactivation of tumor suppressor CAMK2N1 was liable for progression of ovarian cancer [Häfner et al. 2016], but loss of this gene may be liable for pathogenesis of TNBC. Our study found that HGD (homogentisate 1,2-dioxygenase), TYR (tyrosinase), UGT2B10, FADS2, PLP1, IARS (isoleucyl-tRNAsynthetase), ERO1A, APOA2, SLC7A2, SNCA (synuclein alpha), SLC37A4, CYP2F1, PSMA5, DCT (dopachrometautomerase), PSMD6, ADSSL1, AGMAT (agmatinase), GDF6, F11 (coagulation factor XI), UBB (ubiquitin B), FGL1, APOH (apolipoprotein H), IL23A, SERPINC1, PRSS2, SAAL1, TNFSF9, ITIH2, RBMX (RNA binding motif protein X-linked), TAC3, C8A, GAL (galanin and GMAP prepropeptide), ORM1, INA (internexin neuronal intermediate filament protein alpha), HBA2, HBG2, SERPIND1, VIL1, PTTG2, PPP1R26 and CAMK2N2 are down regulated in TNBC and has potential as a novel diagnostic and prognostic biomarker, and therapeutic target.

In PPI network for up regulated genes were identified as hub genes showing with highest node degree, betweenness, stress, closeness, and low clustering coefficient. DERL3 was associated with pathogenesis of TNBC [Shibata et al. 2017]. Our study found that IGFLR1 is up regulated in TNBC and has potential as a novel diagnostic and prognostic biomarker, and therapeutic target. Similarly, in PPI network for down regulated genes were identified as hub genes showing the highest node degree, betweenness, stress, closeness, and low clustering coefficient. Genes such as HSP90AB1 [Cheng et al. 2012] and PCNA [Yu et al. 2013] were responsible for progression of TNBC. HNRNPA1 important for invasion of hormonal-receptor positive breast cancer cells [Loh et al. 2015], but this gene may be liable for invasion of TNBC cells.

In module analysis, up regulated genes were identified as hub genes showing the highest node degree in all four significant modules. SLA was identified as novel biomarker for pathogenesis of TNBC. Similarly, down regulated genes were identified as hub genes showing the highest node degree in all four significant modules. Genes such as CCT2 [Guest et al. 2015], CCT8 [Qiu et al. 2015], CCNA2 [Gao et al. 2014] and CCNB1 [Ding et al. 2014] were involved in proliferation of various cancer cells, but these genes may be linked with proliferation of TNBC cells. Genes such as CCT6A [Huang et al. 2019], CDK1 [Liu et al. 2014], SKP2 [Huang et al. 2015] and CDC20 [Karra et al. 2014] were important for advancement of TNBC. Low expression of genes such as EIF3E [Gillis and Lewis, 2013] and EIF3I [Lin et al. 2014] were liable for development of various cancers, but less expression of these genes may be assoited with progression of TNBC. Our study found that CCT4 and TUBA1B are down regulated in TNBC and has potential as a novel diagnostic and prognostic biomarker, and therapeutic target.

In target gene - miRNA network, up regulated genes were identified as target genes showing the highest number of integration with miRNAs. Decrease expression of CCDC80 was involved in development of thyroid cancer [Ferraro et al. 2013], but less expression of this gene may be liable for progression of TNBC. Methylation inactivation KLF2 was important for development gastric cancer [Nie et al. 2017], but loss this gene may be associated with progression of TNBC. Our study found that CDH7 is up regulated in TNBC and has potential as a novel diagnostic and prognostic biomarker, and therapeutic target. Similarly, down regulated genes were identified as target genes showing the highest number of integration with miRNAs. Genes such as PEG10 [Li et al. 2016] and PTP4A1 [Hu et al. 2016] were associated with invasion of hormonal-receptor positive breast cancer cells, but these genes may be involved in invasion of TNBC cells.

In target gene - TF network, up regulated genes were identified as target genes showing the highest number of integration with TFs. HOPX was responsible for pathogensis of TNBC [Tanaka et al. 2019]. ACTA2 was involved in invasion of hormonal-receptor positive breast cancer cells [Jeon et al. 2016], but this gene may be linked with invasion of TNBC cells. SNP in CYP1B1 was liable for advancement of hormonal-receptor positive breast cancer [Ahsan et al. 2004], but this poly morphic gene may be responsible for development of TNBC. Similarly, down regulated genes were identified as target genes showing the highest number of integration with TFs. Methylation inactivation of tumorsupressor NR0B2 was associated witrh peogression of renal cancer [Kudryavtseva et al. 2018], but loss of this gene may be important for pathogensis of TNBC. CCNB2 was linked with progression of TNBC [Shubbar et al. 2013].

Identified ten hub genes from PPI network were validated by survival analysis, expression analysis, stage analysis, mutation analysis, IHC analysis, ROC analysis, RT-PCR and immune infiltration analysis. Future studies with in-depth verification of more candidate genes.

In conclusion, a total of 949 DEGs are identified by GEO profile, and ADAM15, BATF, NOTCH3, ITGAX, SDC1, RPL4, EEF1G, RPL3, RBMX and ABCC2 may be the key genes relevant to TNBC. Comprehensive bioinformatics analysis is a powerful tool for the biomarker screening. The interaction among selected proteins provides essential report on the development of TNBC, and molecular mechanism underlying the incidence, progression, metastasis and drug resistance of cancers. However, the predictive values of these biomarkers should be proved in more studies before their clinical application. The molecular mechanism elemental the activities of these genes is also required to be clear up, which is helpful for the picture of the predicted network of these genes. We optimize the parameters of integrated bioinformatics analysis to screen the ability biomarkers for the initial TNBC diagnosis, prognosis and thraputic targates, and we prospect our study implements novel approaches for the inital diagnosis, prognosis and thraputic targates of TNBC.

## Acknowledgement

I thank Benjamin Haibe-Kains, Princess Margaret Cancer Centre, Princess Margaret Research, Bioinformatics and Computational Genomics, 610 University Avenue, Toronto, Ontario, Canada, very much, the author who deposited their microarray dataset, GSE88715, into the public GEO database.

## Ethical approval

This article does not contain any studies with human participants or animals performed by any of the authors.

## Informed consent

No informed consent because this study does not contain human or animals participants.

## Availability of data and materials

The datasets supporting the conclusions of this article are available in the GEO (Gene Expression Omnibus) (https://www.ncbi.nlm.nih.gov/geo/) repository. [(GSE88715) (https://www.ncbi.nlm.nih.gov/geo/query/acc.cgi?acc=GSE88715)]

## Consent for publication

Not applicable.

## Author Contributions

B. V. - Writing original draft, and review and editing

C. V. - Software and investigation

I. K. - Supervision and resources

